# Evaluating anaesthetics for improving scientific research and welfare using larval zebrafish

**DOI:** 10.1101/2025.10.06.680636

**Authors:** Sylvia Dimitriadou, Mhairi Miller, Ali Pilehvar, Lynne U. Sneddon, Jessica L. Bamsey, Damilola Hogan-Bassey, Maciej Trznadel, Jonathan S. Ball, Aya Takesono, Courtney Hillman, Hannah Rickard, Grace Popplewell, Jenna Corcoran, Tetsuhiro Kudoh, Charles R. Tyler, Matthew J. Winter

**Affiliations:** Biosciences, Faculty of Health and Life Sciences, University of Exeter, Exeter, Devon EX4 4QD, UK; Department of Biological and Environmental Sciences, University of Gothenburg, Gothenburg, Sweden; Living Systems Institute, University of Exeter, Exeter, Devon EX4 4QD, UK; Surrey Sleep Research Centre, Department of Clinical and Experimental Medicine, School of Biosciences, University of Surrey, Guildford, Surrey, GU2 7XH, UK

## Abstract

Establishing anaesthesia for ensuring both animal welfare and compatibility with protocols required for different areas of scientific research is vital. Zebrafish (*Danio rerio*) are one of the most used animal models in research; however, little is known about the appropriateness of anaesthetic used for this species, especially for embryo-larval life stages. Using a combination of whole-brain functional imaging, quantification of cardiovascular performance, and behaviour, we explore the efficacy and tolerability of six widely used fish anaesthetics (2-phenoxyethanol, benzocaine, etomidate, MS222, isoeugenol and quinaldine sulfate) in larval zebrafish. We show that MS222 and quinaldine sulfate are the most suitable for achieving deep anaesthesia, while etomidate is better suited for studies focused on the cardiovascular system. Only quinaldine sulfate was found to be aversive. Our findings aid researchers for selecting the most suitable anaesthetic compounds and concentrations for their specific research goals, and the refinement of studies using anaesthesia in larval zebrafish.

## Introduction

Zebrafish (*Danio rerio*) are one of the most widely used laboratory animals for scientific research, with global usage estimated to be in the millions of animals per year^1^. In the European Union, in 2022, zebrafish made up 7.1% of all scientific animal procedures^2^, while in the UK (in 2023) this figure was double this level at 14%, with 308,906 experimental procedures recorded^3^. This popularity of the zebrafish is due to several key attributes, including its optical clarity in the embryo-larval form, the ease of its genetic manipulation, and availability of extensive genomic resources. Within the UK and European Union, zebrafish at pre-independent feeding life stages (animals less than 5 days post fertilisation (dpf)) are not considered as protected animals, avoiding legislative restrictions and favouring them further as an experimental model. For these reasons, the zebrafish is now the *de facto* lower vertebrate model of choice across a wide range of biomedical research, including developmental biology, ecotoxicology, and human drug discovery^4,5^. Most zebrafish used in research will undergo anaesthesia at least once as part of routine husbandry (e.g. fin clipping or tagging), for sedation to facilitate experimental manipulation, and/or as the first step towards euthanasia^1^. Despite this, little data are available on the effectiveness or tolerability of anaesthetics in zebrafish^1,6,7^, especially for embryonic and larval forms, which constitute the most frequently used zebrafish life stage in research.

General anaesthesia is defined as a reversible, central loss of sensory perception, accompanied by a sleep-like state, that is usually induced by drugs or anaesthetics^8^. The level (depth) of anaesthesia^10,11^ depends on anaesthetic concentration and exposure duration^11^. Preferred anaesthetics possess several attributes associated with their effectiveness, tolerability, and ease of use. In fish, anaesthesia should induce sedation within 3 minutes with complete recovery from a 10-minute exposure within 15 minutes, without mortalities^12,13^. They should also have low toxicity and good tolerability^10,14^ and, given the typical route for anaesthesia is via immersion^15^, sufficient aqueous solubility^16,17^. Fish anaesthetics should also provide humane induction without causing undue stress or long-lasting harm^10,14^. The most widely used anaesthetic for fish, is tricaine methansulfonate (MS222)^9,18^. Recent studies, however, have suggested this anaesthetic possesses some unfavourable attributes including induction of avoidance (or aversion) behaviours^19–21^, hypoxemia^22^, and bradycardia that can lead to accidental overdose and/or death^23^. These effects have been demonstrated in adult zebrafish^9,22,24^, but very little information is available for their use in embryonic or larval life stages.

Here we address the need for data-informed anaesthetic choices for experimentation with zebrafish embryo-larvae, with consideration for both scientific purpose and animal welfare. We assess the efficacy and tolerability of six of the most commonly used fish anaesthetics, namely 2-phenoxyethanol, benzocaine, etomidate, isoeugenol, MS222, and quinaldine sulfate using multiple indicators of effectiveness and tolerability (see figure 1). Effectiveness was determined using a combination of whole brain functional imaging and video tracking-based assessments of behaviours capturing locomotion and responsiveness to external stimulation. Tolerability was quantified through behavioural responses including aversion, general locomotion and anxiety (thigmotaxis), along with cardiovascular function as a measure of altered physiological health^25^. Bioavailability of these agents was quantified using liquid chromatography with tandem mass spectrometry (LC-MS/MS) on whole body extracts. We show that the anaesthetic agents tested differed in their ability to induce surgical level (deep) anaesthesia, with differing effects on behavioural and on cardiovascular indicators of tolerability. The data presented enables researchers to select the most appropriate anaesthetic for procedures in larval zebrafish, with a particular emphasis on compounds and concentrations suitable for use in studies where wider central nervous system (CNS) and cardiovascular functional impairment need to be avoided.

**Figure 1.**
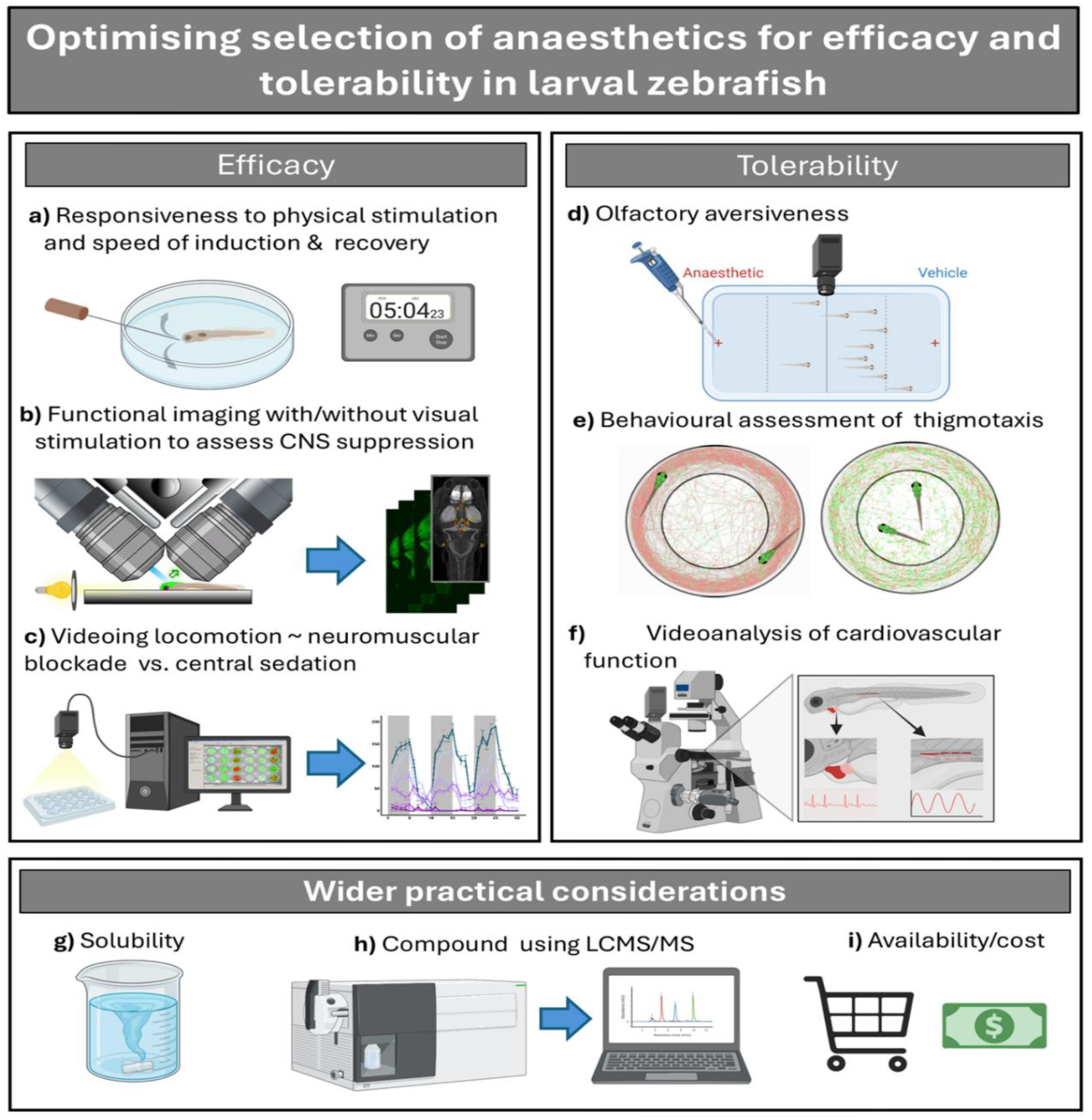
Schematic showing the experimental approaches used to assess the efficacy and tolerability of several commonly used anaesthetics in larval zebrafish. Techniques used for testing efficacy were a) assessments of touch responsiveness and balance maintenance; b) functional brain imaging with or without provision of light stimulation; c) video tracking of larval locomotion, stimulated using a light-dark cycle. Tolerability was assessed using d) assessment of aversion to each agent; e) behavioural assessment of thigmotaxis or ‘wall-hugging indicating anxiety’ and ; f) impacts on heart and vascular functional parameters using video analysis. Other considerations for a ‘good’ fish anaesthetic included g) aqueous solubility; h) compound body tissue uptake measured using mass spectrometry; i) and availability and cost. Figure prepared using BioRender.com.

## Results

### Behavioural measures of effectiveness

Using a stimulus responsiveness test, a standard approach for assessing anaesthetic depth in laboratory fish^9,10,18^, the maximum non-lethal concentration (where animals recovered after removal from the anaesthetic solution), and the lowest effective anaesthetic concentration were defined behaviourally for each of the six anaesthetics (Table 1). Resultant safety margins, defined as the difference between these two values, indicate that 2-phenoxyethanol has the largest safety margin, followed by MS222 and quinaldine sulfate.

**Table 1.**
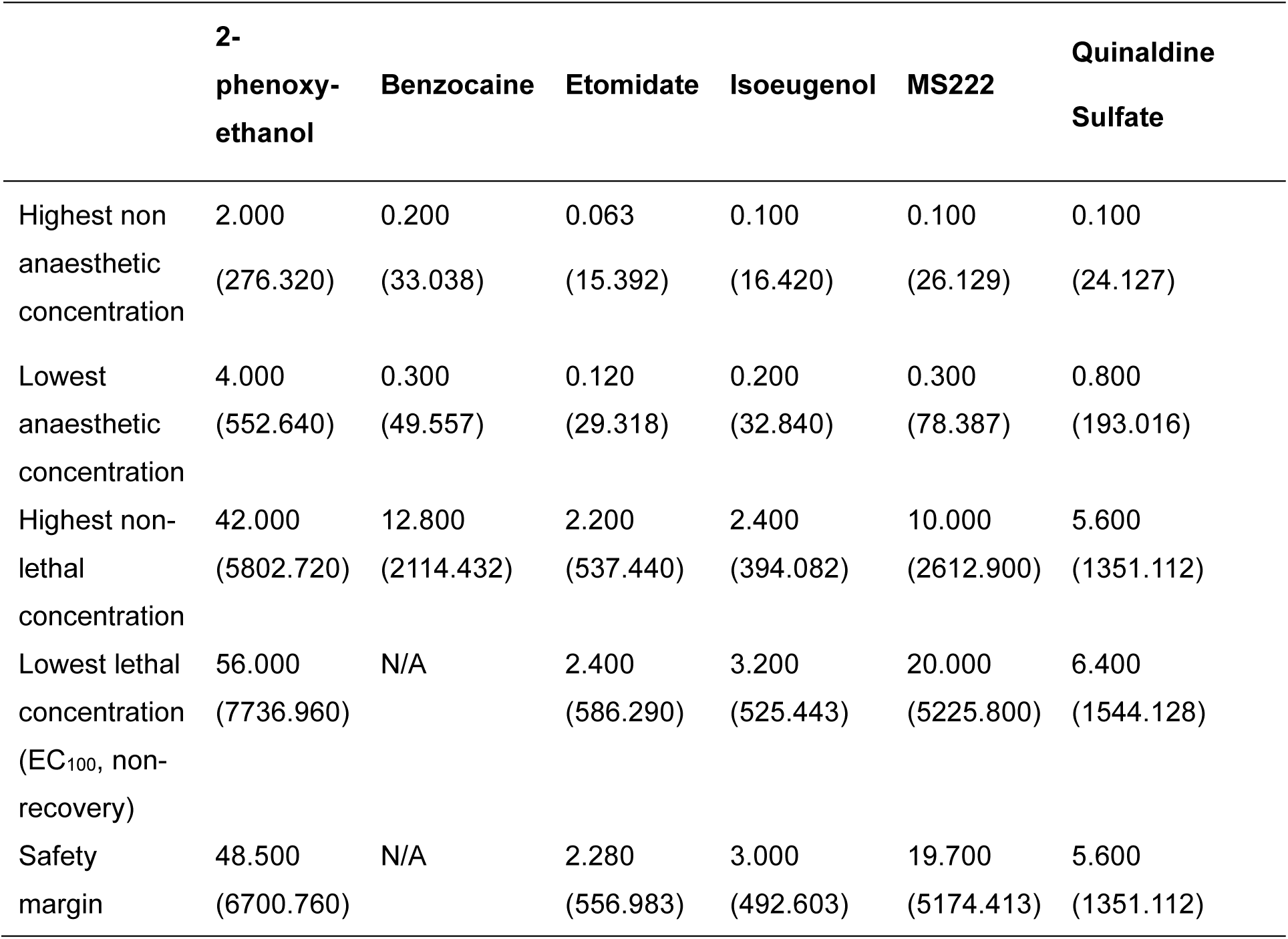
Summary of effective and lethal concentrations of the six anaesthetics tested, in mM (and mg/L). For this assessment, full anaesthesia was defined as the concentration resulting in unresponsiveness to tactile stimulation within 15 minutes of exposure, while non-recovery was determined to by the absence of a heartbeat and unresponsiveness to the same stimulus after 15 minutes of recovery in solution with no anaesthetic. The safety margins shown here are defined as the difference between the maximum non-lethal concentration and the lowest effective concentration. We could not reach a lethal concentration of benzocaine within 15 minutes of induction as the highest tested concentration constituted the limit of solubility for this compound.

### Anaesthetic bioavailability

Bioavailability of tested compounds was assessed by measuring internal concentration by LC-MS/MS after 15-minute exposure. With the exception of 2-phenoxyethanol, which was unmeasurable due to lack of formation of stable parent and fragment ions, all anaesthetics were found in larvae at levels ranging from between 20% to 750% of the external exposure concentration (Table 2). The lowest level of uptake, which was around 35% of the external exposure concentration, was for benzocaine and MS222, which are chemically very similar. For etomidate and quinaldine sulfate, uptake levels were around 140% of the external exposure concentration. There was no clear concentration-dependent uptake for these four compounds. In the case of isoeugenol, there was a slightly higher uptake into larvae exposed to the lowest external concentration, compared to the two higher exposure concentrations.

**Table 2.** Bioavailability of the five anaesthetics used in this study (excluding 2-phenoxyethanol). The internal concentrations were calculated on the basis of the volume of 4.5 dpf embryos at 0.44 µL. Uptake was calculated as ratio of internal and external concentration. CV is the coefficient of variation across three replicate samples.

| Compound | Conc. Tested<br>[mM] | Internal Conc.<br>[mM] | CV<br>[%] | Uptake<br>[%] |
| --- | --- | --- | --- | --- |
| Benzocaine | 1.9 | 0.45 | 9.6 | 24 |
|  | 3.8 | 1.1 | 26 | 28 |
|  | 7.7 | 3.7 | 21 | 49 |
| Etomidate | 0.38 | 0.56 | 17 | 149 |
|  | 0.75 | 1.3 | 13 | 168 |
|  | 1.5 | 1.3 | 21 | 86 |
| Isoeugenol | 0.24 | 1.8 | 5.4 | 751 |
|  | 0.48 | 2.8 | 11 | 576 |
|  | 0.96 | 4.6 | 8 | 477 |
| MS222 | 0.6 | 0.15 | 3.7 | 24 |
|  | 1.2 | 0.52 | 19 | 44 |
|  | 2.4 | 0.94 | 18 | 39 |
| Quinaldine sulfate | 0.24 | 0.37 | 9.3 | 152 |
|  | 0.48 | 0.57 | 5.1 | 118 |
|  | 0.96 | 1.2 | 13 | 128 |

### Functional brain imaging

Functional brain imaging was used to systematically compare anaesthetic effectiveness across concentration ranges observed to impact stimulus responsiveness during initial efficacy assessments (i.e. concentrations greater than the highest non-anaesthetic concentration; Table 1). This approach provides information on CNS activity levels in different brain regions before and after anaesthetic administration, without relying on behavioural responsiveness, which could be affected by neuromuscular blockade without central sensory suppression.

The different anaesthetic treatments generally resulted in widespread and concentration-dependent reductions in neural activity across large areas of the larval zebrafish brain (as indicated by GCaMP mean fluorescence intensity), supporting a general reduction in neural activity with deepening anaesthesia. In the case of 2-phenoxyethanol, brain activity was significantly lower, when compared to the controls, across 86% of the brain regions assessed in larvae exposed to the two highest concentrations (5 mM and 7 mM; Figure 2, Table S1). Similarly, exposure to benzocaine at the highest concentration tested (0.80 mM) resulted in a reduced neural activity in 82% of the brain regions (Figure 2, Table S2), whereas exposure to 0.45 mM resulted in a reduction in activity in regions associated with motor control and the startle/escape response only. Exposure to etomidate led to a general reduction in activity in 47% of the brain regions assessed; however, while these regions showed an overall significant change in activity, no individual treatment group was significantly different from the controls following *post hoc* analysis (Figure 2, Table S3). The highest concentration of isoeugenol tested (0.40 mM) resulted in suppressed neural activity across 59% of all brain regions analysed, while exposure to 0.10 mM and 0.20 mM resulted in reduced activity only in regions associated with motor control, balance, signal perception and integration, as well as photoreception and sleep/wake cycles (Figure 2 Table S4). MS222 exposure resulted in reduced brain activity only at the highest concentration tested (1 mM), and only in around 9% of the brain regions assessed, which were associated with contextual fear, shoaling, and stress responses (Figure 2, Table S5). The highest quinaldine sulfate concentration tested (0.40 mM) resulted in lower activity in 11% of the brain regions tested, notably those associated with contextual fear and shoaling, stress responses, and the regulation of social behaviour. In contrast, at the lower concentration of quinaldine sulfate tested (0.20 mM), we found an increase in activity in 7% of brain regions studied, specifically in those associated with sensory input and motor control (Figure 2, Table S6).

**Figure 2.**
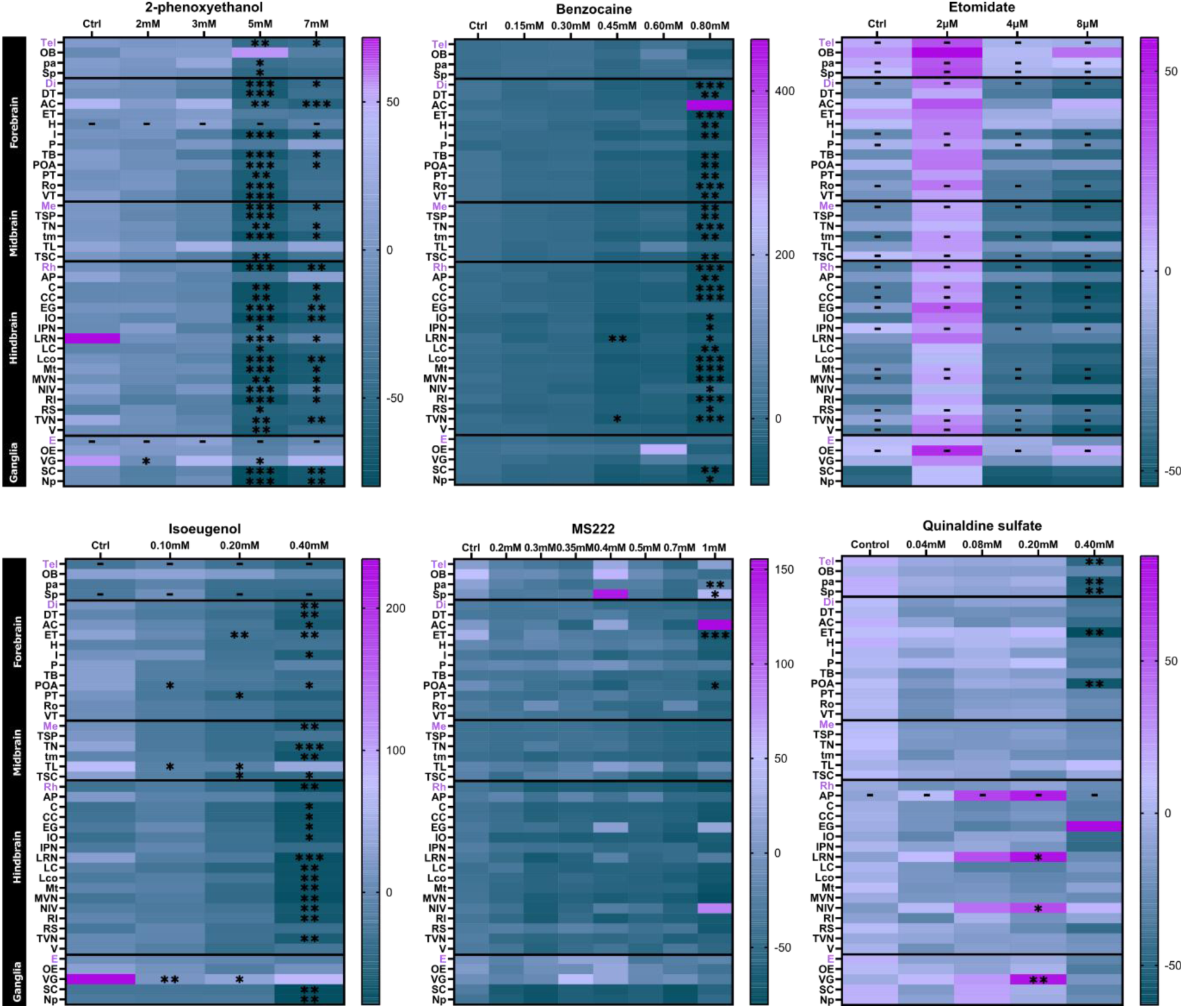
Heat maps illustrating neural activity (as indicated by mean GCaMP fluorescence intensity) in brain regions of interest (ROI) measured over after a 3-minute exposure to the tested anaesthetics. Data are presented as the **pre-exposure**-corrected mean fluorescence intensity averaged over the imaging duration, calculated using the equation: Δf/f = (f_1_ − f_0_)/f_0_ * 100 (where f_1_ = mean fluorescence intensity after anaesthetic exposure, and f_0_ = mean baseline fluorescence intensity, i.e. intensity pre-exposure, for the same animal within that ROI) (n = 7–9 larvae per group). Anaesthetic administration resulted in a lower level of brain activity generally, at the higher exposure concentrations, as expected. Statistically significant differences versus the corresponding control group are presented as *p<0.05, **p<0.01, and ***p<0.001. Cases where there was an overall difference in brain activity across treatments, but there were no individual differences (i.e. where specific concentrations tested did not differ in a statistically significant manner from the control condition) following *post hoc* tests (most notably etomidate), are represented by ‘–‘ in the figure. Abbreviations used for the brain regions assessed are: (Tel) telencephalon, (OB) olfactory bulb, (AC) anterior commissure, (pa) pallium, (Sb) subpallium, (Di) diencephalon, (ET) thalamic eminence, (H) habenuale, (I) intermediate hypothalamus, (P) pineal, (TB) posterior tuberculum, (POA) preoptic area, (PT) pretectum, (Ro) rostral hypothalamus, (VT) ventral thalamus, (Me) mesencephalon, (TSP) tectum stratum periventiculare, (TN) tectum neuropil, (tm) tegmentum, (TL) torus longitudinaris, (TSC), torus semicircularis, (Rh) rhombencephalon, (AP) area postrema, (C) cerebellum, (CC) corpus cerebelli, (EG) granular eminence, (IO) inferior olive, (IPN) interpeduncular nucleus, (LRN) lateral reticular nucleus, (LC) lobus caudalis cerebelli, (LCo) locus coeruleus, (Mt) Mauthner, (MVN) medial vestibular nucleus, (NIV) noradrenergic neurons of the interfascicular and vagal areas, (RI) inferior raphe nucleus, (RS) superior raphe nucleus, (TVN) tangential vestibular nucleus, (V) valvula cerebelli, (E) eyes, (OE) olfactory epithelium, (VG) vagal ganglia, (SC) spinal cord, (Np) neuropil region.

### In vivo imaging with visual stimulation

To more fully test the depth of anaesthesia, we further assessed brain function in Tg(*elavl3*:GCaMP6s) larval zebrafish under anaesthesia after application of a visual stimulus (light flash). The application of a light flash was designed to provide a direct CNS stimulus that would not be impaired through the non-central (e.g. neuromuscular) action of the tested agents, and an unequivocal indication of sensory unconsciousness. Overall, the brain of 4dpf larval zebrafish was relatively responsive to light stimulation, even at concentrations of anaesthetics where non-stimulus evoked activity was markedly reduced (Figure 3; see also Figure S1 for raw data). Most notably, in the case of isoeugenol, despite a clear concentration-dependent reduction in brain activity in the absence of an external stimulation, after stimulation, no difference between isoeugenol-exposed and control animals at any treatment level was observed (Figure 3, Table S10). Similarly, exposure to the highest concentration of 2-phenoxyethanol (28 mM) resulted in a significant decrease in activity only in 5% of the brain, namely the pallium (associated with contextual fear) and the thalamic eminence (a brain region only apparent in the developing, i.e. non-adult brain) (Figure 3 Table S7). Exposure to benzocaine also showed little impact on light-simulated brain activity, with the treatment of 0.60 mM affecting activity in only 7% of ROIs, and no significant changes at the highest concentration (0.80 mM) (Figure S1, Table S8).

**Figure 3.**
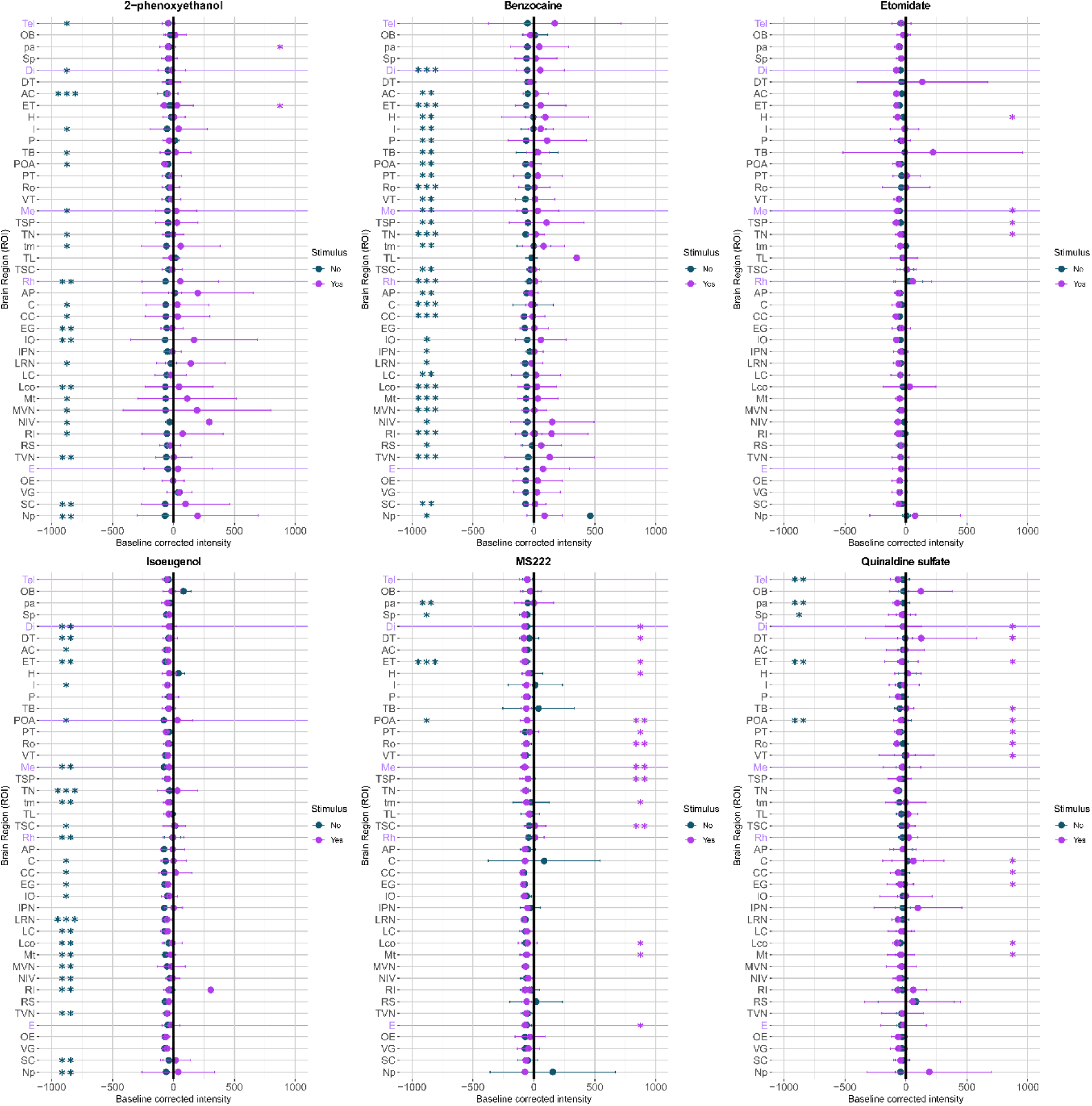
Brain activity (as measured by fluorescence intensity) in brain regions of interest (ROI) in embryo-larvae exposed to the tested anaesthetics, with (purple) and without (dark blue) the application of a light stimulus (only the highest concentrations tested in each experiment are shown in this plot). In most cases, administration of a light stimulus led to a less pronounced decrease in brain activity compared to the effects of the same anaesthetic without light stimulation, suggesting persistent stimulatory responsiveness even in the presence of CNS suppression. Data are presented as the mean values, with error bars showing standard deviation. Statistically significant differences versus the control animals are presented as *p<0.05 and **p<0.01, with the colour corresponding to the related dataset. Abbreviations used for the brain regions assessed are: (Tel) telencephalon, (OB) olfactory bulb, (AC) anterior commissure, (pa) pallium, (Sb) subpallium, (Di) diencephalon, (ET) thalamic eminence, (H) habenuale, (I) intermediate hypothalamus, (P) pineal, (TB) posterior tuberculum, (POA) preoptic area, (PT) pretectum, (Ro) rostral hypothalamus, (VT) ventral thalamus, (Me) mesencephalon, (TSP) tectum stratum periventiculare, (TN) tectum neuropil, (tm) tegmentum, (TL) torus longitudinaris, (TSC), torus semicircularis, (Rh) rhombencephalon, (AP) area postrema, (C) cerebellum, (CC) corpus cerebelli, (EG) granular eminence, (IO) inferior olive, (IPN) interpeduncular nucleus, (LRN) lateral reticular nucleus, (LC) lobus caudalis cerebelli, (LCo) locus coeruleus, (Mt) Mauthner, (MVN) medial vestibular nucleus, (NIV) noradrenergic neurons of the interfascicular and vagal areas, (RI) inferior raphe nucleus, (RS) superior raphe nucleus, (TVN) tangential vestibular nucleus, (V) valvula cerebelli, (E) eyes, (OE) olfactory epithelium, (VG) vagal ganglia, (SC) spinal cord, (Np) neuropil region.

In contrast with the aforementioned compounds, for etomidate there was a pronounced overall impact on light-stimulated activity, with several brain regions, including those associated with contextual fear, homeostasis, stress regulation, and those associated with the processes underlying brain development in early life showing lower neural activity. While, in these instances, we found a significant overall effect of etomidate administration on activity of these ROIs, subsequent *post hoc* analysis did not reveal statistically significant differences between the different individual concentrations. This may be due to the relatively high variation in responsiveness between individuals. *Post hoc* analysis revealed, however, significant reductions in neural activity in only 4 (10%) of the ROIs and at the highest exposure concentration tested (600 µM) only. The affected regions were associated with vision and startle responses (notably, the tectum stratum periventiculare, tectum neuropil, and mesencephalon), and overall information integration (habenulae) (Figure 3, Table S9). These findings mirror those in the absence of light simulation, although in the former the effect of etomidate exposure appeared to be more widespread.

For MS222 and quinaldine sulfate, responses after anaesthetic exposure followed by light stimulation were similar to those obtained in the absence of light stimulation (Figure 3). In MS222-treated animals, after light stimulation, there were significant reductions in neural activity in 39% and 34% of brain regions after exposure to 0.7 mM and 1.0 mM, respectively. These included ROIs associated with motor control and learning, signal perception and integration, photoreception, homeostasis and startle response (Figure S1, Table S11). The highest concentration of quinaldine sulfate tested (0.8 mM) also led to a widespread decrease in light stimulus-induced activity brain activity (33% of the ROIs analysed; Figure 3, Table S12).

### Behavioural measures of tolerability

A key consideration when selecting anaesthetics for use in fish is that the anaesthetic substance is well tolerated and is without properties associated with aversive or stress-related behaviours. To evaluate this, we assessed changes in anxiety (thigmotaxis) and olfactory-mediated aversion. Initial test concentrations were selected across the range comprising minimal to no induction of anaesthesia from the simple assessment of touch responsiveness shown earlier (Table 1). First, we quantified the concentration at which general locomotor activity was unaffected, to ensure any impacts on measures of stress/aversion were not merely due to reduced overall activity. As 4.5dpf larvae show low levels of spontaneous locomotion, a standardised light-dark stimulus protocol was used to elevate activity^26^, the results of which are presented in Supplementary Figure (Figure S2). Subsequent assessments of thigmotaxis and aversion were undertaken on non-inhibitory concentrations of each anaesthetic.

### Thigmotaxis

Assessment of the effects of anaesthetic treatment on thigmotaxis, as an indicator of stress/anxiety, are summarised in Figure 4. For all compounds, we observed a significant interaction between treatment and light phase on the proportion of time spent in the periphery of the well, indicating that, across compounds, the effect of anaesthetic administration was dependent on the light conditions (light vs. dark) (2-phenoxyethanol: χ^2^= 1041.955, p<0.001; benzocaine: χ^2^= 3023.529, p<0.001; etomidate: χ^2^= 101.811, p<0.001; isoeugenol: χ^2^= 6816.148, p<0.001; MS222: χ^2^= 1037.671, p<0.001; quinaldine sulfate: χ^2^= 6764.623, p<0.001). Exposure to the highest concentration of benzocaine (3 mM) increased the thigmotactic response in the dark but not in the light phase (z= 3.284, p=0.005), supporting an anxiogenic effect. At the highest concentration of MS222 not impeding locomotion (0.2 mM), thigmotaxis was increased in the light phase, but not the dark phase, compared with the control (dark: z=5.513, p<0.001, light: z=4.796, p<0.001), suggesting anxiogenesis. Quinaldine sulfate at non-inhibitory concentrations resulted in increased thigmotaxis under both light and dark conditions (0.04 mM, light: z=2.436, p=0.044; 0.08 mM, dark: z=2.627, p=0.026, light: z=3.900, p<0.001; 0.20 mM, dark: z=13.512, p<0.001, light: z=8.338, p<0.001). Etomidate and isoeugenol exposure had similar effects, increasing thigmotaxis in both light and dark conditions, but only at the mid-range concentrations tested (etomidate: 0.0075 mM, dark: z=3.394, p=0.003, light: z=3.447, p=0.002; 0.015 mM, light: z=7.509, p<0.001; isoeugenol: 0.25 mM, dark: z=3.572, p=0.002, light: z=3.513, p=0.002; 0.50 mM, dark: z=6.089, p<0.001, light: z=5.667, p<0.001). We found no effect of 2-phenoxyethanol on thigmotaxis under any of the treatment or lighting conditions tested, suggesting that this compound has no strong anxiogenic or anxiolytic effects.

**Figure 4.**
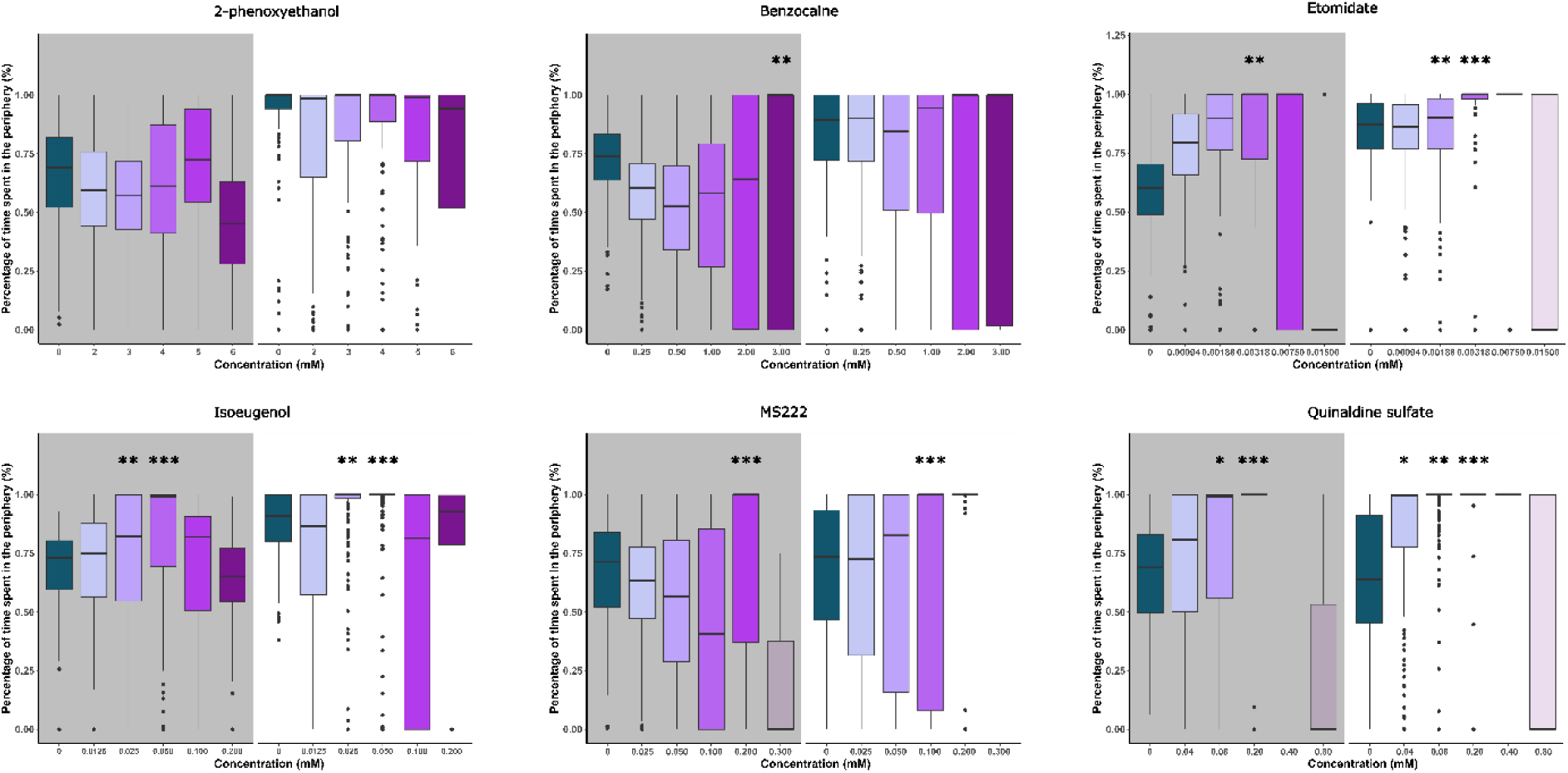
Thigmotaxis, as the percentage of time spent in the periphery of the test arena, during dark (grey shaded areas) and light (white areas) phases. Boxes represent the interquartile range (25th and 75th quartiles), and the horizontal lines represent the medians. The upper whiskers represent the largest value within 1.5 times the interquartile range (1.5*IQR), while the lower whiskers represent the lowest within 1.5*IQR (Tukey boxplot). Transparent boxplots (such as those on 0.300 mM for MS222) denote conditions that were excluded from the analysis as there was very little overall movement. Statistically significant differences compared to the controls are represented by *p< 0.05, **p<0.01, and ***p<0.001.

For all compounds there was a positive relationship between the z-score of the distance travelled and thigmotaxis, meaning that individuals that moved more did so around the edges of the well (2-phenoxyethanol: χ^2^=4806.491, p<0.001; benzocaine: χ^2^=24726.638, p<0.001; etomidate: χ^2^=0.483, p<0.001; isoeugenol: χ^2^=28772.687, p<0.001; MS222: χ^2^=60070.942, p<0.001; quinaldine sulfate: χ^2^=27047.124, p<0.001), thus demonstrating an inherent propensity for wall hugging.

### Aversion

The data from the assessment of aversion are summarised in Figure 5 with additional data provided in the Supplementary Information (Table S15). Both treatment (χ^2^ =41.1965, p<0.001) and the side of the test chamber that the substance was administered (χ^2^ =11.4765, p<0.001) (controlling for any side bias) affected the overall position of the larvae within the arena. When the effects of the different treatments were analysed, individuals were found to avoid the site of administration of cadaverine (our positive control) (t=2.945, p=0.004), as expected. The only anaesthetic to show a similar aversive effect in 4.5dpf zebrafish larvae was quinaldine sulfate (0.1 mM) (t=3.883, p<0.001).

**Figure 5.**
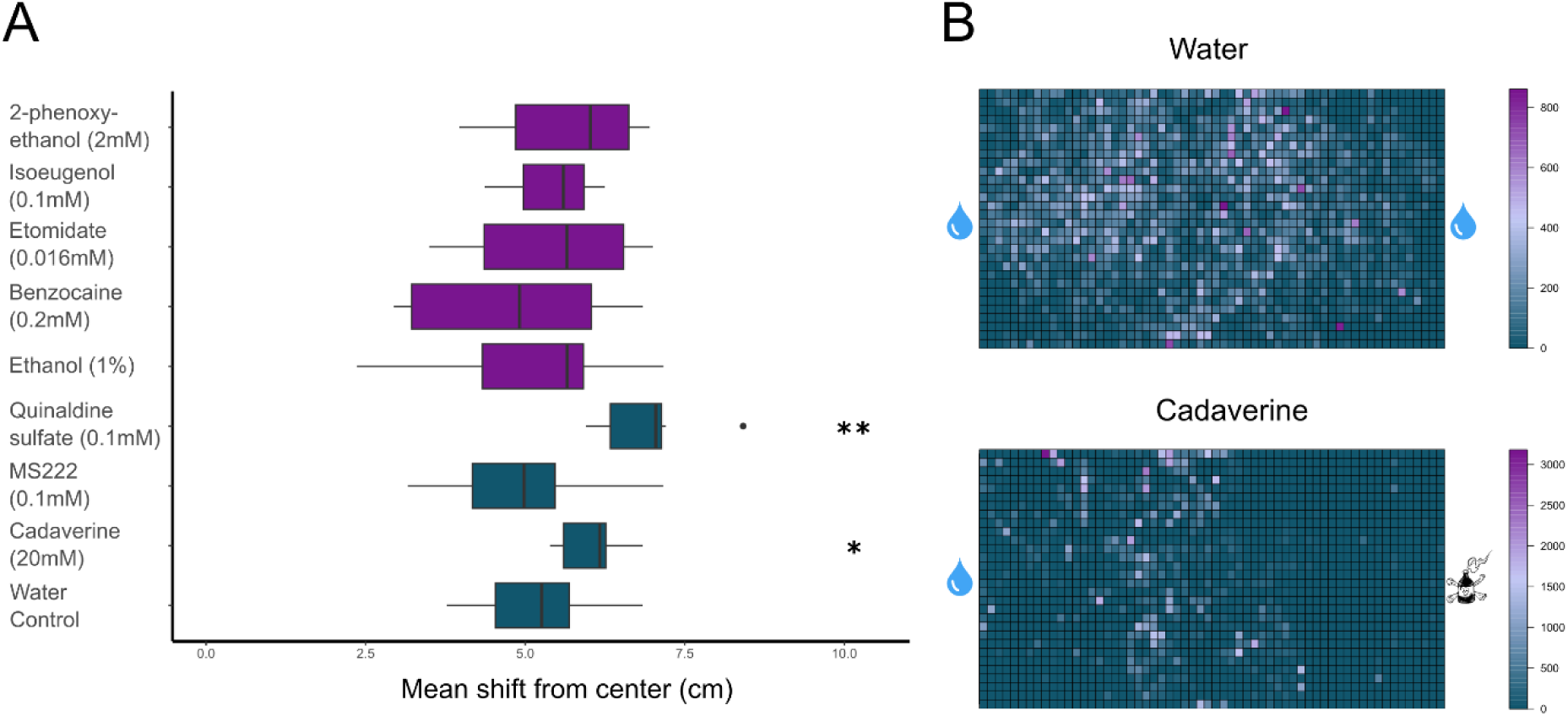
**Panel A. Effects of anaesthetic administration on the average distance moved by groups of embryo larvae from the point of compound administration.** Purple denotes a comparison to a 1% ethanol solvent control (ethanol-soluble compounds), and dark green a comparison to water control (water-soluble compounds). The x-axis represents the distance (in cm) from the site of anaesthetic administration, with higher values indicating greater avoidance. Boxes represent the interquartile range (25th and 75th quartiles), and the vertical lines represent the medians. The right whisker extends to the largest value no more than 1.5 times the interquartile range (1.5*IQR), while the left whisker extends to the smallest value within 1.5 times the interquartile range (Tukey boxplot). Statistically significant difference compared to control *: p< 0.05; **: p<0.01. **Panel B. Example heatmaps of the position occupied by the groups of larvae when water only was administered on both sides of the experimental arena (top), and when cadaverine (positive control) was administered to the right side, while water was administered on the left side (bottom).** Larvae tended to occupy positions further away from the side of administration of aversive compounds, such as cadaverine.

### Cardiovascular function

Cardiovascular function was assessed as a key indicator of physiological status and for which larval zebrafish are widely utilised. Results from this assessment are summarised in Figure 6, with additional data in the Supplementary Information (Figure S2). Both atrial (ABR) and ventricular (VBR) beat rates were reduced in concentration-dependent manners after treatment with all anaesthetics (2-phenoxyethanol ABR: χ^2^ = 24.31, p<0.0001 and VBR: χ^2^ =23.06, p<0.0001; benzocaine ABR: χ^2^ =23.8, p<0.0001 and VBR: χ^2^ =23.64, p<0.0001; etomidate ABR: χ^2^ =29.96, p<0.001 and VBR: χ^2^ =32.76, p<0.001; isoeugenol ABR: χ^2^ =23.95, p<0.0001 and VBR: χ^2^ =23.71, p<0.0001; MS222 ABR: F =14.31, p=0.0001 and VBR: F_(2)_=15.33, p<0.0001; quinaldine sulfate ABR: F_(3)_=80.88, p<0.0001and VBR: F_(3)_=21.69, p<0.0001). In all cases, the reduction observed was statistically significant at either the highest, or two highest, treatment concentrations (Figure 6). For all anaesthetics, except for etomidate, there was also a concentration-dependent reduction in blood flow and blood linear velocity, supported statistically at the highest, or two highest, treatment concentrations (Figure S2). Surrogate stroke volume (SSV) and cardiac output (SCO) data (Figure S3) broadly followed the observed concentration dependent reductions in heart rate and blood flow. Interestingly, in the case of benzocaine and quinaldine there was no significant impact on SSV over the treatment range despite SCO being affected, suggesting perhaps some variability between individual cardiac cycles that are averaged to calculate SCO. Etomidate had no effect on either SSV or SCO (Figure S3).

**Figure 6.**
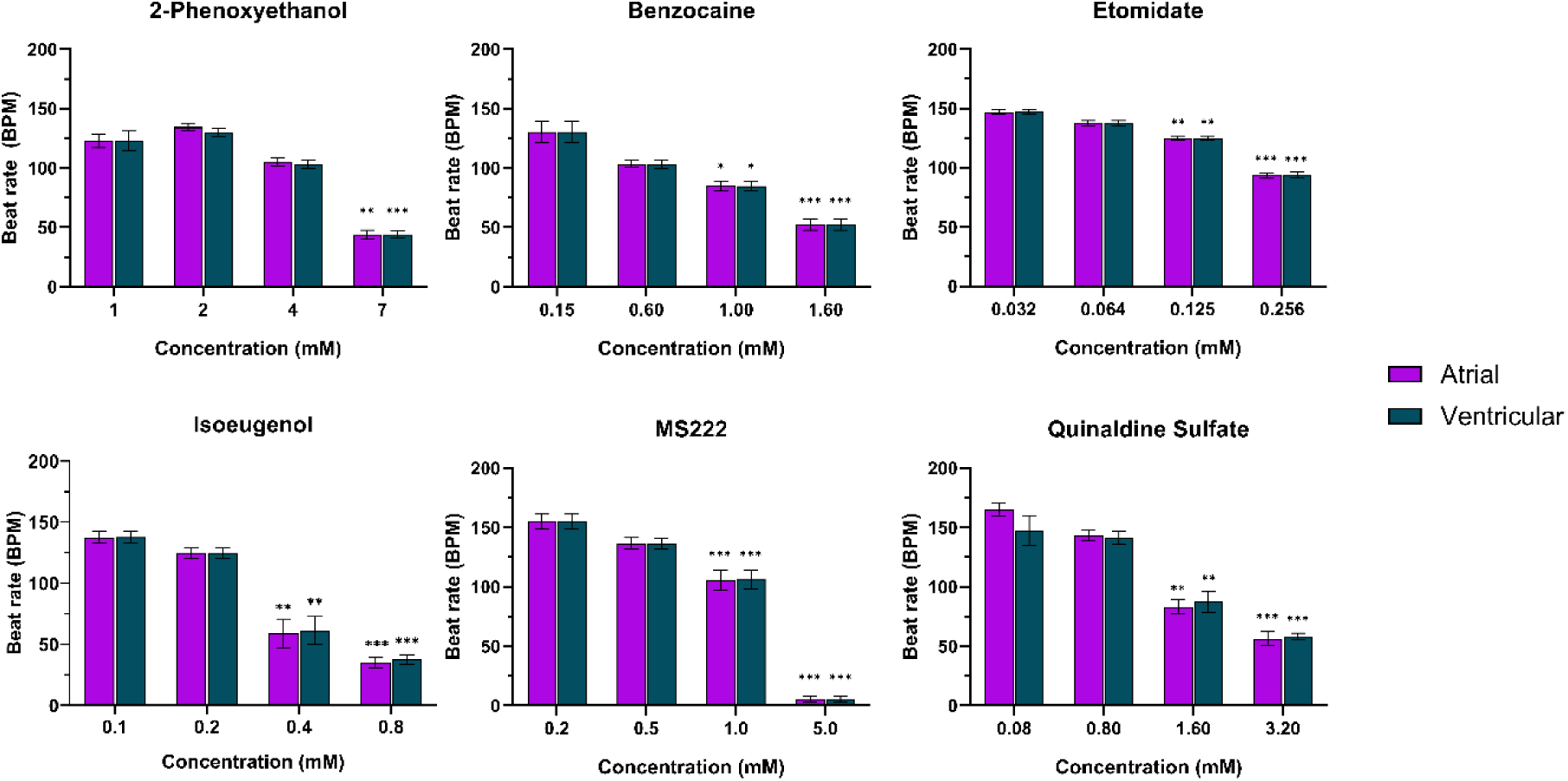
Effects of the six test anaesthetics on atrial and ventricular beat rates in 4dpf zebrafish. Data are presented as mean ± SEM (n=8 for each treatment concentration). Statistically significant differences compared to control (lowest concentration for each compound) are presented as *p< 0.05, **p<0.01 and ***p<0.001.

## Discussion

Here we determine the most effective and tolerable general anaesthetics for use in larval zebrafish, addressing a fundamental knowledge gap in our understanding of anaesthesia in the most widely used laboratory fish species. Employing a range of behavioural, imaging, physiological, and bioanalytical techniques, we establish effective concentrations for inducing sedation versus immobilisation, as well as assessing indicators of stress, aversion, and adversely impacted cardiovascular function. Our findings support researchers in making informed choices regarding the anaesthetic agents most suited to their experimental requirements, while also considering animal welfare.

All six tested anaesthetic compounds (2-phenoxyethanol, benzocaine, etomidate, isoeugenol, MS222, and quinaldine sulfate) induced anaesthesia, defined as a loss of responsiveness to tactile stimulation, after 15 minutes of exposure, and all compounds, except for benzocaine, were also lethal at high concentrations. Safety margins varied considerably between compounds, with 2-phenoxyethanol demonstrating the highest level (48.5 mM) and etomidate the lowest (2.28 mM). The inability to find a lethal concentration for benzocaine, tested up to 12.8 mM, is supported by previous work demonstrating that even after a 1-hour long exposure to 6 mM benzocaine, 10% of the exposed 4dpf zebrafish embryos survived^27^. Our data, therefore, suggest that agents other than benzocaine may be more appropriate for larval zebrafish euthanasia.

Using functional brain imaging, we provide the first detailed analysis of the comparative impact on brain function for these six anaesthetics. *In vivo* Ca^2+^ imaging indicated that exposure to high concentrations of 2-phenoxyethanol, benzocaine, and isoeugenol significantly reduced neuronal activity across most brain regions. We cannot set the threshold for the level of neuronal activity representing (un)consciousness^28^; however, the degree of brain activity depression observed can be considered a reliable indicator of some form of loss of consciousness at these concentrations. We found that, for MS222 (1 mM) and quinaldine sulfate (0.4 mM), decreased neural activity occurred only in a small number of brain regions, including the pallium, subpallium, thalamic eminence and the preoptic area, as well as the telencephalon in the case of quinaldine sulfate. It is important to note that all of these regions are in the forebrain, which is associated with cognitive functions in zebrafish, including aspects of consciousness^28–31^. There was no evidence for a widespread suppression of brain activity after exposure to etomidate up to 16 μM. This is in contrast with a recent study demonstrating a suppression of brain activity in 7dpf zebrafish embryos, as measured via confocal microscopy after acute exposure to 12.3 μM of etomidate^32^. This discrepancy may be due to methodological differences, such as the length of exposure, or the use of slightly different larval ages (e.g. 7 versus 4 dpf). Interestingly, metomidate, a methylated derivative of etomidate, believed to have the same psychoactive effect, has also been reported as unsuitable for inducing surgical anaesthesia in fish, based on its effects on opercular respiration and muscle fasciculations indicative of incomplete relaxation^see^ ^15,33,34^.

The presentation of a light stimulus altered patterns of larval brain activity substantially. Stimulation was introduced to further assess the depth of anaesthesia, and to differentiate a central loss of stimulatory responsiveness from peripheral actions on neuromuscular junctions, i.e. stopping the perception of stimuli peripherally, and not in the CNS. Although effects vary across anaesthetic agents, monitoring of visually evoked neural activity is a valuable tool for the assessment of anaesthetic depth based upon central stimulatory suppression^35^. In mammals at moderate levels of anaesthesia, visual centres can still respond to light stimulation^36^: Liu *et al*.^37^ found that the application of monochromatic blue light to mice anesthetised with 2.5% sevoflurane resulted in electroencephalogram-measured increases in cortical and suprachiasmatic nucleus activity. Here, in the case of 2-phenoxyethanol, concentrations found to reduce brain activity in the absence of stimulation were no longer effective; similarly, there was no clear suppressive effect of benzocaine or isoeugenol exposure on brain activity after visual stimulation. In fact, only MS222 and quinaldine sulfate induced significant reductions in brain activity across multiple regions after light stimulation. In the case of etomidate (at 600 μM), the effects of exposure after light stimulation differed to those observed under non-stimulated conditions, with significantly lower neural activity in just 4 ROIs: the habenula, a region important for information integration^38^, the mesencephalon as a whole, the tectum stratum periventiculare, and tectum neuropil, which are all associated with visual stimulus processing^34,35^. This suggests that while a high concentration of etomidate does not induce unconsciousness, it may affect aspects of vision/visual processing.

An emergent theme across all six anaesthetics tested is that, at concentrations causing a loss of responsiveness to tactile stimulation, there is limited suppression of brain activity. This addresses a central question about the appropriateness for the use of some fish anaesthetics, as they may simply be acting principally as neuromuscular blocking agents or local anaesthetics, rather than central nervous system depressants^40^. Such an example is MS222, where at 0.1 mM there was virtually a complete cessation of locomotor activity, but neural activity was not significantly affected until below 0.7 mM. This suggests that MS222 acts initially as a neuromuscular blocker at concentrations not sufficient to significantly reduce CNS activity. Our results agree with the findings of Ohnesorge and colleagues^28^, who used confocal microscopy on 4-5dpf *elavl3*:H2B-GCaMP6s zebrafish embryo-larvae to show decreased neuronal activity in some brain regions after a 3-minute exposure to 0.64 mM of MS222. Similar findings have also been reported for 5-7dpf embryo larvae measured by Ca^2+^ two-photon imaging^41^. While very few other studies have explored the effects of any anaesthetic directly on brain activity, Machnik and *et al*. used electrophysiological recording from Mauthner cells to show that 0.365 mM isoeugenol also principally acts as a local anaesthetic and blocks sensory input to the CNS of goldfish (*Carassius auratus*), without effectively inducing general anaesthesia^40^. Our data support this observation as, in the presence of a visual stimulus, exposure of up to 2.4 mM isoeugenol resulted in no changes in the activity of any brain region.

A key consideration when choosing an appropriate anaesthetic agent is the delivery of humane induction with minimal stress^14^. Consequently, assessment of indices such as anxiety or aversion can provide important, and easily quantifiable, indicators of tolerability^42^. Thigmotaxis is a simple measure of exploratory behaviour, an increase in which can indicate anxiogenesis^42^. Exposure to benzocaine, etomidate, isoeugenol, MS222, and quinaldine sulfate increased thigmotaxis, especially under dark conditions, suggesting these agents may be anxiogenic at sub-anaesthetic concentrations. Given that 2-phenoxyethanol exposure did not increase thigmotaxis, this compound is particularly well tolerated by larval zebrafish. Interestingly, 2-phenoxyethanol has been found to be only mildly aversive in adult zebrafish compared to MS222 or benzocaine^19^ thus supporting our observations of higher tolerability. This, however, contrasts with previous studies in other fish species, where 2-phenoxyethanol has been reported to increase agitation in the Senegalese sole (*Solea senegalensis*) and the Pacific hagfish (*Eptatretus stoutii*)^43,44^. This discrepancy may be due to methodological or species-related differences, and highlights the importance, where possible, of testing the appropriateness of anaesthetics for a given study species.

Measurements of olfactory-mediated aversion suggested that quinaldine sulfate alone is aversive to zebrafish larvae, and only at the maximum non-anaesthetic exposure concentration of 0.016 mM. To our knowledge, the aversive properties of the various fish anaesthetics have not been studied in larval zebrafish; however, some of these compounds have been shown to be aversive to adult zebrafish, including MS222^19–21^, 2-phenoxyethanol, benzocaine, isoeugenol, and quinaldine sulfate^19^. The general lack of aversion associated with most anaesthetic agents tested in our study suggests good tolerability at this life stage. The discrepancy between our findings and those from studies on adult zebrafish may reflect differences in the olfactory systems between these life stages: zebrafish larvae between 3-6 dpf are only able to form rudimentary spatial odour maps^45^ and olfactory systems continue to mature until adulthood (>3-4 months post fertilization)^46,47^, which is likely to increase adult fish sensitivity to wider range of odorants compared with younger life stage animals.

Another crucial requirement for general anaesthesia is avoidance of adverse effects on cardiovascular function^48^, particularly as embryo-larval zebrafish are widely used for the study of cardiovascular development and disease^49,50^. We found that all compounds negatively impacted heart rate and blood flow (except etomidate) at concentrations that induce deep anaesthesia. In the only published study on the effect of an anaesthetic on zebrafish embryo cardiac function, MS222 was shown to suppress heart rate at concentrations between 0.8 and 1mM in zebrafish embryos^51,52^, and our findings support this, given the significant suppression observed at 1mM. This clear threshold above which cardiac function is impaired has important implications for the use of anaesthetics in the context of studying cardiovascular function. For instance, in the cases of MS222 and quinaldine, the only compounds for which we definitively identified suppression of light-induced sensory stimulation, the ranges of concentrations between the suppression of brain activity and impairment of cardiac function are narrow (between 0.7 – 1 mM for MS222, and 0.8 – 1 mM for quinaldine sulfate). Our findings, therefore, suggest that etomidate may be better suited for sedation (but not deep anaesthesia) during the assessment of CV function in larval zebrafish, especially given its lack of effect on blood flow-related parameters even at relatively high concentrations.

We may expect that the different effects of the six anaesthetics tested to be explained, at least in part, by differences in their respective modes of actions. For instance, benzocaine and MS222 are structurally and pharmacologically similar, both acting predominantly through voltage-sensitive Na^+^ channel blockade^53,54^. While MS222 administration significantly reduced neural activity across 39% of the brain regions analysed, benzocaine had a much less pronounced effect, decreasing brain activity in only 7%. The reason(s) for these differences in unclear, but may relate to differences in the penetration of these compounds into different brain regions – something that was not measured here. MS222 is known, due to its higher water solubility, to be absorbed rapidly^55^, and thus may result in higher concentrations at the site of action faster than benzocaine. While the mechanism of action of 2-phenoxyethanol has not been yet fully elucidated, it is suggested to act, at least partially, through NMDA-mediated modulation of ion channels^56,57^, ultimately suppressing neuronal activity in the higher regions of the CNS^58^. Benzocaine has also been shown to inhibit NMDA receptors *in vitro*^59^, and both compounds showed similar impacts on brain activity, but not locomotion and thigmotaxis. More work is needed to understand the mode of action of 2-phenoxyethanol and its effect on the endpoints measured here.

Both etomidate and isoeugenol are considered to act, at least partially, via modulation of γ-aminobutyric acid (GABA) signalling^58,60,61^. Activation of GABA receptors induces sedation and hypnosis^62^ but, as corroborated by our findings, limited anaesthesia and immobilisation. It should be noted, however, that in our study lower concentrations of etomidate (but not isoeugenol) showed evidence of a potential stimulatory effect, as indicated by a trend for increased neural activity in the absence of visual stimulation. It is, therefore, possible, that lower concentrations of some anaesthetics such as etomidate, which are insufficient to inhibit neuromuscular or CNS activity, have a mildly irritating effect.

Unlike the other compounds tested, the mechanism of action of quinaldine sulfate is completely unknown. Our findings suggest that the effects of this compound are phenotypically similar to those of MS222; however, quinaldine sulfate was the only anaesthetic we identified with aversive properties. Quinaldine sulfate, alongside MS222, benzocaine, 2-phenoxyethanol, isoeugenol, lidocaine hydrochloride, and propoxate, has been reported as aversive in adult zebrafish using a similar behavioural avoidance assessment method^19^, highlighting the importance of making appropriate anaesthetic choices not only based on species, but also life stage. The fact that quinaldine sulfate has been found to be particularly aversive in adult zebrafish^19^, as well as for larval zebrafish given our own findings, suggest that this compound specifically may be less suitable for use with zebrafish of any age.

## Conclusions

Our evaluation provides a comprehensive reference framework of empirical data to support researchers in their choice of anaesthetic agents most suited for their specific experimental requirements when using embryo-larval zebrafish. Collectively, our data reveal the comparative strengths and weaknesses of six anaesthetic agents commonly used in procedures employing larval zebrafish. Of the anaesthetics tested, 2-phenoxyethanol was identified as having the largest safety margin, followed by benzocaine and, as such, these compounds may be particularly suited for maintenance anaesthesia due to a lower risk of accidental overdose. Functional brain imaging revealed that MS222 and quinaldine sulfate are the most suitable for achieving deep anaesthesia, as the only agents to significantly and extensively decrease brain activity during visual stimulation, while etomidate may be more suited to studies on the cardiovascular system. In terms of tolerability, benzocaine, etomidate, MS222, quinaldine sulfate and isoeugenol appear to be anxiogenic in zebrafish larvae, although only quinaldine sulfate was found to be aversive. We conclude, therefore, that the most suitable anaesthetic depends on the context of the scientific procedure, and that none of the currently most widely used anaesthetics is universally applicable for studies with zebrafish larvae.

## Materials and Methods

Further details of the methods used are contained within the Supplementary Materials and Methods section.

### Compounds and concentration ranges

Six anaesthetics commonly used for anaesthesia in small freshwater fish species^9,15,19,21,33,63^ were selected for this study: 2-phenoxyethanol (2-PE, ethylene glycol monophenyl ether, Cas No. 122-99-6), benzocaine (4-aminobenzoic acid ethyl ester, Cas No. 94-09-7), etomidate ((*R*)-1-(α-Methylbenzyl)imidazole-5-carboxylic acid ethyl ester, Cas No. 33125-97-2), isoeugenol (2-Methoxy-4-propenylphenol, Cas No. 97-54-1), MS222 (tricaine methasulfonate, ethyl 3-aminobenzoate methanesulfonate, Cas No. 886-86-2), and quinaldine sulfate (2-methylquinoline, Cas No. 655-76-5). Quinaldine sulfate and MS222 were dissolved in zebrafish embryo-larvae culture water (see below), and 2-phenoxyethanol, benzocaine, etomidate, and isoeugenol were dissolved in 100% ethanol as a stock solution, which was then diluted in zebrafish embryo-larvae culture water to a final ethanol concentration of 1% for the working solutions. The pH for all solutions was (and where required adjusted to) around 7. The detailed description of methods for the determination of maximum tolerated concentration is provided in the Supplementary Materials and Methods and the final concentrations used for each experiment are summarised in Table S17.

### Experimental animals

All experiments were undertaken using 4.5 days post fertilisation (dpf) wild-type (WT), or 4 dpf Tg(*elavl3*:GCaMP6s) zebrafish larvae for brain activity measurements (*in vivo* Ca^2+^ imaging – supplied by Dr. Misha Ahrens, Janelia Research Campus, Howard Hughes Medical Institute, Ashburn, Virginia, USA). Adult zebrafish were held in aquaria at the University of Exeter, at 28°C ± 1°C under optimal conditions for spawning (14h:10h light:dark cycle, with 20 min dusk-dawn transition periods). Culture water consisted of dechlorinated mains tap water filtered by reverse osmosis (Environmental Water Systems UK Ltd.) and then reconstituted with Analar-grade mineral salts to a standard synthetic freshwater composition. Embryos were collected shortly after spawning (at lights on) and cultured in Petri Dishes under the same conditions as the adult fish used to supply the embryos. All work was undertaken under Project Licence (PP3242260) granted by the UK Home Office under the UK Animals (Scientific Procedures) Act 1986, and approved by the University of Exeter’s Animal Welfare and Ethical Review Body.

### Behavioural measures of effectiveness

Initial assessment of compound efficacy was undertaken as follows. Larvae were deposited individually into the wells of a 24 well plate across 5 concentrations of the treatment anaesthetic and a water or solvent control, and exposed for 15 minutes (n=8 per treatment). After 15 minutes had elapsed, larvae were observed under a microscope for loss of dorsoventral balance: this was assessed using a pipette tip to gently push the larvae onto its side and record the response, scored as 1 in the case of animals that righted themselves and 2 if they did not. Recovery was assessed by transferring the larvae back to freshwater after the 15-minute exposure period. Using this approach, we defined the maximum non-lethal concentration (where animals recovered after removal from the anaesthetic solution), and the lowest effective anaesthetic concentration, as well as the safety margin for each compound.

### Anaesthetic bioavailability

Bioanalysis of five of the six test compounds (the exception was for 2-phenoxyethanol) was performed on pooled samples of 4 embryos, run in triplicate for three test concentrations following the method outlined in Winter *et al*.^64^. These analyses were run for embryo-larvae exposed for 15 minutes, matching the exposure period used for initial efficacy assessment (above). After exposure, larvae were terminated by overdosing with an anaesthetic differing from the one being measured to avoid background contamination. The residual test solution was removed under vacuum and larvae washed in 12×300 μL of culture water, homogenised, extracted using acetonitrile containing an internal standard, and then centrifuged at 4000 rpm for 30 minutes. The supernatants were then analysed alongside samples of exposure solutions. LC-MS/MS Analysis was undertaken on a TSQ Vantage triple quadrupole mass spectrometer equipped with heated electrospray (HESI II) source coupled to Surveyor MS Pump Plus HPLC pump with HTC PAL autosampler (all Thermo Fisher Scientific, San Jose, CA). Chromatographic separation was achieved using a reversed-phase, 3 µm particle size, C18 Hypersil GOLD column 50 mm × 2.1 mm internal diameter (Thermo Scientific, San Jose CA, USA). Separation was achieved via a linear gradient of water and methanol (0.1% of ammonium hydroxide) increasing from 20% to 100% methanol over 1.5 minutes, maintained for a further 1.5 minutes before returning to the initial 20% at a flow rate of 500 µL/min. The HESI probe was operating in both positive (3.75 kV), and negative (2.7 kV) modes. The heated capillary temperature was set at 275 °C, the vaporizer temperature set at 350 °C, and N_2_ was employed as sheath and auxiliary gas. Quantification of analytes was performed using characteristic multiple reaction monitoring (MRM) with quantitation using the internal standard method.

### Whole brain functional imaging

Monitoring anaesthetic depth using a motor-mediated response does not account for the fact that neuromuscular blockade and central sensory suppression can occur independently of one another, potentially resulting in unresponsive, yet fully conscious, animals. Here we applied functional brain imaging to measure neural activity, using Tg(*elavl3*:GCaMP6s) zebrafish larvae with an M2 Aurora light sheet microscope (M squared, Glasgow, UK – see below for more details) to systematically compare the effectiveness of the anaesthetics ^64,65^.

Briefly, each larva was individually exposed to 4 mM tubocurarine to induce rapid paralysis and immobilisation without central nervous system suppression^66^. The larva was then placed into 1% low melting point agarose (containing 4 mM tubocurarine) and positioned at 45° in the dorsal-ventral plane in a bespoke 3D printed plastic recording chamber, orientated so that the collection objective was perpendicular to the top of the head, and the laser objective perpendicular to the side of the larva. After positioning, 3 ml of culture water was added to the stage chamber, fully immersing the larva. Dorsal-ventral optical sectioning was then undertaken on the light sheet microscope (10 equidistant Z-slices across 200 μm taken in ∼0.8 s, repeated 400 times for a total imaging duration of just over 5 minutes) to record baseline brain activity on the light sheet microscope with a 488 nm laser line, 10x illumination and 20x collection objectives. Images were captured via a Hamamatsu Orca Flash 4.0 camera (Hamamatsu Photonics, Hamamatsu City, Japan) at a resolution of 21524 dpi (vertical and horizontal resolution). After recording basal activity, the anaesthetic compound (or water or ethanol for the control treatments) was added to the larva on the stage, and left for a 3-minute diffusion period based on pilot data that demonstrated clear effects. After this, post-exposure imaging was undertaken for a further 5 minutes. Finally, the larva was removed and humanely killed by anaesthetic overdose. In the instance of functional brain imaging with the presentation of a light stimulus, a blue light flash was introduced on timepoint 100 (of 400), lasting for 8 timepoints (5 seconds). This was the case both for pre-exposure (baseline) and post-exposure imaging sessions.

Pre-exposure and exposure phase images were analysed following a process reported previously^64,65^, that uses a custom Python image processing pipeline (available at: https://exeter-zebrafish-research.gitlab.io/zebrafish-neuroanalyser-launcher/). Briefly, images were down-sampled and then 3D aligned against a labelled reference brain using 21 user-selected key points from a representative image stack. Images were automatically shift-corrected to allow for any x, y, and z drift for each region of interest (ROI). Quantitative data were then extracted as the mean of incorporated voxels for each time-point, generating a temporal activity profile for each registered brain region, again, as previously described^64,65^. In the instance of functional brain imaging with the presentation of a light stimulus, the data analysed included only the 10 timepoints directly succeeding the stimulus presentation, as we were interested in the responsiveness of the brain to external stimulation, rather than overall neural activity.

### Behavioural measures of tolerability

#### Locomotion and thigmotaxis

Locomotor activity and thigmotaxis, were assessed using a 4-way video tracking system (VideoTrack v2.5, Viewpoint, France)^67,68^. For these assessments, larvae were individually deposited into wells of a 24-well plate (Corning Incorporated, UK) containing 600 μl of test solution across 6 concentrations (including a control treatment). For each compound, treatments were spread across 4 plates, providing simultaneous measurement in 16 fish per treatment. Video tracking was undertaken for 40 minutes, during which there was alternation between a dark phase (under infrared lighting) and a light phase (under white light ca. 500 Lux). The protocol used comprised a 10 minute light acclimation period followed by 6 abrupt 100% light/dark transitions, each lasting 5 minutes, designed to stimulate movement in relatively inactive 4.5 dpf larvae as well as potentially supplying a mild anxiogenic stimulus^42^. From the resulting video recordings, the total distance travelled and the time spent moving were quantified. In addition, each well was divided virtually into an outer and inner zone of equal area and the percentage of time individuals spent moving in the outer zone was calculated as a measure of thigmotaxis, which is a well characterised measure of anxiety/boldness in larval zebrafish^42^.

#### Aversion

Olfaction-mediated avoidance behaviour (aversion) was assessed using a protocol based on that of Takesono *et al.*^69^. Briefly, ten 4.5 dpf WT larvae were placed in 15ml of culture water in the middle compartment of a 3-compartment test arena (Figure S4), and allowed to acclimate for 15 minutes. The final 5-minute period of the acclimation phase was videoed to assess pre-treatment movement (acclimation phase). The anaesthetic agent solution (at a volume between 20-60 μl, appropriate to achieve the desired concentration) was then carefully added to one side of the test chamber by pipette, and either culture water or solvent control, as appropriate, at the opposite end of the test chamber. The concentration ranges used to assess aversion were selected based on the absence of sedation, using data from the initial assessment of touch responsiveness (Table 1). The side of the arena into which each sample was added was alternated between tests. Cadaverine (20mM, pH 7.5), an odorant known to induce an aversive response in larval zebrafish^70^ was used as a positive control^69^. Pilot experiments using phenol red to track substance diffusion indicated that a 5-minute equilibration period was sufficient to create a chemical gradient across the test arena. After the equilibration period, the dividers were carefully removed from both sides of the test arena and the movement of larvae was videoed for 10 minutes (at 20 fps) in the dark using an infrared camera to reduce the potential for visual cues. Resultant videos were analysed using the multi-animal tracking software TRex^71^. For each trial, larval movement was analysed and extracted in the form of x-y coordinates from which the total distance moved and average speed for each group of fish tested were calculated. The mean centre of mass (centroid) for each experimental group was also calculated, and the shift of the centroid from the centre of the arena was used to indicate avoidance (aversion) of the compound compared with the administered control (water or solvent control).

#### Cardiovascular physiology

Cardiovascular effects were analysed using a previously reported methodology^72^. Due to the need for immobilisation, it was not possible to employ an untreated control and, as such, the lowest concentration used, a level at which light immobilisation was achieved, was considered the control against which all other treatments were compared. Each larva was exposed to the anaesthetic for 10 minutes and then embedded in 1% agarose containing the same concentration of anaesthetic within a well of a press-to-seal silicon isolator (Sigma-Aldrich, Poole, UK) attached to a transparent microscope slide. Larvae were then gently orientated onto their side and a drop of exposure solution added to prevent the gel from drying out. The sample size was n=8 for each test concentration. Five minute videos of the heart and peripheral vasculature dorsal to the swim bladder were then captured at 25fps (Grasshopper® GRAS-50S5C-C, Point Grey, Richmond, Canada) and at 120fps (Grasshopper® GRAS-03K2M-C, Point Grey, Richmond, Canada) respectively, at 10x magnification on an inverted light microscope (Leica DM IRB, Leica Microsystems UK Ltd., Milton Keynes, UK). Atrial and ventricular beat rates (ABR and VBR, respectively) were automatically quantified in beats per minute (BPM) across the whole experimental duration using MicroZebraLab™ (v3.5, ViewPoint, Lyon, France). In some cases, automatic detection was not possible, for example due to an excessively low beat rate under the highest concentration of MS222 (5mM). For these fish, data were collected manually by counting the heartbeat for 3 minutes to allow calculation of BPM. Vascular parameters, namely blood flow, linear velocity and dorsal aorta diameter were quantified using ZebraBlood™ (v3.4.6, ViewPoint, Lyon, France). From the blood flow data, surrogate measures of stroke volume (surrogate stroke volume or SSV) and cardiac output (surrogate cardiac output or SCO) were also calculated using blood flow as a surrogate measure for ventricular volume, as previously described^72^.

#### Data analysis

Approaches used for analysis varied according to the dataset analysed and are described below.

#### Functional brain imaging

For each ROI, the median intensity for that region was averaged across all time points within pre-exposure (baseline) or the exposure periods to obtain one value per region per fish for each period. Δf/f was calculated as (f1 − f0)/f0 (where f1 = mean fluorescence intensity after anaesthetic exposure, and f0 = mean baseline fluorescence intensity, i.e. intensity pre-exposure, for the same animal within that ROI). The resultant mean Δf/f for each treatment was compared with the corresponding ROI control values using a Wilcoxon rank sum test with a statistical significance threshold of p<0.05 (GraphPad Prism V10, GraphPad Software, Boston, Massachusetts, USA). In the case of functional brain imaging with visual stimulation, we only analysed the 10 time points directly succeeding the presentation of the light stimulus, in order to capture the stimulus-evoked brain response.

#### Locomotion

Total distance travelled by individual larvae was calculated as the percent change between acclimation and test periods, and analyses were conducted separately for light and dark periods^73^. For each analysis, linear mixed effect models were fitted separately, using the lme4 v 1.1 R package^74^. All models included an interaction between treatment and time (1^st^, 2^nd^, or 3^rd^ period of light or dark); individual larval ID was included as a random factor to account for individual level differences in movement. The residuals of all models were inspected visually to ensure a good fit, and if that was not the case, the values for overall movement were appropriately transformed. This was the case for 2-phenoxyethanol, light (square root transformation); benzocaine, light (square root transformation); etomidate, light (square root transformation); etomidate, dark (square root transformation); isoeugenol, light and dark (square root transformation); MS222, dark (square root transformation); quinaldine sulfate, light (squared) and quinaldine sulfate, dark (log_10_ transformation after adding 0.00001). Where a statistically significant interaction between treatment and time was observed, *post hoc* comparisons against the control treatment were undertaken using the emmeans v 1.10.3 R package^75^, allowing for Šidák adjustment of p values. When a significant interaction between treatment and time was not observed (in the case of isoeugenol, dark condition), the interaction was not included based on the Akaike Information Criterion (AIC), in order to observe the full effect of the fixed factors. Analyses were carried out using R v 4.3^76^.

#### Thigmotaxis

Thigmotaxis was assessed by quantifying the duration of time spent in the periphery of the test chamber (defined as the outer 50% of the surface area of the well^42^), versus the total duration of time spent moving for each minute of the experiment, through fitting binomial generalised linear mixed models (lme4 v 1.1 R package)^74^. Data for each anaesthetic compound were analysed separately. All models included an interaction between treatment (anaesthetic dosage) and light status (light/dark), and individual ID was included as a random factor. The distance travelled (as a z-score) within the same period of time was also included as a fixed factor, to account for any effects of the anaesthetics on overall movement. The residuals of all models were inspected visually to ensure good fit. For exposures to the highest concentration of etomidate, MS222, and the highest and second highest concentrations of quinaldine sulfate, there was no larval movement over the experimental duration; these treatments were thus excluded from the analysis. *Post hoc* comparisons with the control treatment were carried out using the emmeans v 1.10.3 R package^75^, allowing for Šidák adjustment of p values. Analyses were carried out using R v 4.3^76^.

#### Aversion

Aversion data were analysed by fitting a generalised least squares estimation model (GLS) to account for differences in heteroscedasticity between the different treatments, using the nlme v 3.1 R package^77^. The total shift of each group from the central point of the arena along the x axis was modelled against the treatment (anaesthetic vs control, vs positive control, cadaverine), the side of administration of the treatment (L/R) (to account for any side bias), and the average distance moved by each experimental group. The latter metric was used to account for any sedative effects that the anaesthetics may have had on the expression of aversive behaviour. Analyses were carried out using R v 4.3^76^.

#### Cardiovascular function

Statistical analyses of cardiovascular function were performed using GraphPad Prism 10 (GraphPad Software 10.0.0, Boston, USA). Data were first tested for normality and heteroscedasticity using Shapiro-Wilk and Bartlett’s or Levene’s tests, respectively. For data meeting the assumptions of parametric testing, a one-way analysis of variance (ANOVA) was applied, followed by Dunn’s *post-hoc* testing, comparing each anaesthetic concentration with the corresponding control group. For non-parametric testing, a Kruskal-Wallis test was used, followed by a Dunn’s *post-hoc* comparison comparing treated and control groups.

## Supporting information

Supplemental Results

## Author contributions

L.U.S., A.T., T.K., A.R., C.R.T., and M.J.W. conceived the research project and secured funding. S.D., M.M., A.P., C.R.T. and M.J.W. designed the experiments, with input from J.T.B. and M.T.. S.D., M.M., A.P., J.L.B., D.H.-B., H.R., J.C., and J.S.B., collected the data. S.D. and M.M. analysed the data, with input from C.H. and M.J.W.. M.T. collected and analysed the bioavailability data. S.D., M.M., C.R.T., and M.J.W. wrote the manuscript. All authors reviewed the manuscript and approved the final version.

## Acknowledgments

The authors would like to thank the team of the Aquatic Resources Centre of the University of Exeter, in particular Dave Maley and Alex Bell, as well as Gaby Wasser-Bennet, Teigan Veale, Siobhan Monaghan, and Michael Love. This project was funded by the UKRI (BBSRC: BB/V000411/1). L.U.S. is grateful for funding from the Swedish Research Council (Vetenskapsrådet 2022-01365).

## Data availability

All data will be made publicly available via Figshare upon publication.The Python code used for processing light sheet images including image registration to the z-brain atlas is publicly accessible here: https://exeter-zebrafish-research.gitlab.io/zebrafish-neuroanalyser-launcher/.

## References

1. Lidster, K., Readman, G. D., Prescott, M. J. & Owen, S. F. International survey on the use and welfare of zebrafish Danio rerio in research. J. Fish Biol. 90, 1891–1905 (2017).

2. European Commission. Union overview on the implementation of Directive 2010/63/EU on the protection of animals used for scientific purposes in the Member States of the European Union in 2018-2022. https://circabc.europa.eu/ui/group/8ee3c69a-bccb-4f22-89ca-277e35de7c63/library/16a0a839-8f80-4aef-951b-d49cfe21935e/details?download=true (2024).

3. Annual statistics of scientific procedures on living animals, Great Britain 2023. *GOV.UK* https://www.gov.uk/government/statistics/statistics-of-scientific-procedures-on-living-animals-great-britain-2023/annual-statistics-of-scientific-procedures-on-living-animals-great-britain-2023.

4. Irion, U. & Nüsslein-Volhard, C. Developmental genetics with model organisms. Proc. Natl. Acad. Sci. 119, e2122148119 (2022).

5. Patton, E. E., Zon, L. I. & Langenau, D. M. Zebrafish disease models in drug discovery: from preclinical modelling to clinical trials. Nat. Rev. Drug Discov. 20, 611–628 (2021).

6. Cressey, D. Fish-kill method questioned. Nature 506, 419–420 (2014).

7. Martins, T., Valentim, A. M., Pereira, N. & Antunes, L. M. Anaesthesia and analgesia in laboratory adult zebrafish: a question of refinement. Lab. Anim. 50, 476–488 (2016).

8. Heavner, J. E. Animal models and methods in anesthetic research. Methods Anim. Exp. 313–357 (2013).

9. Sneddon, L. U. Clinical Anesthesia and Analgesia in Fish. J. Exot. Pet Med. 21, 32–43 (2012).

10. Soldatov, A. A. Functional Effects of the Use of Anesthetics on Teleostean Fishes (Review). Inland Water Biol. 14, 67–77 (2021).

11. Summerfelt, R. Methods for fish biology. Am. Fish. Soc. (1990).

12. Marking, L. L. & Meyer, F. P. Are Better Anesthetics Needed in Fisheries? Fisheries 10, 2–5 (1985).

13. Gilderhus, P. A. Benzocaine as a Fish Anesthetic: Efficacy and Safety for Spawning-Phase Salmon. Progress. Fish-Cult. 52, 189–191 (1990).

14. Leach, M., Bowell, V., Allan, T. & Morton, D. Measurement of aversion to determine humane methods of anaesthesia and euthanasia | Animal Welfare | Cambridge Core. (2004).

15. Ross, L. G. & Ross, B. Anaesthetic and Sedative Techniques for Aquatic Animals. (John Wiley & Sons, 2009).

16. Laird, L. M. & Oswald, R. A note on the use of benzocaine (ethyl p-aminobenzoate) as a fish anaesthetic. Aquac. Res. 6, 92–94 (1975).

17. King V, W., Hooper, B., Hillsgrove, S., Benton, C. & Berlinsky, D. L. The use of clove oil, metomidate, tricaine methanesulphonate and 2-phenoxyethanol for inducing anaesthesia and their effect on the cortisol stress response in black sea bass (Centropristis striata L.). Aquac. Res. 36, 1442–1449 (2005).

18. Matthews, M. & Varga, Z. M. Anesthesia and euthanasia in zebrafish. ILAR J. 53, 192–204 (2012).

19. Readman, G. D., Owen, S. F., Murrell, J. C. & Knowles, T. G. Do Fish Perceive Anaesthetics as Aversive? PLOS ONE 8, e73773 (2013).

20. Wong, D., Keyserlingk, M. A. G. von, Richards, J. G. & Weary, D. M. Conditioned Place Avoidance of Zebrafish (Danio rerio) to Three Chemicals Used for Euthanasia and Anaesthesia. PLOS ONE 9, e88030 (2014).

21. Ferreira, J. M. et al. Behavioural Aversion and Cortisol Level Assessment When Adult Zebrafish Are Exposed to Different Anaesthetics. Biology 11, 1433 (2022).

22. Sánchez-Vázquez, F. J., Terry, M. I., Felizardo, V. O. & Vera, L. M. Daily Rhythms of Toxicity and Effectiveness of Anesthetics (MS222 and Eugenol) in Zebrafish (Danio Rerio). Chronobiol. Int. 28, 109–117 (2011).

23. Huang, W.-C. et al. Combined use of MS-222 (tricaine) and isoflurane extends anesthesia time and minimizes cardiac rhythm side effects in adult zebrafish. Zebrafish 7, 297–304 (2010).

24. Huang, H. et al. Toxicity, uptake kinetics and behavior assessment in zebrafish embryos following exposure to perfluorooctanesulphonicacid (PFOS). Aquat. Toxicol. 98, 139–147 (2010).

25. Gut, P., Reischauer, S., Stainier, D. Y. R. & Arnaout, R. Little fish, big data: Zebrafish as a model for cardiovascular and metabolic disease. Physiol. Rev. 97, 889–938 (2017).

26. MacPhail, R. C. et al. Locomotion in larval zebrafish: Influence of time of day, lighting and ethanol. NeuroToxicology 30, 52–58 (2009).

27. Mocho, J.-P. et al. A Multi-Site Assessment of Anesthetic Overdose, Hypothermic Shock, and Electrical Stunning as Methods of Euthanasia for Zebrafish (Danio rerio) Embryos and Larvae. Biology 11, 546 (2022).

28. Ohnesorge, N. et al. Assessment of the effect of tricaine (MS-222)-induced anesthesia on brain-wide neuronal activity of zebrafish (Danio rerio) larvae. Front. Neurosci. 18, (2024).

29. Cheng, R.-K., Jesuthasan, S. J. & Penney, T. B. Zebrafish forebrain and temporal conditioning. Philos. Trans. R. Soc. B Biol. Sci. 369, 20120462 (2014).

30. Salas, C., Broglio, C. & Rodríguez, F. Evolution of Forebrain and Spatial Cognition in Vertebrates: Conservation across Diversity. Brain. Behav. Evol. 62, 72–82 (2003).

31. Bonhomme, V. L. G., Boveroux, P., Brichant, J. F., Laureys, S. & Boly, M. Neural correlates of consciousness during general anesthesia using functional magnetic resonance imaging (fMRI). Arch. Ital. Biol. 150, 155–163 (2012).

32. Zhuang, Z. et al. Developmental neurotoxicity of anesthetic etomidate in zebrafish larvae: Alterations in motor function, neurotransmitter signaling, and lipid metabolism. J. Hazard. Mater. 494, 138598 (2025).

33. Neiffer, D. L. & Stamper, M. A. Fish Sedation, Anesthesia, Analgesia, and Euthanasia: Considerations, Methods, and Types of Drugs. ILAR J. 50, 343–360 (2009).

34. Mattson, N. S. & Riple, T. H. Metomidate, a better anesthetic for cod (*Gadus morhua*) in comparison with benzocaine, MS-222, chlorobutanol, and phenoxyethanol. Aquaculture 83, 89–94 (1989).

35. Hayashi, H. & Kawaguchi, M. Intraoperative monitoring of flash visual evoked potential under general anesthesia. Korean J. Anesthesiol. 70, 127–135 (2017).

36. Lee, H., Tanabe, S., Wang, S. & Hudetz, A. G. Differential effect of anesthesia on visual cortex neurons with diverse population coupling. Neuroscience 458, 108–119 (2021).

37. Liu, D., et al. Monochromatic Blue Light Activates Suprachiasmatic Nucleus Neuronal Activity and Promotes Arousal in Mice Under Sevoflurane Anesthesia. Front. Neural Circuits 14, (2020).

38. Bühler, A. & Carl, M. Zebrafish Tools for Deciphering Habenular Network-Linked Mental Disorders. Biomolecules 11, 324 (2021).

39. Xiao, T., Roeser, T., Staub, W. & Baier, H. A GFP-based genetic screen reveals mutations that disrupt the architecture of the zebrafish retinotectal projection. Development 132, 2955–2967 (2005).

40. Machnik, P., Biazar, N. & Schuster, S. Recordings in an integrating central neuron reveal the mode of action of isoeugenol. *Commun*. Biol. 6, 1–10 (2023).

41. Leyden, C. et al. Efficacy of Tricaine (MS-222) and Hypothermia as Anesthetic Agents for Blocking Sensorimotor Responses in Larval Zebrafish. Front. Vet. Sci. 9, (2022).

42. Schnörr, S. J., Steenbergen, P. J., Richardson, M. K. & Champagne, D. L. Measuring thigmotaxis in larval zebrafish. Behav. Brain Res. 228, 367–374 (2012).

43. Weber, R. A., Peleteiro, J. B., Martín, L. O. G. & Aldegunde, M. The efficacy of 2-phenoxyethanol, metomidate, clove oil and MS-222 as anaesthetic agents in the Senegalese sole (*Solea senegalensis* Kaup 1858). Aquaculture 288, 147–150 (2009).

44. McCord, C. L. et al. Concentration effects of three common fish anesthetics on Pacific hagfish (Eptatretus stoutii). Fish Physiol. Biochem. 46, 931–943 (2020).

45. Li, J. et al. Early Development of Functional Spatial Maps in the Zebrafish Olfactory Bulb. J. Neurosci. 25, 5784–5795 (2005).

46. Calvo-Ochoa, E. & Byrd-Jacobs, C. A. The Olfactory System of Zebrafish as a Model for the Study of Neurotoxicity and Injury: Implications for Neuroplasticity and Disease. Int. J. Mol. Sci. 20, 1639 (2019).

47. Braubach, O. R., Fine, A. & Croll, R. P. Distribution and functional organization of glomeruli in the olfactory bulbs of zebrafish (Danio rerio). J. Comp. Neurol. 520, 2317–2339 (2012).

48. Ghanawi, J., Monzer, S. & Saoud, I. P. Anaesthetic efficacy of clove oil, benzocaine, 2-phenoxyethanol and tricaine methanesulfonate in juvenile marbled spinefoot (iganus rivulatus). Aquac. Res. 44, 359–366 (2013).

49. Bakkers, J. Zebrafish as a model to study cardiac development and human cardiac disease. Cardiovasc. Res. 91, 279–288 (2011).

50. González-Rosa, J. M. Zebrafish Models of Cardiac Disease: From Fortuitous Mutants to Precision Medicine. Circ. Res. 130, 1803–1826 (2022).

51. Malone, M. H. et al. Laser-scanning velocimetry: A confocal microscopy method for quantitative measurement of cardiovascular performance in zebrafish embryos and larvae. BMC Biotechnol. 7, 40 (2007).

52. Craig, M. P., Gilday, S. D. & Hove, J. R. Dose-dependent effects of chemical immobilization on the heart rate of embryonic zebrafish. Lab Anim. 35, 41–47 (2006).

53. Neumcke, B., Schwarz, W. & Stämpfli, R. Block of Na channels in the membrane of myelinated nerve by benzocaine. Pflüg. Arch. 390, 230–236 (1981).

54. Frazier, D. T. & Narahashi, T. Tricaine (MS-222): Effects on ionic conductances of squid axon membranes. Eur. J. Pharmacol. 33, 313–317 (1975).

55. Kiessling, A., Johansson, D., Zahl, I. H. & Samuelsen, O. B. Pharmacokinetics, plasma cortisol and effectiveness of benzocaine, MS-222 and isoeugenol measured in individual dorsal aorta-cannulated Atlantic salmon (Salmo salar) following bath administration. Aquaculture 286, 301–308 (2009).

56. Mußhoff, U., Madeja, M., Binding, N., Witting, U. & Speckmann, E.-J. Effects of 2-phenoxyethanol on N-methyl-d-aspartate (NMDA) receptor-mediated ion currents. Arch. Toxicol. 73, 55–59 (1999).

57. Priborsky, J. & Velisek, J. A Review of Three Commonly Used Fish Anesthetics. Rev. Fish. Sci. Aquac. 26, 417–442 (2018).

58. Zahl, I. H., Samuelsen, O. & Kiessling, A. Anaesthesia of farmed fish: implications for welfare. Fish Physiol. Biochem. 38, 201–218 (2012).

59. Hahnenkamp, K. et al. Local anaesthetics inhibit signalling of human NMDA receptors recombinantly expressed in *Xenopus laevis* oocytes: role of protein kinase C. Br. J. Anaesth. 96, 77–87 (2006).

60. Ashton, D. & Wauquier, A. Modulation of a GABA-Ergic Inhibitory Circuit in the In Vitro Hippocampus by Etomidate Isomers. Anesth. Analg. 64, 975 (1985).

61. Zhang, H. et al. The Application and Pharmaceutical Development of Etomidate: Challenges and Strategies. Mol. Pharm. (2024) doi:10.1021/acs.molpharmaceut.4c00325.

62. Grasshoff, C., Drexler, B., Rudolph, U. & Antkowiak, B. Anaesthetic Drugs: Linking Molecular Actions to Clinical Effects. http://www.eurekaselect.com.

63. von Krogh, K., Higgins, J., Saavedra Torres, Y. & Mocho, J.-P. Screening of Anaesthetics in Adult Zebrafish (Danio rerio) for the Induction of Euthanasia by Overdose. Biology 10, 1133 (2021).

64. Winter, M. J., et al. Functional brain imaging in larval zebrafish for characterising the effects of seizurogenic compounds acting via a range of pharmacological mechanisms. (2021).

65. Winter, M. J. et al. 4-dimensional functional profiling in the convulsant-treated larval zebrafish brain. Sci. Rep. 7, 6581 (2017).

66. Favre-Bulle, I. A., Vanwalleghem, G., Taylor, M. A., Rubinsztein-Dunlop, H. & Scott, E. K. Cellular-Resolution Imaging of Vestibular Processing across the Larval Zebrafish Brain. Curr. Biol. 28, 3711–3722.e3 (2018).

67. Winter, M. J. et al. Validation of a larval zebrafish locomotor assay for assessing the seizure liability of early-stage development drugs. J. Pharmacol. Toxicol. Methods 57, 176–187 (2008).

68. Gould, S. L. et al. Exposure Effects of Environmentally Relevant Concentrations of the Tricyclic Antidepressant Amitriptyline in Early Life Stage Zebrafish. Environ. Sci. Technol. 58, 13194–13204 (2024).

69. Takesono, A. et al. Zinc oxide nanoparticles disrupt development and function of the olfactory sensory system impairing olfaction-mediated behaviour in zebrafish. Environ. Int. 180, 108227 (2023).

70. Dieris, M. Amine detection in aquatic organisms: receptor evolution, neuronal circuits and behavior in the model organism zebrafish. (Universität zu Köln, 2018).

71. Walter, T. & Couzin, I. D. TRex, a fast multi-animal tracking system with markerless identification, and 2D estimation of posture and visual fields. Elife 10, e64000 (2021).

72. Parker, T. et al. A multi-endpoint *in vivo* larval zebrafish (*Danio rerio*) model for the assessment of integrated cardiovascular function. J. Pharmacol. Toxicol. Methods 69, 30–38 (2014).

73. Hillman, C., Kearn, J. & Parker, M. O. Protocol for investigating light/dark locomotion in larval stage zebrafish using a standardized behavioral assay. STAR Protoc. 5, 103346 (2024).

74. Bates, D., Maechler, M., Bolker, B. & Walker, S. lme4: Linear mixed-effects models using Eigen and S4. R Package Version 1, 1–23 (2014).

75. Lenth, R. Emmeans: Estimated marginal means, aka least-squares means. R Package Version 1, (2018).

76. R Development Core Team. R: A Language and Environment for Statistical Computing. Vienna, Austria: R Foundation for Statistical Computing; 2014. R Foundation for Statistical Computing. … Free. Available Internet *Httpwww*R-Proj. … (2015).

77. Pinheiro, J., Bates, D., DebRoy, S. & Sarkar, D. R Core Team (2014) nlme: linear and nonlinear mixed effects models. R package version 3.1-117. Available H TtpCRAN R-Proj. Orgpackage Nlme (2014).

