## Supplemental Results for "Evaluating anaesthetics for improving scientific research and welfare using larval zebrafish"

### SUPPLEMENTARY MATERIAL

### Supplementary results

#### In vivo $Ca^{2+}$ functional brain imaging without visual stimulation

##### 2-phenoxyethanol

**Table S1. Mean fluorescence intensity in brain regions of interest (ROIs) measured using light-sheet imaging after a 3-minute exposure to 2-phenoxyethanol.** Bold denotes ROIs where anaesthetic administration was found to have a statistically significant effect on activity.

| ROI | Control | | 2mM | | 3mM | | 5mM | | 7mM | | Kruskal-Wallis statistic | p-value |
|---|---|---|---|---|---|---|---|---|---|---|---|---|
| | Mean | SD | Mean | SD | Mean | SD | Mean | SD | Mean | SD | | |
| **Telencephalon** | **6.505** | **36.584** | **1.418** | **19.337** | **4.396** | **46.444** | **-46.479** | **13.443** | **-42.244** | **13.419** | **17.21** | **0.0018** |
| **Olfactory Bulbs** | -12.038 | 38.576 | 6.797 | 57.006 | 6.956 | 30.586 | 55.011 | 252.363 | -25.001 | 39.190 | 5.052 | 0.282 |
| **Anterior commissure** | **37.691** | **51.251** | **7.385** | **24.690** | **32.473** | **91.006** | **-48.818** | **18.984** | **-57.066** | **15.385** | **23.3** | **0.0001** |
| **Pallium** | **7.766** | **40.336** | **-0.111** | **14.472** | **19.978** | **48.595** | **-42.441** | **14.732** | **-36.230** | **19.105** | **17.62** | **0.0015** |
| **Subpallium** | **3.866** | **40.960** | **-1.912** | **29.553** | **-6.640** | **35.319** | **-50.109** | **9.877** | **-42.805** | **12.194** | **17.06** | **0.0019** |
| **Diencephalon** | **0.959** | **29.187** | **-5.152** | **58.178** | **-18.410** | **29.157** | **-64.598** | **11.243** | **-46.047** | **20.628** | **19.36** | **0.0007** |
| **Dorsal thalamus** | **-5.961** | **33.702** | **-6.530** | **77.156** | **-23.887** | **32.622** | **-65.422** | **13.742** | **-40.302** | **19.875** | **16.23** | **0.0027** |
| **Eminentia thalami** | -1.396 | 44.986 | -1.628 | 32.025 | 5.819 | 65.658 | -28.268 | 29.836 | -31.922 | 18.975 | 5.914 | 0.2057 |
| **Habenulae** | **-11.530** | **22.819** | **-0.337** | **19.733** | **12.439** | **47.507** | **-40.423** | **14.323** | **-18.988** | **20.902** | **13.02** | **0.0112** |
| **Intermediate hypothalamus** | **-2.037** | **34.367** | **-2.722** | **46.879** | **-25.600** | **20.623** | **-68.917** | **13.549** | **-52.794** | **23.870** | **20.81** | **0.0003** |
| **Pineal** | **-6.797** | **36.007** | **-11.582** | **23.122** | **1.513** | **57.954** | **-31.179** | **21.616** | **12.431** | **33.607** | **7.915** | **0.0948** |
| **Posterior tuberculum** | **7.294** | **33.463** | **-6.623** | **58.092** | **-29.367** | **19.400** | **-67.277** | **13.115** | **-46.083** | **23.728** | **20.78** | **0.0004** |
| **Preoptic area** | **4.668** | **26.673** | **-7.711** | **34.883** | **-7.849** | **45.792** | **-53.570** | **13.414** | **-43.289** | **21.744** | **18.21** | **0.0011** |
| **Pretectum** | **-5.764** | **32.813** | **-0.953** | **68.990** | **-22.887** | **31.311** | **-63.563** | **13.999** | **-41.444** | **15.326** | **14.84** | **0.005** |

| | | | | | | | | | | | | |
|---|---|---|---|---|---|---|---|---|---|---|---|---|
| **Rostral hypothalamus** | **-2.511** | **29.500** | **0.899** | **59.326** | **-22.235** | **29.911** | **-62.142** | **13.297** | **-36.926** | **27.360** | **16.37** | **0.0026** |
| **Ventral thalamus** | **7.689** | **27.919** | **-11.593** | **58.509** | **-27.322** | **29.430** | **-68.675** | **12.830** | **-40.975** | **22.961** | **18.47** | **0.001** |
| **Mesencephalon** | **-0.487** | **32.103** | **-12.119** | **51.244** | **-21.887** | **26.142** | **-68.732** | **12.281** | **-49.638** | **20.330** | **20.8** | **0.0003** |
| **Tectum striatum periventiculare** | **1.114** | **35.673** | **-15.837** | **44.943** | **-21.089** | **23.194** | **-66.855** | **12.363** | **-43.016** | **21.560** | **19.19** | **0.0007** |
| **Tectum neuropil** | **-1.191** | **33.388** | **-5.485** | **56.941** | **-21.066** | **25.430** | **-59.142** | **16.300** | **-45.108** | **19.104** | **17.83** | **0.0013** |
| **Tegmentum** | **-0.594** | **33.811** | **-8.730** | **53.054** | **-24.451** | **26.883** | **-75.388** | **12.401** | **-58.644** | **17.628** | **21.9** | **0.0002** |
| **Torus longitudinaris** | 17.184 | 48.995 | 2.192 | 26.845 | 31.143 | 46.796 | -15.556 | 40.773 | 17.201 | 30.103 | 5.206 | 0.2668 |
| **Torus semicircularis** | **1.121** | **39.553** | **-6.475** | **58.238** | **-22.681** | **23.775** | **-57.136** | **13.106** | **-36.094** | **30.648** | **14.48** | **0.0059** |
| **Rhombencephalon** | **-4.749** | **27.404** | **-18.064** | **42.381** | **-13.114** | **49.573** | **-75.038** | **10.078** | **-65.342** | **16.421** | **23.26** | **0.0002** |
| **Area postrema** | 14.870 | 42.990 | -6.517 | 30.233 | -20.466 | 29.533 | -35.510 | 30.664 | 12.304 | 51.750 | 8.059 | 0.0895 |
| **Cerebellum** | **-14.823** | **26.730** | **-15.259** | **51.878** | **-18.031** | **31.758** | **-73.439** | **9.411** | **-63.643** | **15.017** | **21.03** | **0.0003** |
| **Corpus cerebelli** | **-15.075** | **27.421** | **-14.365** | **53.154** | **-19.293** | **31.105** | **-73.477** | **9.075** | **-62.139** | **17.734** | **20.32** | **0.0004** |
| **Eminentia granularis** | **-2.590** | **22.757** | **-2.402** | **51.592** | **-10.816** | **40.190** | **-65.911** | **10.992** | **-54.980** | **17.708** | **22.16** | **0.0002** |
| **Inferior olive** | **-10.194** | **21.282** | **-17.836** | **40.156** | **-10.957** | **48.993** | **-78.737** | **9.250** | **-68.474** | **18.074** | **24.21** | **<0.0001** |
| **Interpeduncular nucleus** | **-10.759** | **37.187** | **6.812** | **49.857** | **-25.788** | **24.069** | **-56.863** | **9.376** | **-49.319** | **22.767** | **16.39** | **0.0025** |
| **Lateral reticular nucleus** | **71.745** | **71.577** | **-5.042** | **19.578** | **-9.761** | **42.005** | **-39.780** | **25.459** | **-21.175** | **45.945** | **15.9** | **0.0032** |
| **Lobus caudalis cerebelli** | **-18.222** | **29.519** | **-17.243** | **40.128** | **-12.214** | **38.175** | **-64.387** | **14.903** | **-54.975** | **14.687** | **16.02** | **0.003** |
| **Locus coeruleus** | **-3.813** | **28.723** | **-10.280** | **56.690** | **-19.460** | **38.697** | **-72.601** | **10.108** | **-66.920** | **13.103** | **23.88** | **<0.0001** |
| **Mauthner** | **4.803** | **37.458** | **-11.202** | **47.138** | **-8.555** | **70.566** | **-71.580** | **10.449** | **-64.026** | **12.782** | **21.17** | **0.0003** |
| **Medial vestibular nucleus** | **-5.572** | **29.307** | **-11.594** | **48.087** | **-18.312** | **35.959** | **-73.898** | **10.218** | **-65.245** | **13.277** | **21.64** | **0.0002** |

| | | | | | | | | | | | | |
|---|---|---|---|---|---|---|---|---|---|---|---|---|
| **Noradrenergic neurons of the interfascicular and vagal areas** | **9.780** | **27.663** | **-18.867** | **16.930** | **0.360** | **44.388** | **-55.910** | **20.851** | **-31.925** | **35.000** | **18.21** | **0.0011** |
| **Raphe-inferior** | **-3.546** | **29.833** | **-27.680** | **37.479** | **-22.810** | **31.273** | **-74.307** | **11.768** | **-53.770** | **21.626** | **19.62** | **0.0006** |
| **Raphe-superior** | **-27.100** | **27.484** | **-7.756** | **46.925** | **-24.103** | **26.259** | **-56.841** | **10.139** | **-52.629** | **11.395** | **16.59** | **0.0023** |
| **Tangential vestibular nucleus** | **23.811** | **43.195** | **-5.258** | **42.052** | **-16.309** | **36.771** | **-58.900** | **11.956** | **-59.224** | **16.029** | **21.07** | **0.0003** |
| **Valvula cerebelli** | **-8.493** | **37.255** | **-8.408** | **61.191** | **-12.579** | **30.525** | **-63.066** | **17.015** | **-44.736** | **26.409** | **14.35** | **0.0063** |
| **Eyes** | **-1.975** | **16.412** | **6.646** | **39.961** | **13.141** | **16.771** | **-26.884** | **15.822** | **-13.461** | **20.746** | **13** | **0.0113** |
| **Olfactory epithelium** | 9.890 | 31.980 | 3.356 | 22.067 | 4.578 | 21.330 | -23.199 | 25.842 | -3.016 | 25.011 | 7.103 | 0.1305 |
| **Vagal ganglia** | **54.929** | **55.369** | **-7.749** | **17.706** | **38.957** | **51.835** | **-2.306** | **41.878** | **38.414** | **30.707** | **12.47** | **0.0142** |
| **Spinal cord** | **-8.857** | **26.219** | **-26.275** | **35.896** | **-16.296** | **29.076** | **-78.796** | **14.302** | **-68.011** | **24.124** | **22.83** | **0.0001** |
| **Neuropil region** | **-9.743** | **25.569** | **-23.621** | **38.372** | **-7.864** | **49.577** | **-79.844** | **10.245** | **-68.986** | **25.674** | **22.3** | **0.0002** |

#### Benzocaine

**Table S2. Mean fluorescence intensity in brain regions of interest (ROIs) measured using light-sheet imaging after a 3-minute exposure to benzocaine.** Bold denotes ROIs where anaesthetic administration was found to have a statistically significant effect on activity.

| ROI | Control | | 0.15mM | | 0.30mM | | 0.45mM | | 0.60mM | | 0.80mM | | Kruskal-Wallis statistic | p-value |
|---|---|---|---|---|---|---|---|---|---|---|---|---|---|---|
| | Mean | SD | Mean | SD | Mean | SD | Mean | SD | Mean | SD | Mean | SD | | |
| Telencephalon | 14.576 | 19.265 | 20.124 | 34.199 | 22.994 | 22.151 | -8.335 | 36.640 | 28.114 | 97.733 | -3.729 | 102.538 | 5.431 | 0.2665 |
| Olfactory Bulbs | 17.732 | 46.732 | 18.050 | 61.331 | 22.041 | 18.164 | 4.124 | 44.965 | 75.637 | 196.321 | -37.504 | 32.695 | 8.847 | 0.1153 |
| Anterior commissure | 14.786 | 24.588 | 7.439 | 48.120 | 24.921 | 17.506 | -17.879 | 38.171 | -8.570 | 68.897 | 463.062 | 973.470 | 7.895 | 0.1621 |
| Pallium | 11.484 | 15.621 | 22.769 | 37.747 | 15.437 | 24.142 | -8.304 | 39.259 | 26.238 | 106.077 | -0.487 | 138.456 | 6.401 | 0.2691 |
| Subpallium | 17.701 | 13.491 | 22.512 | 34.489 | 19.392 | 19.508 | -1.741 | 43.251 | 14.166 | 73.317 | 24.719 | 172.500 | 6.603 | 0.2518 |
| Diencephalon | **28.401** | **36.736** | **2.251** | **46.059** | **1.662** | **29.655** | **-26.932** | **35.329** | **-24.269** | **39.358** | **-60.001** | **19.529** | **20.28** | **0.0011** |
| Dorsal thalamus | **30.408** | **34.463** | **0.056** | **39.288** | **0.901** | **36.592** | **-18.809** | **36.087** | **-24.419** | **37.189** | **-46.149** | **28.103** | **15.99** | **0.0069** |
| Eminentia thalami | **29.493** | **25.821** | **14.475** | **22.909** | **6.646** | **26.424** | **10.957** | **76.519** | **-7.196** | **40.572** | **-74.185** | **14.496** | **23.29** | **0.0003** |
| Habenulae | **12.257** | **15.548** | **17.752** | **33.319** | **-1.974** | **14.949** | **-14.948** | **28.302** | **15.453** | **54.079** | **-54.230** | **28.427** | **17.96** | **0.003** |
| Intermediate hypothalamus | **16.540** | **30.167** | **3.796** | **54.664** | **4.189** | **27.673** | **-32.877** | **33.884** | **-25.013** | **52.077** | **-64.371** | **15.602** | **18.43** | **0.0025** |
| Pineal | -2.681 | 11.371 | 4.321 | 43.589 | 8.400 | 18.777 | 5.542 | 52.881 | 15.185 | 84.582 | -19.501 | 46.326 | 1.972 | 0.8531 |
| Posterior tuberculum | **29.983** | **50.650** | **-2.239** | **47.519** | **3.554** | **37.750** | **-28.816** | **41.359** | **-31.464** | **38.308** | **-63.093** | **14.481** | **19.29** | **0.0017** |
| Preoptic area | **30.795** | **22.721** | **8.341** | **39.561** | **12.214** | **20.764** | **-14.075** | **52.259** | **-13.386** | **37.227** | **-67.242** | **17.562** | **24.1** | **0.002** |
| Pretectum | **28.786** | **35.002** | **6.498** | **47.286** | **2.207** | **32.725** | **-19.513** | **32.327** | **-16.980** | **48.670** | **-48.671** | **22.353** | **16.02** | **0.0068** |
| Rostral hypothalamus | **31.933** | **31.473** | **9.358** | **45.001** | **9.730** | **34.735** | **-21.176** | **37.531** | **-15.669** | **48.048** | **-50.301** | **19.947** | **18.44** | **0.0024** |

| | | | | | | | | | | | | | | |
|---|---|---|---|---|---|---|---|---|---|---|---|---|---|---|
| **Ventral thalamus** | **29.909** | **36.066** | **2.823** | **48.409** | **5.884** | **39.707** | **-25.577** | **35.219** | **-28.573** | **36.570** | **-51.391** | **25.852** | **17.01** | **0.0045** |
| **Mesencephalon** | **22.283** | **30.212** | **7.117** | **52.009** | **1.318** | **30.935** | **-21.593** | **33.367** | **-13.136** | **53.224** | **-54.582** | **20.423** | **15.36** | **0.0089** |
| **Tectum striatum periventiculare** | **22.848** | **28.231** | **6.784** | **51.179** | **-1.264** | **30.265** | **-16.209** | **42.692** | **3.718** | **77.957** | **-53.457** | **20.158** | **14.2** | **0.0144** |
| **Tectum neuropil** | **20.084** | **30.455** | **6.613** | **38.692** | **3.660** | **26.075** | **-15.933** | **30.219** | **-7.639** | **46.743** | **-51.102** | **21.785** | **17.47** | **0.0037** |
| **Tegmentum** | **19.432** | **39.367** | **7.207** | **60.290** | **3.208** | **37.118** | **-30.980** | **33.826** | **-28.319** | **51.385** | **-61.715** | **19.489** | **18.72** | **0.0022** |
| **Torus longitudinaris** | 18.332 | 24.791 | -2.251 | 29.621 | 7.820 | 24.847 | -0.960 | 39.488 | 69.228 | 203.486 | -5.739 | 64.055 | 3.664 | 0.5987 |
| **Torus semicircularis** | **19.226** | **24.989** | **-3.115** | **31.239** | **4.466** | **24.185** | **-28.766** | **30.995** | **-17.043** | **44.284** | **-51.158** | **24.416** | **18.17** | **0.0027** |
| **Rhombencephalon** | **13.662** | **40.233** | **-4.366** | **43.905** | **-5.090** | **32.731** | **-34.557** | **32.930** | **-30.309** | **53.611** | **-72.894** | **17.237** | **19.5** | **0.0015** |
| **Area postrema** | **4.172** | **31.314** | **7.012** | **30.611** | **-12.662** | **27.965** | **-23.328** | **45.674** | **-3.508** | **62.178** | **-67.643** | **22.335** | **18.31** | **0.0026** |
| **Cerebellum** | **11.801** | **33.674** | **-4.002** | **44.470** | **-7.841** | **32.350** | **-31.256** | **33.370** | **-29.307** | **47.756** | **-66.656** | **17.488** | **17.43** | **0.0038** |
| **Corpus cerebelli** | **10.653** | **36.841** | **-4.409** | **40.800** | **-10.300** | **32.253** | **-30.796** | **35.084** | **-31.635** | **45.771** | **-66.935** | **17.092** | **17.93** | **0.003** |
| **Eminentia granularis** | 6.279 | 28.218 | 9.242 | 41.781 | -5.382 | 12.683 | -31.883 | 27.203 | -10.830 | 68.508 | -14.246 | 79.889 | 7.033 | 0.2182 |
| **Inferior olive** | **-5.576** | **38.760** | **-20.097** | **34.772** | **-4.244** | **29.778** | **-48.254** | **29.629** | **-34.338** | **53.207** | **-64.491** | **20.800** | **16.18** | **0.0063** |
| **Interpeduncular nucleus** | **-8.166** | **22.490** | **-11.401** | **26.789** | **5.700** | **22.586** | **-39.777** | **24.182** | **-33.526** | **33.563** | **-57.996** | **20.897** | **22.2** | **0.0005** |
| **Lateral reticular nucleus** | **19.491** | **27.345** | **-19.909** | **32.868** | **-5.439** | **20.571** | **-51.857** | **28.438** | **-21.623** | **42.927** | **-34.203** | **38.859** | **15.9** | **0.0071** |
| **Lobus caudalis cerebelli** | **11.790** | **26.057** | **2.404** | **36.073** | **-7.341** | **26.228** | **-29.814** | **37.496** | **-23.792** | **48.410** | **-65.264** | **19.719** | **19.21** | **0.0018** |

| | | | | | | | | | | | | | | |
|---|---|---|---|---|---|---|---|---|---|---|---|---|---|---|
| Locus coeruleus | 8.869 | 44.110 | -9.127 | 45.170 | -5.612 | 36.601 | -46.115 | 24.535 | -36.941 | 45.412 | -71.453 | 15.743 | 22.98 | 0.0003 |
| Mauthner | 10.950 | 47.404 | -2.142 | 56.544 | 1.355 | 34.680 | -34.506 | 33.288 | -36.286 | 38.394 | -74.468 | 15.147 | 21.71 | 0.0006 |
| Medial vestibular nucleus | 18.167 | 50.744 | -4.086 | 45.629 | -6.416 | 28.927 | -36.141 | 33.660 | -28.881 | 50.193 | -80.842 | 15.294 | 22.16 | 0.0005 |
| Noradrenergic neurons of the interfascicular and vagal areas | 7.341 | 32.664 | -1.325 | 39.540 | -18.136 | 20.533 | -40.901 | 36.298 | -3.007 | 76.936 | -5.833 | 166.626 | 12.28 | 0.0311 |
| Raphe-inferior | 19.393 | 47.308 | -7.079 | 39.673 | 0.452 | 43.215 | -30.701 | 35.189 | -22.384 | 56.445 | -70.265 | 22.005 | 20.04 | 0.0012 |
| Raphe-superior | -5.465 | 30.404 | -15.296 | 27.581 | -9.690 | 22.734 | -29.857 | 24.932 | -22.520 | 32.346 | -50.386 | 24.564 | 12.42 | 0.0295 |
| Tangential vestibular nucleus | 13.232 | 32.924 | -2.636 | 38.939 | -7.685 | 26.052 | -45.603 | 27.054 | -38.353 | 34.239 | -60.263 | 26.582 | 22.2 | 0.0005 |
| Valvula cerebelli | 8.344 | 33.482 | 7.481 | 47.438 | -4.625 | 44.989 | -13.741 | 57.993 | -21.493 | 47.110 | -53.560 | 24.915 | 10.47 | 0.0629 |
| Eyes | 10.313 | 19.163 | 17.031 | 37.614 | 5.740 | 23.851 | -7.866 | 14.623 | -0.321 | 33.909 | 6.292 | 57.820 | 3.879 | 0.567 |
| Olfactory epithelium | 21.216 | 17.940 | 33.710 | 52.721 | 28.753 | 30.433 | 22.251 | 68.291 | 267.330 | 708.789 | -27.838 | 31.393 | 10.81 | 0.0552 |
| Vagal ganglia | 20.126 | 18.267 | 11.559 | 45.563 | -2.277 | 17.196 | -13.375 | 39.771 | 55.710 | 139.300 | 8.700 | 103.496 | 7.914 | 0.1611 |
| Spinal cord | 19.329 | 54.966 | -6.325 | 47.121 | -4.748 | 29.786 | -43.041 | 26.040 | -22.819 | 67.843 | -67.731 | 23.691 | 17.45 | 0.0037 |
| Neuropil region | 13.282 | 54.857 | -9.117 | 49.443 | -6.313 | 37.718 | -48.740 | 26.879 | -35.508 | 58.804 | -59.418 | 19.468 | 15.66 | 0.0079 |

#### Etomidate

**Table S3. Mean fluorescence intensity in brain regions of interest (ROIs) measured using light-sheet imaging after a 3-minute exposure to etomidate.** Bold denotes ROIs where anaesthetic administration was found to have a statistically significant effect on activity.

| ROI | Control | | 2µM | | 4µM | | 8µM | | Kruskal-Wallis statistic | p-value |
|---|---|---|---|---|---|---|---|---|---|---|
| | Mean | SD | Mean | SD | Mean | SD | Mean | SD | | |
| **Telencephalon** | **7.482** | **13.826** | **34.943** | **21.576** | **-7.027** | **16.255** | **-8.559** | **14.732** | **16.9** | **0.0007** |
| Olfactory Bulbs | 14.299 | 45.192 | 58.478 | 76.184 | -3.701 | 34.923 | 25.064 | 114.100 | 3.105 | 0.3757 |
| Anterior commissure | 1.266 | 48.744 | 34.065 | 40.071 | -21.406 | 38.532 | 5.232 | 34.887 | 7.731 | 0.0519 |
| **Pallium** | **7.783** | **14.344** | **33.025** | **21.521** | **-4.397** | **20.678** | **-0.373** | **9.146** | **13.01** | **0.0046** |
| **Subpallium** | **1.945** | **14.039** | **36.614** | **30.648** | **-6.199** | **12.970** | **-8.789** | **12.164** | **12.37** | **0.0062** |
| **Diencephalon** | **-18.296** | **24.372** | **21.167** | **64.858** | **-30.359** | **25.881** | **-38.529** | **11.970** | **9.164** | **0.0272** |
| Dorsal thalamus | -21.809 | 28.407 | 7.745 | 55.807 | -33.163 | 25.419 | -43.356 | 13.055 | 7.109 | 0.0685 |
| Eminentia thalami | 11.891 | 31.747 | 22.774 | 44.745 | -9.857 | 22.299 | -6.237 | 23.137 | 2.633 | 0.4518 |
| **Habenulae** | **-2.541** | **10.315** | **18.788** | **19.628** | **-6.714** | **22.465** | **-13.284** | **12.958** | **10.76** | **0.0131** |
| Intermediate hypothalamus | -25.937 | 23.397 | 18.866 | 54.681 | -34.753 | 30.102 | -42.727 | 15.873 | 7.721 | 0.0521 |
| **Pineal** | **-7.066** | **13.074** | **12.266** | **26.347** | **-12.005** | **22.461** | **-26.835** | **15.081** | **10.53** | **0.0146** |
| Posterior tuberculum | -25.355 | 31.382 | 21.932 | 79.283 | -36.007 | 29.616 | -42.410 | 14.466 | 6.145 | 0.1048 |
| Preoptic area | -5.986 | 35.513 | 23.305 | 57.358 | -22.048 | 19.107 | -19.337 | 10.779 | 4.989 | 0.1726 |
| Pretectum | -24.787 | 24.968 | 11.100 | 64.602 | -32.142 | 22.821 | -41.636 | 14.842 | 5.816 | 0.1209 |
| **Rostral hypothalamus** | **-17.419** | **29.587** | **23.130** | **67.885** | **-34.067** | **24.699** | **-37.727** | **14.367** | **9.132** | **0.0276** |
| Ventral thalamus | -21.958 | 25.825 | 10.003 | 65.678 | -32.917 | 27.386 | -42.534 | 11.268 | 7.189 | 0.0661 |
| **Mesencephalon** | **-26.475** | **21.563** | **8.389** | **54.189** | **-37.641** | **24.705** | **-44.315** | **11.002** | **9.245** | **0.0262** |
| Tectum striatum periventiculare | -26.164 | 23.895 | 9.902 | 57.380 | -37.726 | 24.957 | -43.457 | 12.913 | 9.076 | 0.2833 |

| | | | | | | | | | | |
|---|---|---|---|---|---|---|---|---|---|---|
| **Tectum neuropil** | -14.572 | 30.303 | 3.811 | 39.329 | -30.846 | 22.298 | -32.984 | 12.490 | 7.539 | 0.0566 |
| **Tegmentum** | **-34.312** | **20.905** | **12.441** | **71.452** | **-41.925** | **27.686** | **-52.111** | **11.566** | **8.136** | **0.0433** |
| **Torus longitudinaris** | -18.806 | 13.174 | 12.279 | 49.239 | -23.013 | 26.666 | -18.280 | 20.493 | 4.279 | 0.2328 |
| **Torus semicircularis** | **-1.423** | **47.231** | **6.444** | **33.371** | **-32.373** | **16.307** | **-35.487** | **13.087** | **10.21** | **0.0169** |
| **Rhombencephalon** | **-31.283** | **18.905** | **14.894** | **75.887** | **-43.590** | **23.849** | **-51.104** | **10.851** | **8.775** | **0.0324** |
| **Area postrema** | -19.690 | 24.544 | 5.266 | 50.708 | -25.401 | 32.727 | -32.915 | 21.902 | 3.639 | 0.3032 |
| **Cerebellum** | **-31.063** | **19.721** | **14.791** | **74.018** | **-44.043** | **22.012** | **-50.498** | **13.036** | **11.01** | **0.0117** |
| **Corpus cerebelli** | **-31.266** | **18.774** | **11.176** | **75.127** | **-43.436** | **22.737** | **-49.536** | **12.883** | **8.193** | **0.0422** |
| **Eminentia granularis** | **-17.330** | **30.663** | **32.391** | **85.774** | **-40.651** | **20.633** | **-43.995** | **17.993** | **8.912** | **0.0305** |
| **Inferior olive** | -35.234 | 24.919 | 22.089 | 78.318 | -43.151 | 25.327 | -50.534 | 9.928 | 6.979 | 0.0726 |
| **Interpeduncular nucleus** | **-0.886** | **22.079** | **12.296** | **30.407** | **-31.816** | **19.866** | **-25.624** | **14.847** | **13.11** | **0.0044** |
| **Lateral reticular nucleus** | -19.546 | 20.270 | 24.965 | 99.146 | -32.149 | 17.052 | -31.608 | 23.129 | 4.423 | 0.2193 |
| **Lobus caudalis cerebelli** | -27.760 | 19.852 | 2.931 | 50.328 | -38.638 | 22.864 | -46.490 | 16.669 | 7.069 | 0.0697 |
| **Locus coeruleus** | -30.681 | 14.706 | -3.774 | 48.139 | -37.641 | 23.721 | -42.851 | 13.308 | 5.912 | 0.116 |
| **Mauthner** | **-23.486** | **25.413** | **6.598** | **50.529** | **-36.437** | **24.135** | **-47.072** | **12.335** | **8.244** | **0.0412** |
| **Medial vestibular nucleus** | **-32.744** | **19.516** | **9.024** | **61.186** | **-40.216** | **24.319** | **-50.691** | **10.093** | **8.616** | **0.0349** |
| **Noradrenergic neurons of the interfascicular and vagal areas** | -22.282 | 31.184 | -4.521 | 41.519 | -36.910 | 17.351 | -35.620 | 19.924 | 3.032 | 0.3867 |
| **Raphe-inferior** | -36.070 | 21.297 | 8.287 | 76.149 | -43.043 | 35.321 | -53.837 | 12.832 | 5.128 | 0.1627 |
| **Raphe-superior** | **-23.062** | **18.142** | **1.693** | **31.519** | **-27.719** | **23.365** | **-35.294** | **15.659** | **7.995** | **0.0461** |
| **Tangential vestibular nucleus** | -14.118 | 32.248 | 16.140 | 61.918 | -31.628 | 19.374 | -39.082 | 17.425 | 5.693 | 0.1275 |
| **Valvula cerebelli** | **-31.076** | **18.141** | **21.900** | **87.023** | **-37.612** | **27.404** | **-51.431** | **9.316** | **10.44** | **0.0152** |

| | | | | | | | | | | |
|---|---|---|---|---|---|---|---|---|---|---|
| **Eyes** | -2.410 | 17.687 | 5.326 | 27.495 | -9.332 | 29.960 | -22.974 | 15.406 | 5.783 | 0.1227 |
| **Olfactory epithelium** | **-0.227** | **15.377** | **46.655** | **47.423** | **-10.910** | **18.278** | **7.551** | **54.323** | **8.111** | **0.0483** |
| **Vagal ganglia** | -7.564 | 22.695 | 17.595 | 73.941 | -21.641 | 14.241 | -11.837 | 30.492 | 4.003 | 0.2611 |
| **Spinal cord** | -44.319 | 26.511 | -1.351 | 55.710 | -46.563 | 22.052 | -43.428 | 27.537 | 3.679 | 0.2983 |
| **Neuropil region** | -41.224 | 31.187 | -1.157 | 55.445 | -49.340 | 23.056 | -47.165 | 22.237 | 4.407 | 0.2207 |

#### Isoeugenol

**Table S4. Mean fluorescence intensity in brain regions of interest (ROIs) measured using light-sheet imaging after a 3-minute exposure to isoeugenol.** Bold denotes ROIs where anaesthetic administration was found to have a statistically significant effect on activity.

| ROI | Control | | 0.10mM | | 0.20mM | | 0.40mM | | Kruskal-Wallis statistic | p-value |
|---|---|---|---|---|---|---|---|---|---|---|
| | Mean | SD | Mean | SD | Mean | SD | Mean | SD | | |
| **Telencephalon** | **-27.081** | **21.552** | **-7.627** | **29.572** | **-30.895** | **20.352** | **-47.622** | **14.277** | **8.467** | **0.0373** |
| **Olfactory Bulbs** | 29.421 | 78.386 | 33.257 | 54.134 | 22.723 | 131.686 | -7.354 | 68.326 | 3.507 | 0.3199 |
| **Anterior commissure** | **15.357** | **97.311** | **-5.736** | **42.832** | **-24.718** | **23.827** | **-60.551** | **24.873** | **7.992** | **0.0462** |
| **Pallium** | -21.406 | 19.529 | -7.081 | 27.602 | -33.889 | 21.214 | -33.090 | 10.881 | 5.873 | 0.1179 |
| **Subpallium** | **-19.633** | **21.562** | **9.172** | **41.226** | **-35.549** | **20.454** | **-40.882** | **23.468** | **11.38** | **0.0099** |
| **Diencephalon** | **12.591** | **40.509** | **-16.289** | **41.414** | **-25.556** | **22.991** | **-47.547** | **26.069** | **11.58** | **0.009** |
| **Dorsal thalamus** | **-3.363** | **26.361** | **-19.050** | **45.295** | **-33.276** | **20.199** | **-54.759** | **20.344** | **11.49** | **0.0094** |
| **Eminentia thalami** | **32.172** | **20.532** | **1.120** | **47.928** | **-27.499** | **23.325** | **-30.990** | **24.734** | **14.21** | **0.0026** |
| **Habenulae** | -7.399 | 25.370 | -12.683 | 23.702 | -31.011 | 13.624 | -28.922 | 21.810 | 5.329 | 0.1492 |
| **Intermediate hypothalamus** | **10.856** | **44.301** | **13.357** | **45.910** | **-28.471** | **38.843** | **-53.002** | **23.091** | **13.48** | **0.0037** |
| **Pineal** | 27.347 | 63.670 | -14.131 | 22.978 | -24.852 | 7.498 | -8.058 | 25.756 | 5.283 | 0.1522 |
| **Posterior tuberculum** | 27.995 | 68.915 | -14.969 | 46.006 | -20.655 | 28.997 | -34.606 | 34.806 | 6.552 | 0.0876 |
| **Preoptic area** | **29.486** | **41.831** | **-22.168** | **40.438** | **-23.606** | **19.832** | **-31.249** | **31.127** | **9.819** | **0.0202** |
| **Pretectum** | **14.310** | **50.527** | **-23.509** | **44.394** | **-41.226** | **22.671** | **-52.299** | **17.760** | **9.932** | **0.0192** |
| **Rostral hypothalamus** | 5.460 | 37.949 | -14.710 | 31.003 | -25.190 | 34.784 | -36.085 | 23.179 | 3.717 | 0.312 |
| **Ventral thalamus** | 7.720 | 38.417 | -15.082 | 49.056 | -23.620 | 25.196 | -39.949 | 31.253 | 5.964 | 0.1134 |
| **Mesencephalon** | **10.493** | **52.892** | **-21.986** | **36.612** | **-37.511** | **28.056** | **-51.431** | **14.188** | **10.78** | **0.013** |
| **Tectum striatum periventiculare** | 18.155 | 54.460 | -21.738 | 39.388 | -35.633 | 25.436 | -26.636 | 21.595 | 7.333 | 0.062 |

| | | | | | | | | | | |
|---|---|---|---|---|---|---|---|---|---|---|
| **Tectum neuropil** | **35.500** | **85.560** | **-20.443** | **34.084** | **-35.736** | **19.916** | **-58.355** | **11.594** | **14.63** | **0.0022** |
| **Tegmentum** | **-5.028** | **57.871** | **-14.296** | **34.247** | **-30.202** | **46.854** | **-67.784** | **14.358** | **14.2** | **0.0026** |
| **Torus longitudinaris** | **70.195** | **79.897** | **-10.261** | **32.904** | **-9.028** | **35.312** | **40.543** | **52.178** | **10.66** | **0.0137** |
| **Torus semicircularis** | **29.529** | **68.819** | **-12.514** | **51.297** | **-41.542** | **21.781** | **-41.054** | **16.860** | **10.63** | **0.0139** |
| **Rhombencephalon** | **-30.406** | **23.667** | **-18.454** | **32.744** | **-37.558** | **46.910** | **-78.583** | **9.460** | **17.02** | **0.0007** |
| **Area postrema** | 21.057 | 65.095 | -9.771 | 36.318 | -10.102 | 47.062 | -37.738 | 48.755 | 4.74 | 0.1919 |
| **Cerebellum** | **-32.904** | **31.324** | **-14.563** | **43.839** | **-44.846** | **36.943** | **-73.061** | **10.702** | **13.22** | **0.0042** |
| **Corpus cerebelli** | **-29.205** | **38.440** | **-16.341** | **38.874** | **-43.068** | **38.101** | **-71.476** | **11.612** | **12.88** | **0.0049** |
| **Eminentia granularis** | **-20.908** | **27.047** | **0.312** | **47.272** | **-34.405** | **44.228** | **-69.827** | **14.577** | **13.85** | **0.0031** |
| **Inferior olive** | **-35.578** | **19.347** | **-16.810** | **35.853** | **-31.172** | **51.660** | **-66.698** | **17.918** | **10.44** | **0.0152** |
| **Interpeduncular nucleus** | -19.895 | 21.574 | -3.347 | 40.492 | -32.770 | 26.525 | -39.516 | 37.041 | 6.187 | 0.1028 |
| **Lateral reticular nucleus** | **29.039** | **79.810** | **-14.700** | **44.299** | **-19.838** | **57.255** | **-74.970** | **17.341** | **15.1** | **0.0017** |
| **Lobus caudalis cerebelli** | **-9.227** | **72.155** | **-26.387** | **49.379** | **-48.809** | **27.399** | **-71.632** | **7.360** | **13.33** | **0.004** |
| **Locus coeruleus** | **-20.340** | **45.909** | **-13.762** | **39.241** | **-27.508** | **64.446** | **-71.339** | **16.910** | **13.17** | **0.0043** |
| **Mauthner** | **-6.570** | **39.273** | **-11.500** | **38.919** | **-40.344** | **40.119** | **-72.599** | **14.809** | **16.18** | **0.001** |
| **Medial vestibular nucleus** | **-28.383** | **29.988** | **-17.285** | **41.803** | **-41.185** | **36.943** | **-75.674** | **10.943** | **13.73** | **0.0033** |
| **Noradrenergic neurons of the interfascicular and vagal areas** | **-15.150** | **31.494** | **-19.554** | **47.536** | **-28.903** | **39.585** | **-64.112** | **16.708** | **11.36** | **0.0099** |
| **Raphe-inferior** | **-3.611** | **63.384** | **-17.471** | **28.378** | **-23.387** | **54.453** | **-66.635** | **7.352** | **13.14** | **0.0043** |
| **Raphe-superior** | -24.012 | 20.871 | -4.361 | 39.426 | -32.688 | 32.883 | -35.929 | 26.530 | 5.625 | 0.1314 |
| **Tangential vestibular nucleus** | **-0.319** | **35.216** | **-5.320** | **42.913** | **-31.722** | **31.887** | **-58.850** | **19.959** | **14.26** | **0.0026** |
| **Valvula cerebelli** | -0.358 | 36.491 | -13.896 | 44.870 | -38.154 | 33.643 | -26.935 | 28.225 | 5.471 | 0.1404 |

| | | | | | | | | | |
|---|---|---|---|---|---|---|---|---|---|
| **Eyes** | 9.197 | 32.434 | 1.023 | 25.100 | -2.454 | 29.170 | -15.267 | 25.206 | 1.848 | 0.6046 |
| **Olfactory epithelium** | 30.993 | 58.880 | 11.567 | 24.992 | -16.233 | 26.781 | 11.367 | 29.294 | 5.91 | 0.1161 |
| **Vagal ganglia** | **234.541** | **131.582** | **16.087** | **37.972** | **36.224** | **19.428** | **81.795** | **62.087** | **13.75** | **0.0033** |
| **Spinal cord** | **-25.861** | **35.315** | **-25.246** | **36.297** | **-27.381** | **49.807** | **-78.987** | **13.679** | **13.31** | **0.004** |
| **Neuropil region** | **-20.380** | **42.617** | **-22.883** | **39.231** | **-21.488** | **53.080** | **-77.816** | **15.436** | **12.59** | **0.0056** |

## MS222

**Table S5. Mean fluorescence intensity in brain regions of interest (ROIs) measured using light-sheet imaging after a 3-minute exposure to MS222.** Bold denotes ROIs where anaesthetic administration was found to have a statistically significant effect on activity.

| ROI | Control | | 0.2mM | | 0.3mM | | 0.35mM | | 0.4mM | | 0.5mM | | 0.7mM | | 1mM | | Kruskal-Wallis statistic | p-value |
|---|---|---|---|---|---|---|---|---|---|---|---|---|---|---|---|---|---|---|
| | Mean | SD | Mean | SD | Mean | SD | Mean | SD | Mean | SD | Mean | SD | Mean | SD | Mean | SD | | |
| Telencephalon | 9.51 | 52.72 | -11.88 | 43.61 | -33.12 | 34.53 | -41.59 | 28.77 | 22.57 | 101.98 | -25.44 | 27.41 | -50.82 | 29.19 | 11.92 | 223.16 | 14 | 0.0512 |
| Olfactory Bulbs | 49.04 | 87.63 | -30.00 | 24.48 | -10.10 | 43.83 | -25.59 | 36.99 | 56.29 | 121.90 | -9.90 | 32.86 | -30.01 | 36.13 | -43.30 | 22.70 | 11.78 | 0.1082 |
| Anterior commissure | 3.45 | 56.28 | -21.75 | 23.31 | -24.72 | 34.58 | -54.40 | 23.81 | 16.32 | 96.55 | -38.98 | 30.61 | -59.15 | 23.89 | 155.44 | 515.35 | 12.76 | 0.0782 |
| Pallium | 14.16 | 58.76 | -30.18 | 26.50 | -31.74 | 30.13 | -41.76 | 27.60 | -24.93 | 46.74 | -24.92 | 31.51 | -46.15 | 26.43 | -22.79 | 151.00 | 14.39 | 0.0446 |
| Subpallium | 0.00 | 39.66 | -36.82 | 35.08 | -29.22 | 29.72 | -26.68 | 28.29 | 144.42 | 340.46 | -31.90 | 22.86 | -48.26 | 28.19 | 38.59 | 293.16 | 13.23 | 0.0668 |
| Diencephalon | -28.39 | 33.27 | -36.17 | 36.75 | -34.10 | 42.09 | -33.90 | 31.43 | -47.38 | 21.60 | -45.79 | 26.08 | -49.11 | 22.69 | -58.22 | 41.09 | 5.619 | 0.5849 |
| Dorsal thalamus | -33.19 | 30.27 | -33.07 | 33.77 | -37.10 | 42.52 | -30.21 | 29.89 | -27.20 | 26.31 | -42.78 | 29.94 | -41.03 | 28.75 | -51.02 | 42.67 | 3.237 | 0.8623 |
| Eminentia thalami | 30.42 | 71.58 | -23.38 | 16.33 | -13.51 | 78.28 | -27.16 | 51.16 | -33.14 | 66.07 | -29.06 | 27.24 | -40.91 | 21.61 | -70.60 | 40.11 | 15.82 | 0.068 |
| Habenulae | -14.62 | 31.04 | -40.62 | 36.22 | -15.72 | 27.21 | -22.60 | 36.29 | -29.18 | 60.05 | -20.99 | 18.34 | -30.58 | 18.58 | -61.86 | 47.11 | 13.8 | 0.0548 |
| Intermediate hypothalamus | -31.12 | 33.24 | -14.90 | 24.20 | -41.72 | 32.74 | -34.14 | 44.77 | -55.95 | 15.09 | -43.34 | 33.91 | -47.80 | 32.24 | -66.02 | 26.59 | 6.098 | 0.5283 |
| Pineal | -19.23 | 28.70 | -35.09 | 44.21 | -18.99 | 26.21 | -7.32 | 36.91 | -2.72 | 45.87 | -20.22 | 21.85 | -31.80 | 17.99 | -29.39 | 78.42 | 5.06 | 0.6527 |
| Posterior tuberculum | -34.28 | 30.77 | -26.45 | 33.47 | -36.99 | 41.75 | -33.19 | 36.57 | -49.55 | 21.13 | -48.49 | 26.35 | -45.77 | 35.52 | -54.88 | 42.45 | 3.692 | 0.8144 |
| Preoptic area | -8.52 | 48.25 | -39.59 | 36.89 | -39.94 | 34.47 | -45.28 | 27.70 | -60.00 | 15.85 | -41.91 | 19.13 | -45.84 | 22.30 | -64.65 | 38.43 | 14.09 | 0.0496 |
| Pretectum | -35.95 | 30.12 | -36.08 | 31.05 | -43.24 | 28.73 | -33.78 | 31.42 | -28.35 | 32.07 | -41.63 | 31.40 | -42.73 | 28.88 | -47.46 | 46.77 | 2.12 | 0.9526 |
| Rostral hypothalamus | -26.31 | 35.90 | -33.12 | 37.39 | -8.77 | 35.88 | -32.10 | 43.49 | -48.37 | 17.17 | -49.63 | 21.23 | -14.91 | 35.38 | -58.04 | 39.07 | 10.51 | 0.1614 |

| | | | | | | | | | | | | | | | | | | |
|---|---|---|---|---|---|---|---|---|---|---|---|---|---|---|---|---|---|---|
| **Ventral thalamus** | -32.67 | 30.30 | -42.56 | 37.82 | -37.21 | 38.95 | -35.09 | 33.98 | -35.54 | 22.89 | -50.81 | 23.30 | -49.25 | 23.43 | -53.04 | 41.56 | 4.211 | 0.7551 |
| **Mesencephalon** | -40.14 | 29.68 | -41.91 | 37.54 | -39.97 | 35.19 | -40.35 | 32.75 | -44.23 | 20.98 | -51.15 | 24.54 | -55.96 | 23.63 | -57.76 | 38.98 | 3.554 | 0.8295 |
| **Tectum striatum periventiculare** | -39.13 | 30.71 | -41.68 | 32.64 | -41.44 | 33.67 | -38.06 | 35.01 | -39.94 | 23.13 | -49.86 | 25.99 | -54.88 | 22.27 | -55.64 | 40.26 | 3.589 | 0.8257 |
| **Tectum neuropil** | -37.12 | 28.86 | -44.18 | 45.33 | -28.00 | 44.46 | -39.11 | 29.61 | -35.83 | 24.24 | -48.31 | 27.59 | -53.03 | 22.11 | -53.58 | 33.99 | 3.552 | 0.8297 |
| **Tegmentum** | -42.29 | 33.91 | -25.78 | 36.38 | -54.14 | 27.53 | -46.04 | 41.08 | -52.00 | 23.17 | -53.80 | 32.72 | -55.16 | 29.03 | -64.75 | 32.12 | 2.512 | 0.9262 |
| **Torus longitudinaris** | -18.87 | 27.41 | -45.23 | 22.74 | -32.67 | 22.87 | -18.33 | 33.68 | 8.94 | 36.58 | -3.59 | 44.33 | -37.84 | 20.99 | -14.45 | 86.74 | 8.662 | 0.2779 |
| **Torus semicircularis** | -22.49 | 33.02 | -47.42 | 41.23 | -22.73 | 50.58 | -17.09 | 42.57 | -32.90 | 26.66 | -53.05 | 22.15 | -61.01 | 17.34 | -38.26 | 78.28 | 12.57 | 0.0833 |
| **Rhombencephalon** | -36.49 | 41.87 | -19.10 | 55.47 | -64.79 | 23.08 | -60.94 | 31.07 | -63.00 | 19.06 | -58.02 | 27.11 | -65.90 | 21.43 | -74.64 | 27.38 | 6.19 | 0.5178 |
| **Area postrema** | -29.42 | 41.69 | -45.15 | 39.49 | -39.63 | 31.91 | -18.51 | 35.17 | -38.55 | 34.07 | -19.00 | 33.92 | -48.90 | 34.38 | -50.32 | 83.58 | 10.86 | 0.145 |
| **Cerebellum** | -38.11 | 37.06 | -42.70 | 40.58 | -58.67 | 24.35 | -55.77 | 30.65 | -55.98 | 18.93 | -56.41 | 21.11 | -59.34 | 19.20 | -69.87 | 30.92 | 4.867 | 0.6761 |
| **Corpus cerebelli** | -37.50 | 37.37 | -42.49 | 42.50 | -57.22 | 26.82 | -52.66 | 34.01 | -55.05 | 21.40 | -53.14 | 25.19 | -57.57 | 20.24 | -71.15 | 28.57 | 5.329 | 0.6199 |
| **Eminentia granularis** | -31.20 | 40.71 | -59.35 | 31.73 | -40.05 | 33.45 | -49.12 | 32.04 | 11.87 | 149.04 | -53.97 | 20.26 | -62.38 | 19.70 | 18.53 | 216.19 | 5.958 | 0.5447 |
| **Inferior olive** | -34.68 | 46.52 | -38.79 | 51.44 | -58.51 | 17.21 | -63.07 | 19.68 | -60.07 | 17.39 | -57.61 | 32.52 | -69.00 | 22.43 | -62.12 | 46.97 | 5.836 | 0.559 |
| **Interpeduncular nucleus** | -25.75 | 44.33 | -46.53 | 19.94 | -40.36 | 34.98 | -40.41 | 39.64 | -49.86 | 24.34 | -46.34 | 19.62 | -45.65 | 22.74 | -66.73 | 29.99 | 5.937 | 0.5471 |
| **Lateral reticular nucleus** | -32.13 | 42.15 | -33.87 | 32.59 | -67.24 | 17.92 | -57.10 | 20.89 | -23.05 | 83.83 | -51.23 | 26.48 | -45.57 | 44.78 | -28.35 | 83.05 | 5.312 | 0.3194 |
| **Lobus caudalis cerebelli** | -42.12 | 33.65 | -46.61 | 29.14 | -55.49 | 20.82 | -58.76 | 25.63 | -56.00 | 16.97 | -54.11 | 18.86 | -57.97 | 15.24 | -67.85 | 32.45 | 8.152 | 0.3194 |
| **Locus coeruleus** | -38.93 | 33.99 | -48.57 | 35.90 | -54.34 | 24.73 | -47.15 | 35.88 | -53.75 | 17.96 | -56.84 | 20.73 | -62.76 | 23.46 | -69.60 | 25.78 | 5.942 | 0.5466 |

| | | | | | | | | | | | | | | | | | | |
|---|---|---|---|---|---|---|---|---|---|---|---|---|---|---|---|---|---|---|
| **Mauthner** | -36.89 | 37.81 | -50.05 | 39.77 | -54.74 | 30.08 | -51.71 | 36.71 | -55.14 | 20.84 | -55.39 | 20.99 | -55.00 | 23.80 | -74.85 | 18.16 | 6.471 | 0.486 |
| **Medial vestibular nucleus** | -36.67 | 38.43 | -39.74 | 29.25 | -63.06 | 23.91 | -57.02 | 33.17 | -63.76 | 22.71 | -59.34 | 20.45 | -66.49 | 21.99 | -80.23 | 17.21 | 9.362 | 0.2277 |
| **Noradrenergic neurons of the interfascicular and vagal areas** | -28.64 | 48.10 | -48.62 | 35.05 | -64.46 | 17.76 | -51.18 | 25.17 | -26.35 | 74.78 | -35.03 | 20.44 | -56.86 | 25.55 | 84.87 | 458.21 | 10.29 | 0.1726 |
| **Raphe-inferior** | -35.37 | 44.98 | -24.47 | 50.15 | -65.70 | 15.41 | -46.01 | 39.28 | -66.48 | 14.47 | -54.28 | 43.97 | -59.86 | 17.38 | -66.52 | 36.38 | 5.678 | 0.5778 |
| **Raphe-superior** | -29.61 | 35.43 | -47.72 | 31.44 | -33.71 | 28.64 | -21.27 | 29.34 | -29.19 | 29.06 | -40.13 | 22.86 | -35.10 | 23.22 | -69.54 | 25.65 | 10.7 | 0.1522 |
| **Tangential vestibular nucleus** | -25.70 | 35.60 | -35.95 | 38.89 | -52.88 | 26.58 | -47.41 | 27.56 | -50.65 | 18.40 | -44.97 | 32.37 | -60.16 | 19.15 | -54.16 | 63.66 | 8.446 | 0.2949 |
| **Valvula cerebelli** | -29.81 | 43.89 | -21.72 | 20.20 | -57.14 | 20.84 | -45.51 | 30.04 | -37.76 | 31.64 | -52.56 | 22.72 | -34.00 | 32.57 | -50.60 | 49.61 | 4.78 | 0.6968 |
| **Eyes** | -1.77 | 16.97 | -25.56 | 18.76 | -3.76 | 55.36 | 15.94 | 83.98 | 16.20 | 28.94 | -32.77 | 23.20 | -23.96 | 28.08 | -18.82 | 64.61 | 13.37 | 0.0637 |
| **Olfactory epithelium** | 5.78 | 27.29 | -19.75 | 27.69 | -6.07 | 29.81 | -8.67 | 27.24 | 20.63 | 61.42 | -11.94 | 8.53 | -27.05 | 25.94 | -39.07 | 31.73 | 13.89 | 0.0532 |
| **Vagal ganglia** | -7.13 | 25.32 | -65.46 | 23.09 | -12.29 | 33.14 | 46.69 | 78.19 | 20.85 | 68.42 | -13.02 | 17.76 | -5.35 | 44.95 | -28.70 | 60.51 | 7.241 | 0.4042 |
| **Spinal cord** | -28.32 | 58.62 | -61.84 | 28.73 | -72.75 | 14.73 | -60.82 | 40.38 | -67.35 | 24.24 | -60.23 | 30.80 | -75.47 | 24.10 | -55.62 | 56.72 | 6.159 | 0.5213 |
| **Neuropil region** | -37.87 | 49.91 | -11.88 | 43.61 | -73.44 | 14.16 | -60.98 | 42.72 | -64.05 | 24.26 | -59.27 | 32.57 | -74.89 | 24.96 | -47.93 | 64.01 | 4.832 | 0.6804 |

#### Quinaldine sulfate

**Table S6. Mean fluorescence intensity in brain regions of interest (ROIs) measured using light-sheet imaging after a 3-minute exposure to quinaldine sulfate.** Bold denotes ROIs where anaesthetic administration was found to have a statistically significant effect on activity.

| ROI | Control | | 0.04mM | | 0.08M | | 0.2mM | | 0.4mM | | Kruskal-Wallis statistic | p-value |
|---|---|---|---|---|---|---|---|---|---|---|---|---|
| | Mean | SD | Mean | SD | Mean | SD | Mean | SD | Mean | SD | | |
| **Telencephalon** | **15.815** | **21.404** | **-5.151** | **19.145** | **-7.611** | **21.648** | **-5.834** | **34.138** | **-46.753** | **38.106** | **13.18** | **0.0104** |
| Olfactory Bulbs | 0.257 | 48.474 | -10.070 | 29.957 | -11.822 | 28.734 | -12.535 | 64.013 | -28.720 | 31.742 | 1.403 | 0.8473 |
| Anterior commissure | 16.372 | 38.819 | -9.740 | 19.703 | -23.633 | 40.963 | -15.768 | 38.246 | -43.784 | 44.396 | 9.121 | 0.0581 |
| **Pallium** | **17.572** | **27.620** | **-2.983** | **21.991** | **-6.806** | **25.598** | **-1.913** | **28.362** | **-52.761** | **36.325** | **13.21** | **0.0103** |
| **Subpallium** | **16.209** | **22.628** | **-6.232** | **21.931** | **-9.699** | **26.148** | **-9.876** | **41.900** | **-50.403** | **40.688** | **11.71** | **0.0196** |
| Diencephalon | 7.076 | 26.245 | -17.969 | 14.106 | -10.160 | 37.951 | -9.958 | 72.859 | -38.711 | 37.272 | 7.705 | 0.132 |
| Dorsal thalamus | 4.140 | 26.977 | -22.514 | 17.113 | -20.537 | 28.651 | -23.139 | 60.256 | -34.709 | 42.362 | 6.598 | 0.1587 |
| **Eminentia thalami** | **8.080** | **26.066** | **6.726** | **28.367** | **4.007** | **24.586** | **11.685** | **70.624** | **-62.971** | **27.595** | **15.24** | **0.0042** |
| Habenulae | 16.537 | 27.821 | -1.408 | 15.223 | 3.454 | 28.600 | 0.644 | 21.078 | -36.159 | 51.782 | 8.914 | 0.0633 |
| Intermediate hypothalamus | 3.162 | 31.356 | -11.944 | 28.145 | -1.455 | 48.356 | -14.721 | 63.024 | -31.325 | 37.419 | 4.171 | 0.3833 |
| Pineal | 4.616 | 28.266 | 1.314 | 25.569 | 1.677 | 20.090 | 9.367 | 36.901 | -36.931 | 40.269 | 6.016 | 0.198 |
| Posterior tuberculum | -1.896 | 29.591 | -22.695 | 15.371 | -8.207 | 39.877 | -13.225 | 93.812 | -24.888 | 45.207 | 3.971 | 0.41 |
| **Preoptic area** | **6.987** | **26.700** | **-16.387** | **24.509** | **-4.443** | **29.978** | **-6.021** | **55.389** | **-58.266** | **26.196** | **14.45** | **0.006** |
| Pretectum | 3.134 | 26.883 | -23.929 | 22.597 | -16.052 | 32.537 | -16.650 | 62.906 | -27.162 | 45.308 | 4.299 | 0.367 |
| Rostral hypothalamus | 5.620 | 23.123 | -20.789 | 18.799 | -14.189 | 32.194 | -11.813 | 84.564 | -23.184 | 43.004 | 6.672 | 0.1543 |
| Ventral thalamus | 2.833 | 27.969 | -22.419 | 20.400 | -9.848 | 37.559 | -9.178 | 92.988 | -25.468 | 53.833 | 3.905 | 0.419 |
| Mesencephalon | 4.908 | 27.916 | -18.822 | 20.164 | -18.931 | 29.593 | -19.041 | 44.679 | -24.716 | 44.493 | 4.293 | 0.3678 |
| Tectum striatum periventiculare | 7.872 | 29.862 | -15.676 | 23.996 | -17.814 | 29.775 | -12.996 | 50.315 | -24.143 | 44.198 | 4.18 | 0.3821 |
| Tectum neuropil | 9.796 | 26.557 | -17.508 | 13.652 | -17.354 | 25.936 | -16.204 | 29.032 | -21.697 | 42.778 | 6.115 | 0.1908 |

| | | | | | | | | | | | | |
|---|---|---|---|---|---|---|---|---|---|---|---|---|
| **Tegmentum** | 4.211 | 37.208 | -23.890 | 23.151 | -12.223 | 47.380 | -22.514 | 55.455 | -24.804 | 43.909 | 3.168 | 0.5302 |
| **Torus longitudinaris** | -0.152 | 21.503 | -9.604 | 22.429 | -9.032 | 23.714 | -4.076 | 41.090 | 11.141 | 74.010 | 2.12 | 0.7136 |
| **Torus semicircularis** | 7.260 | 22.510 | -4.246 | 21.873 | -9.318 | 44.096 | -25.772 | 34.527 | -5.907 | 60.968 | 5.2 | 0.2674 |
| **Rhombencephalon** | -5.318 | 34.541 | -22.599 | 25.449 | -18.466 | 38.048 | -8.672 | 57.511 | -25.112 | 53.986 | 1.507 | 0.8254 |
| **Area postrema** | **-9.231** | **36.034** | **5.952** | **42.074** | **60.078** | **79.978** | **74.812** | **78.882** | **-26.689** | **59.390** | **13.29** | **0.01** |
| **Cerebellum** | -2.662 | 32.493 | -20.072 | 30.645 | -27.741 | 35.043 | -20.232 | 47.516 | -31.776 | 41.684 | 2.826 | 0.5874 |
| **Corpus cerebelli** | -0.845 | 33.925 | -19.432 | 32.167 | -22.742 | 40.208 | -17.426 | 48.519 | -30.585 | 41.889 | 2.495 | 0.6454 |
| **Eminentia granularis** | -2.475 | 28.269 | -19.106 | 24.508 | -32.243 | 31.392 | -26.979 | 42.324 | 84.466 | 313.367 | 5.852 | 0.2105 |
| **Inferior olive** | -6.878 | 38.356 | -27.785 | 24.366 | -9.245 | 47.432 | -9.690 | 72.475 | -42.905 | 42.029 | 4.825 | 0.3058 |
| **Interpeduncular nucleus** | 2.369 | 24.982 | -10.500 | 17.308 | -6.368 | 36.814 | -18.548 | 29.314 | -45.540 | 33.210 | 8.644 | 0.0707 |
| **Lateral reticular nucleus** | **-11.201** | **31.090** | **11.743** | **55.411** | **55.506** | **50.419** | **79.780** | **57.900** | **-25.115** | **62.530** | **16.27** | **0.0027** |
| **Lobus caudalis cerebelli** | 1.831 | 32.010 | -18.202 | 32.204 | -32.457 | 18.846 | -31.668 | 25.301 | -17.310 | 62.382 | 5.798 | 0.2147 |
| **Locus coeruleus** | -1.730 | 31.648 | -17.787 | 22.670 | -20.847 | 26.584 | -26.794 | 35.650 | -24.211 | 51.487 | 4.013 | 0.4043 |
| **Mauthner** | 7.416 | 25.866 | -20.039 | 22.930 | -18.234 | 35.042 | -24.994 | 47.727 | -10.110 | 67.167 | 6.152 | 0.1881 |
| **Medial vestibular nucleus** | -4.216 | 28.583 | -17.875 | 24.951 | -16.176 | 41.333 | -19.739 | 56.910 | -24.002 | 57.150 | 2.514 | 0.6421 |
| **Noradrenergic neurons of the interfascicular and vagal areas** | **-16.459** | **28.191** | **5.380** | **53.636** | **41.853** | **65.548** | **56.462** | **40.200** | **14.060** | **130.227** | **10.26** | **0.0363** |
| **Raphe-inferior** | -9.922 | 37.872 | -29.359 | 30.500 | -0.235 | 58.805 | -19.629 | 60.974 | -8.184 | 87.694 | 1.485 | 0.8293 |
| **Raphe-superior** | 6.381 | 38.481 | -4.308 | 36.715 | -11.188 | 53.958 | -25.828 | 35.273 | -43.831 | 30.980 | 8.099 | 0.088 |
| **Tangential vestibular nucleus** | -5.226 | 16.437 | -7.767 | 24.011 | -13.729 | 26.027 | -12.852 | 41.764 | -21.024 | 56.822 | 1.622 | 0.8048 |
| **Valvula cerebelli** | 0.412 | 37.319 | -21.653 | 24.700 | -12.250 | 54.437 | -14.054 | 63.790 | -18.680 | 50.864 | 2.489 | 0.6467 |
| **Eyes** | 15.901 | 37.912 | -10.807 | 21.192 | -10.829 | 28.350 | -5.223 | 30.646 | -28.060 | 36.269 | 5.318 | 0.2562 |

| | | | | | | | | | | | |
|---|---|---|---|---|---|---|---|---|---|---|---|
| **Olfactory epithelium** | 5.642 | 22.734 | -0.891 | 21.440 | -10.004 | 18.077 | 0.504 | 34.855 | -34.132 | 32.946 | 6.402 | 0.1711 |
| **Vagal ganglia** | **-3.064** | **17.726** | **12.838** | **45.351** | **25.895** | **41.964** | **78.851** | **25.324** | **-18.316** | **41.277** | **16.14** | **0.0028** |
| **Spinal cord** | 15.815 | 56.390 | -14.976 | 33.652 | 21.841 | 66.227 | -0.922 | 43.260 | -28.260 | 73.124 | 4.605 | 0.3303 |
| **Neuropil region** | 15.418 | 60.153 | -10.531 | 43.660 | 20.592 | 65.318 | -3.279 | 45.599 | -23.329 | 78.241 | 3.579 | 0.4659 |

**In vivo $Ca^{2+}$ functional brain imaging in zebrafish embryo-larvae exposed to the test anaesthesia with the application of a visual stimulus**

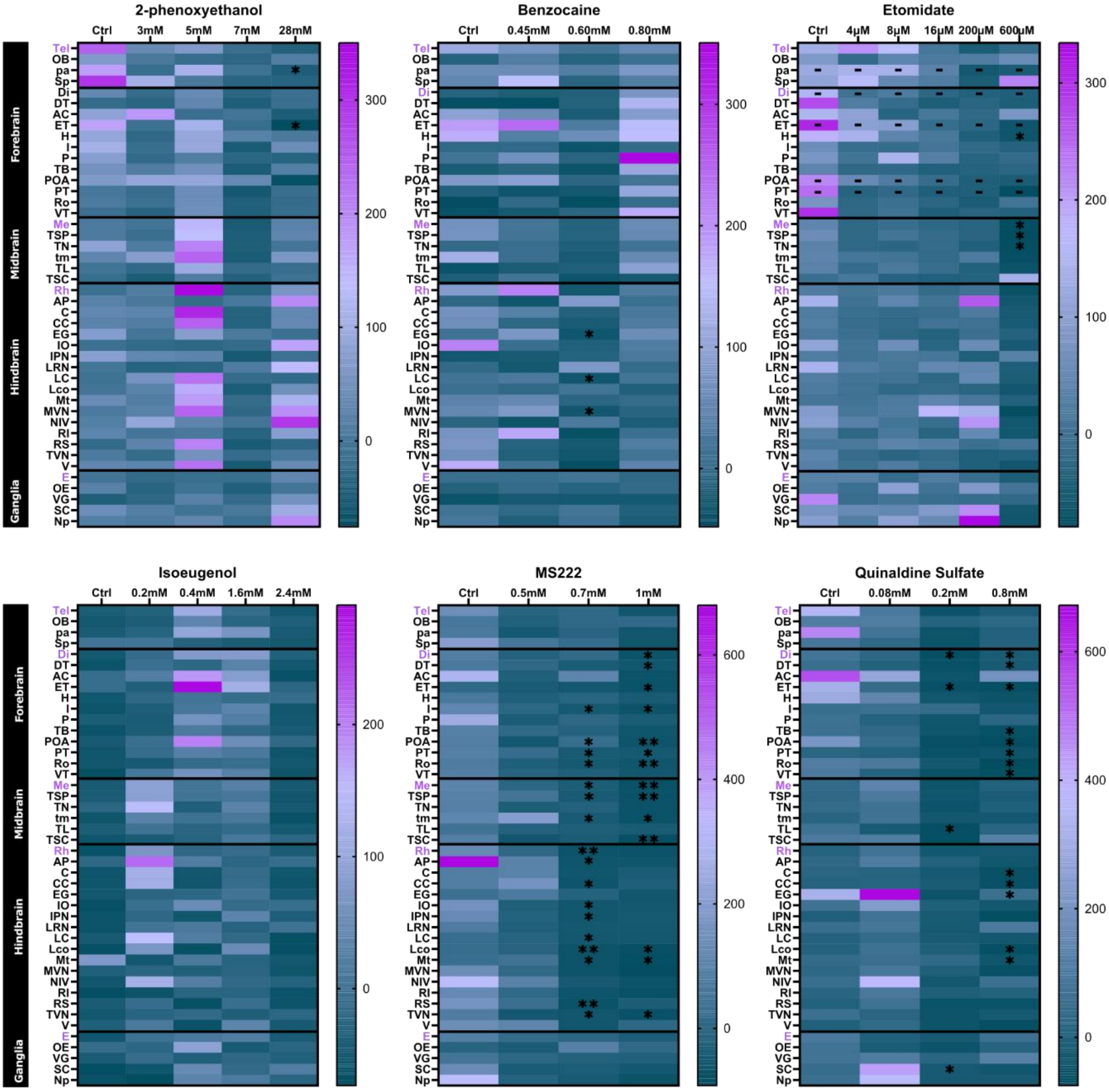


**Figure S1. Heat maps showing mean brain activity (as measured by fluorescence intensity) in brain regions of interest (ROI) in embryo-larvae exposed to the tested anaesthetics, with the application of a light stimulus.** Data are presented as the average baseline (pre-exposure) corrected mean neural activity (fluorescence intensity), calculated using the equation: $\Delta f/f = (f_1 - f_0)/f_0 * 100$ (where $f_1$ = mean fluorescence intensity and $f_0$ = baseline fluorescence intensity for the same embryo larva within each ROI) (n = 7–9 larvae per group). Overall, anaesthetic administration led to a decrease in brain activity (fluorescence intensity) associated with light stimulation at the higher test concentrations, although this effect was not as pronounced as for the case without light stimulation (**Figure 2**). Statistically significant differences compared with control animals are presented as *p<0.05 and **p<0.01. Cases where an overall difference across treatments was detected but there were no statistically significant single-dose differences to the control condition following *post hoc* testing are represented by – in the figure. Abbreviations used for the brain regions assessed are: (Tel) telencephalon, (OB) olfactory bulb, (AC) anterior commissure, (pa) pallium, (Sb) subpallium, (Di) diencephalon, (ET) thalamic eminence, (H) habenuale, (I) intermediate hypothalamus, (P) pineal, (TB) posterior tuberculum, (POA) preoptic area, (PT) pretectum, (Ro) rostral hypothalamus, (VT) ventral thalamus, (Me) mesencephalon, (TSP) tectum stratum periventiculare, (TN) tectum neuropil, (tm) tegmentum, (TL) torus longitudinaris, (TSC), torus semicircularis, (Rh) rhombencephalon, (AP) area postrema, (C) cerebellum, (CC) corpus cerebelli, (EG) granular eminence, (IO) inferior olive, (IPN) interpeduncular nucleus, (LRN) lateral reticular nucleus, (LC) lobus caudalis cerebelli, (LCo) locus coeruleus, (Mt) Mauthner, (MVN) medial vestibular nucleus, (NIV) noradrenergic neurons of the interfascicular and vagal areas, (RI) inferior raphe nucleus, (RS) superior raphe nucleus, (TVN) tangential vestibular nucleus, (V) valvula cerebelli, (E) eyes, (OE) olfactory epithelium, (VG) vagal ganglia, (SC) spinal cord, (Np) neuropil region.

#### 2-phenoxyethanol

Table S7. **Mean fluorescence intensity in brain regions of interest (ROIs) measured using light-sheet imaging during the first 8 seconds post application of a light stimulus, after a 3-minute exposure to 2-phenoxyethanol.** Bold denotes ROIs where anaesthetic administration was found to have a statistically significant effect on activity.

| ROI | Control | | 3mM | | 5mM | | 7mM | | 28mM | | Kruskal-Wallis statistic | p-value |
|---|---|---|---|---|---|---|---|---|---|---|---|---|
| | Mean | SD | Mean | SD | Mean | SD | Mean | SD | Mean | SD | | |
| Telencephalon | 245.768 | 301.859 | 38.139 | 183.505 | 65.616 | 167.715 | -14.656 | 61.242 | -40.317 | 50.099 | 10.15 | 0.0712 |
| Olfactory Bulbs | 67.533 | 117.276 | 3.802 | 108.708 | -16.684 | 65.255 | -29.099 | 53.674 | 12.337 | 92.261 | 4.221 | 0.518 |
| **Anterior commissure** | **81.781** | **140.922** | **182.939** | **300.947** | **-14.257** | **72.812** | **-3.816** | **85.377** | **-47.043** | **83.234** | **12.15** | **0.0328** |
| **Pallium** | **173.715** | **183.881** | **-15.974** | **65.403** | **111.135** | **254.255** | **-8.316** | **65.111** | **-45.863** | **64.443** | **14.21** | **0.0143** |
| Subpallium | 296.032 | 548.522 | 107.452 | 433.869 | 3.663 | 76.240 | -31.804 | 63.709 | -35.265 | 65.446 | 10.01 | 0.0751 |
| Diencephalon | 33.990 | 91.094 | -19.257 | 76.632 | 37.371 | 114.802 | -29.648 | 51.152 | -13.333 | 111.174 | 5.454 | 0.363 |
| Dorsal thalamus | -21.044 | 61.088 | -31.754 | 51.923 | 30.159 | 98.142 | -44.194 | 45.269 | -14.609 | 74.753 | 6.696 | 0.2443 |
| **Eminentia thalami** | **170.220** | **242.330** | **-11.107** | **68.219** | **109.196** | **271.904** | **-15.035** | **84.889** | **-75.626** | **12.835** | **12.74** | **0.0259** |
| Habenulae | 87.552 | 152.585 | -26.773 | 45.755 | 90.033 | 206.989 | 20.080 | 113.300 | 5.178 | 90.773 | 3.745 | 0.5867 |
| Intermediate hypothalamus | 106.964 | 303.758 | -15.927 | 75.312 | 91.243 | 227.322 | -49.152 | 31.471 | 42.379 | 234.550 | 5.183 | 0.394 |
| Pineal | 73.081 | 248.127 | -22.988 | 71.404 | 12.768 | 124.167 | -31.143 | 57.072 | -35.376 | 49.496 | 4.707 | 0.4527 |
| Posterior tuberculum | 51.381 | 185.011 | 11.255 | 142.442 | 49.456 | 98.628 | -47.807 | 46.146 | 16.350 | 125.794 | 6.045 | 0.3019 |
| Preoptic area | 65.076 | 131.169 | 88.424 | 222.548 | 88.795 | 272.593 | 35.144 | 192.951 | -71.189 | 27.201 | 9.447 | 0.0925 |
| Pretectum | 0.215 | 86.597 | -17.394 | 62.815 | 41.841 | 145.101 | -56.421 | 21.617 | -14.132 | 78.115 | 6.779 | 0.2376 |
| Rostral hypothalamus | 7.029 | 100.101 | -11.626 | 83.752 | 43.217 | 97.794 | -41.123 | 39.218 | -20.614 | 70.029 | 7.664 | 0.177 |
| Ventral thalamus | -11.265 | 80.044 | -34.795 | 35.871 | 46.520 | 139.957 | -43.314 | 51.403 | -17.097 | 78.737 | 5.392 | 0.3699 |
| Mesencephalon | 7.299 | 132.120 | -4.231 | 49.219 | 147.761 | 448.344 | -49.488 | 28.087 | 22.320 | 167.302 | 8.305 | 0.1402 |
| Tectum striatum periventiculare | 6.646 | 128.029 | -14.689 | 44.409 | 136.550 | 427.860 | -46.031 | 30.265 | 28.390 | 172.258 | 7.094 | 0.2137 |

| | | | | | | | | | | | | |
|---|---|---|---|---|---|---|---|---|---|---|---|---|
| **Tectum neuropil** | 81.519 | 293.016 | 9.831 | 77.079 | 206.958 | 637.369 | -44.620 | 38.355 | -1.950 | 87.427 | 7.23 | 0.2041 |
| **Tegmentum** | 27.977 | 171.100 | 71.210 | 223.481 | 243.128 | 551.137 | -49.271 | 21.597 | 59.210 | 323.266 | 9.345 | 0.0961 |
| **Torus longitudinaris** | -5.980 | 92.591 | -23.800 | 95.101 | 98.010 | 300.018 | -22.620 | 50.674 | -14.928 | 69.345 | 4.82 | 0.4383 |
| **Torus semicircularis** | -14.983 | 83.221 | -46.634 | 24.130 | 11.457 | 114.804 | -38.203 | 51.352 | -9.900 | 84.277 | 7.884 | 0.1627 |
| **Rhombencephalon** | -11.134 | 117.981 | 13.339 | 82.646 | 350.253 | 923.390 | -33.443 | 29.060 | 55.972 | 314.550 | 6.332 | 0.2752 |
| **Area postrema** | 21.124 | 99.292 | 2.396 | 144.385 | -19.277 | 76.992 | 2.019 | 62.579 | 199.556 | 454.009 | 3.819 | 0.5757 |
| **Cerebellum** | 31.434 | 163.348 | 25.714 | 99.767 | 313.770 | 739.876 | -36.919 | 32.088 | 32.315 | 258.318 | 5.111 | 0.4025 |
| **Corpus cerebelli** | 24.088 | 145.827 | 13.459 | 76.707 | 246.656 | 579.789 | -36.890 | 31.372 | 34.647 | 264.929 | 5.609 | 0.3461 |
| **Eminentia granularis** | 71.948 | 207.437 | -1.965 | 86.518 | 72.171 | 213.762 | 4.980 | 85.220 | -10.881 | 92.080 | 1.644 | 0.8959 |
| **Inferior olive** | -10.932 | 105.012 | -23.187 | 68.021 | -28.753 | 58.328 | -20.754 | 42.578 | 168.927 | 521.152 | 4.463 | 0.4848 |
| **Interpeduncular nucleus** | 74.196 | 236.515 | 30.783 | 152.709 | 24.787 | 135.881 | -53.286 | 25.662 | -9.317 | 72.380 | 3.109 | 0.6831 |
| **Lateral reticular nucleus** | -2.533 | 88.783 | -35.815 | 45.920 | -31.852 | 52.802 | 9.240 | 76.483 | 142.168 | 279.107 | 6.565 | 0.2551 |
| **Lobus caudalis cerebelli** | -8.288 | 84.685 | 53.346 | 102.460 | 225.045 | 507.838 | -19.947 | 49.450 | -22.850 | 127.826 | 6.108 | 0.2958 |
| **Locus coeruleus** | -3.091 | 91.739 | 22.373 | 99.987 | 160.458 | 379.396 | -47.257 | 30.829 | 46.351 | 279.211 | 7.588 | 0.1804 |
| **Mauthner** | 40.063 | 136.973 | 12.522 | 100.175 | 175.550 | 436.882 | -20.343 | 46.620 | 112.437 | 402.814 | 3.057 | 0.6911 |
| **Medial vestibular nucleus** | 5.916 | 118.599 | 19.940 | 116.446 | 242.157 | 722.085 | -25.865 | 36.954 | 192.990 | 606.674 | 3.644 | 0.6017 |
| **Noradrenergic neurons of the interfascicular and vagal areas** | 16.039 | 81.773 | 88.723 | 332.332 | 25.760 | 146.927 | -3.154 | 58.287 | 293.853 | 801.291 | 3.057 | 0.6911 |
| **Raphe-inferior** | 26.650 | 159.735 | -1.418 | 96.224 | 4.475 | 116.237 | -34.179 | 33.905 | 74.694 | 332.302 | 1.525 | 0.9101 |
| **Raphe-superior** | -22.335 | 80.243 | 24.715 | 87.431 | 206.546 | 427.412 | -49.852 | 25.853 | -27.812 | 85.256 | 6.719 | 0.2424 |
| **Tangential vestibular nucleus** | 6.881 | 128.876 | -20.934 | 65.960 | 50.288 | 189.883 | -19.739 | 49.438 | 2.096 | 148.699 | 3.58 | 0.6113 |
| **Valvula cerebelli** | 31.317 | 113.740 | 24.332 | 125.689 | 227.796 | 518.525 | -51.868 | 32.810 | 36.840 | 278.392 | 6.438 | 0.2659 |

| | | | | | | | | | | | | |
|---|---|---|---|---|---|---|---|---|---|---|---|---|
| **Eyes** | -3.418 | 67.797 | -22.409 | 49.951 | -13.822 | 55.480 | -15.707 | 50.658 | 28.309 | 134.203 | 1.011 | 0.9617 |
| **Olfactory epithelium** | 21.431 | 71.068 | -32.960 | 38.099 | -7.987 | 62.019 | -41.005 | 49.746 | -3.070 | 91.692 | 7.272 | 0.2012 |
| **Vagal ganglia** | -44.631 | 55.489 | -28.075 | 69.903 | 20.884 | 116.952 | -20.312 | 39.427 | 49.999 | 100.588 | 7.272 | 0.2012 |
| **Spinal cord** | 43.385 | 131.455 | 15.180 | 120.128 | -7.301 | 94.093 | 3.753 | 72.045 | 98.956 | 362.427 | 3.615 | 0.6061 |
| **Neuropil region** | 16.565 | 104.507 | 7.169 | 116.381 | 29.283 | 160.379 | -11.114 | 56.677 | 198.151 | 497.380 | 2.541 | 0.7703 |

#### Benzocaine

Table S8. **Mean fluorescence intensity in brain regions of interest (ROIs) measured using light-sheet imaging during the first 8 seconds post application of a light stimulus, after a 3-minute exposure to benzocaine.** Bold denotes ROIs where anaesthetic administration was found to have a statistically significant effect on activity.

| ROI | Control | | 0.45mM | | 0.60mM | | 0.80mM | | Kruskal-Wallis statistic | p-value |
|---|---|---|---|---|---|---|---|---|---|---|
| | Mean | SD | Mean | SD | Mean | SD | Mean | SD | | |
| **Telencephalon** | 110.679 | 153.753 | 72.089 | 123.346 | 19.600 | 96.542 | 56.049 | 101.688 | 2.517 | 0.6416 |
| **Olfactory Bulbs** | -11.632 | 42.931 | 35.281 | 64.572 | -10.947 | 79.615 | 11.015 | 48.940 | 4.691 | 0.3204 |
| **Anterior commissure** | **95.097** | **135.192** | **60.722** | **260.668** | **-23.207** | **65.944** | **87.392** | **143.697** | **10.55** | **0.0322** |
| **Pallium** | 60.700 | 104.843 | 59.096 | 99.208 | 36.196 | 112.780 | 79.672 | 171.470 | 1.066 | 0.8996 |
| **Subpallium** | 51.217 | 96.316 | 148.367 | 412.603 | -8.589 | 104.588 | 33.754 | 93.599 | 6.243 | 0.1817 |
| **Diencephalon** | -9.554 | 81.108 | -20.563 | 60.223 | -15.550 | 67.525 | 75.230 | 216.568 | 4.778 | 0.3108 |
| **Dorsal thalamus** | -45.274 | 37.742 | -46.643 | 35.168 | -13.259 | 61.192 | 130.473 | 366.518 | 7.312 | 0.1203 |
| **Eminentia thalami** | 186.607 | 375.990 | 242.310 | 787.017 | 30.396 | 95.155 | 145.356 | 299.420 | 3.37 | 0.498 |
| **Habenulae** | 169.214 | 223.835 | 37.218 | 100.071 | 61.960 | 160.411 | 150.568 | 343.161 | 2.133 | 0.7113 |
| **Intermediate hypothalamus** | -0.990 | 204.008 | 0.870 | 107.635 | -37.637 | 44.148 | 3.277 | 101.748 | 6.243 | 0.1817 |
| **Pineal** | 16.283 | 68.285 | 67.733 | 127.452 | -8.749 | 70.563 | 350.599 | 1021.515 | 3.96 | 0.4114 |
| **Posterior tuberculum** | -22.691 | 97.828 | -7.856 | 77.070 | -38.073 | 49.253 | 108.626 | 322.236 | 6.062 | 0.1946 |
| **Preoptic area** | 82.308 | 155.336 | 95.206 | 302.364 | -4.954 | 110.388 | 16.824 | 69.148 | 6.986 | 0.1366 |
| **Pretectum** | -16.176 | 102.234 | -36.022 | 41.378 | -30.208 | 50.411 | 104.830 | 306.680 | 5.137 | 0.2736 |
| **Rostral hypothalamus** | -48.084 | 35.157 | 0.013 | 69.402 | -47.120 | 35.585 | 4.428 | 127.577 | 6.397 | 0.1714 |
| **Ventral thalamus** | -41.264 | 37.744 | -27.571 | 92.331 | -7.139 | 71.442 | 171.750 | 544.724 | 7.072 | 0.1321 |
| **Mesencephalon** | 69.987 | 326.762 | 20.577 | 143.831 | -12.231 | 86.743 | 58.066 | 208.232 | 2.938 | 0.5683 |
| **Tectum striatum periventiculare** | 72.014 | 328.753 | 17.374 | 137.134 | -11.187 | 84.826 | 52.877 | 196.296 | 2.808 | 0.5905 |

| | | | | | | | | | | |
|---|---|---|---|---|---|---|---|---|---|---|
| **Tectum neuropil** | 0.547 | 145.122 | 9.362 | 80.165 | -16.985 | 62.007 | 15.528 | 103.879 | 1.531 | 0.8211 |
| **Tegmentum** | 121.589 | 524.463 | 8.956 | 109.508 | -26.045 | 65.659 | 55.899 | 206.468 | 6.408 | 0.1707 |
| **Torus longitudinaris** | -39.800 | 41.176 | -22.273 | 58.376 | -17.663 | 49.914 | 93.763 | 357.427 | 3.53 | 0.4733 |
| **Torus semicircularis** | -19.416 | 63.498 | 19.369 | 98.776 | -36.763 | 25.122 | -29.023 | 45.816 | 1.484 | 0.8296 |
| **Rhombencephalon** | 87.632 | 445.639 | 215.256 | 629.646 | -30.217 | 74.712 | 31.637 | 171.135 | 8.965 | 0.062 |
| **Area postrema** | -3.588 | 101.477 | -35.311 | 45.851 | 85.163 | 206.182 | 10.088 | 89.240 | 4.805 | 0.3078 |
| **Cerebellum** | 69.153 | 376.482 | 40.205 | 259.005 | -14.525 | 89.894 | 26.340 | 189.716 | 5.876 | 0.2096 |
| **Corpus cerebelli** | 65.147 | 368.764 | -3.034 | 149.636 | -24.138 | 66.474 | 30.738 | 201.561 | 6.091 | 0.1924 |
| **Eminentia granularis** | **14.998** | **158.501** | **78.307** | **353.849** | **-35.257** | **46.916** | **60.033** | **163.927** | **10.49** | **0.0329** |
| **Inferior olive** | 217.161 | 722.917 | 8.411 | 99.051 | -16.582 | 81.421 | 32.812 | 163.693 | 1.612 | 0.8067 |
| **Interpeduncular nucleus** | -30.531 | 59.065 | -26.569 | 83.504 | -12.282 | 77.754 | 25.530 | 158.512 | 6.604 | 0.1584 |
| **Lateral reticular nucleus** | 0.196 | 166.501 | -30.172 | 78.464 | 80.377 | 237.864 | 2.486 | 73.997 | 5.925 | 0.2048 |
| **Lobus caudalis cerebelli** | **17.632** | **165.880** | **27.358** | **138.616** | **-32.638** | **74.395** | **18.036** | **201.000** | **9.541** | **0.0489** |
| **Locus coeruleus** | 10.166 | 154.432 | -0.991 | 145.673 | 17.283 | 169.836 | -17.930 | 89.712 | 6.516 | 0.1638 |
| **Mauthner** | 44.549 | 286.125 | 54.234 | 236.761 | -6.756 | 134.688 | 2.624 | 117.538 | 7.535 | 0.1102 |
| **Medial vestibular nucleus** | 52.170 | 313.927 | 81.267 | 274.424 | -31.010 | 78.121 | -8.973 | 98.999 | 9.818 | 0.0436 |
| **Noradrenergic neurons of the interfascicular and vagal areas** | -7.491 | 87.193 | -36.826 | 41.143 | 28.214 | 103.380 | -20.992 | 47.395 | 8.354 | 0.0794 |
| **Raphe-inferior** | 72.815 | 343.171 | 176.108 | 391.711 | -42.305 | 57.730 | 10.311 | 160.068 | 6.024 | 0.1974 |
| **Raphe-superior** | 78.309 | 320.952 | -10.466 | 110.445 | -28.333 | 63.101 | 31.868 | 199.506 | 5.588 | 0.2321 |
| **Tangential vestibular nucleus** | 65.163 | 324.564 | -16.796 | 102.637 | -32.551 | 62.123 | 16.748 | 172.279 | 8.127 | 0.087 |
| **Valvula cerebelli** | 170.465 | 657.768 | 15.588 | 141.014 | -44.623 | 42.410 | 45.590 | 238.073 | 6.627 | 0.1569 |

| | | | | | | | | | | |
|---|---|---|---|---|---|---|---|---|---|---|
| **Eyes** | 8.811 | 94.212 | 8.327 | 101.631 | 17.717 | 80.709 | -2.080 | 68.193 | 0.9018 | 0.9243 |
| **Olfactory epithelium** | -13.283 | 47.465 | 2.615 | 49.972 | -10.683 | 64.041 | 0.238 | 45.660 | 1.371 | 0.8492 |
| **Vagal ganglia** | -39.062 | 38.961 | -28.132 | 104.392 | -29.691 | 60.149 | -28.829 | 59.496 | 4.465 | 0.3467 |
| **Spinal cord** | 16.163 | 119.539 | -0.706 | 118.888 | 18.763 | 95.450 | -15.578 | 73.057 | 3.396 | 0.494 |
| **Neuropil region** | 23.402 | 162.416 | -17.326 | 55.600 | 14.756 | 110.043 | -21.658 | 57.685 | 2.539 | 0.6377 |

#### Etomidate

Table S9. **Mean fluorescence intensity in brain regions of interest (ROIs) measured using light-sheet imaging during the first 8 seconds post application of a light stimulus, after a 3-minute exposure to etomidate.** Bold denotes ROIs where anaesthetic administration was found to have a statistically significant effect on activity.

| ROI | Control | | 0.004mM | | 0.008mM | | 0.016mM | | 0.200mM | | 0.600mM | | Kruskal-Wallis statistic | p-value |
|---|---|---|---|---|---|---|---|---|---|---|---|---|---|---|
| | Mean | SD | Mean | SD | Mean | SD | Mean | SD | Mean | SD | Mean | SD | | |
| **Telencephalon** | 131.972 | 150.313 | 209.098 | 456.208 | 165.654 | 429.584 | 9.200 | 136.036 | -48.154 | 44.590 | -11.280 | 118.372 | 12.28 | 0.0560 |
| **Olfactory Bulbs** | 83.477 | 112.674 | -15.778 | 55.675 | 39.322 | 56.446 | 17.248 | 71.591 | 22.057 | 67.933 | 56.360 | 154.634 | 6.364 | 0.3837 |
| **Anterior commissure** | 146.044 | 134.204 | 104.875 | 153.653 | -30.586 | 61.834 | 10.642 | 100.447 | -20.009 | 69.293 | 76.043 | 372.168 | 12.26 | 0.0564 |
| **Pallium** | **121.559** | **89.762** | **138.624** | **299.618** | **135.919** | **362.833** | **37.053** | **127.901** | **-57.249** | **39.471** | **-46.626** | **42.356** | **17.97** | **0.0063** |
| **Subpallium** | 108.477 | 209.198 | 159.779 | 246.191 | 47.189 | 172.765 | 22.112 | 159.115 | -45.658 | 57.083 | 222.980 | 737.486 | 9.869 | 0.1303 |
| **Diencephalon** | **147.296** | **194.777** | **1.414** | **66.090** | **21.224** | **89.154** | **-23.894** | **87.895** | **-37.267** | **60.726** | **-40.047** | **65.775** | **13.05** | **0.0423** |
| **Dorsal thalamus** | 272.835 | 659.870 | 23.435 | 136.669 | -12.300 | 53.650 | -43.894 | 43.605 | -28.175 | 68.821 | -43.840 | 71.484 | 9.443 | 0.1501 |
| **Eminentia thalami** | **306.334** | **388.110** | **99.571** | **156.713** | **94.561** | **340.920** | **-11.101** | **85.422** | **-26.352** | **113.213** | **-64.045** | **28.559** | **19.66** | **0.0032** |
| **Habenulae** | **181.905** | **321.018** | **138.481** | **205.243** | **39.711** | **148.347** | **12.006** | **102.619** | **-40.910** | **46.609** | **-64.600** | **26.336** | **14.9** | **0.0211** |
| **Intermediate hypothalamus** | 59.045 | 107.940 | -8.391 | 97.683 | -12.258 | 53.522 | 7.096 | 135.516 | -19.889 | 124.655 | -34.811 | 42.958 | 5.527 | 0.4782 |
| **Pineal** | 77.438 | 143.213 | 3.329 | 93.929 | 142.538 | 368.752 | 27.944 | 113.986 | -30.989 | 46.081 | -15.794 | 110.515 | 8.931 | 0.1775 |
| **Posterior tuberculum** | 40.439 | 103.895 | -20.949 | 73.257 | -20.451 | 44.167 | -38.662 | 65.201 | -34.666 | 58.863 | -27.349 | 65.230 | 5.498 | 0.4818 |
| **Preoptic area** | **218.221** | **212.428** | **54.002** | **160.502** | **21.342** | **96.526** | **-12.113** | **93.438** | **-40.387** | **42.577** | -44.951 | 39.360 | **16.7** | **0.0104** |
| **Pretectum** | **241.172** | **674.935** | **-32.906** | **47.858** | **-19.178** | **66.402** | **-40.277** | **48.950** | **-41.933** | **55.773** | **-78.375** | **26.536** | **13.33** | **0.0381** |
| **Rostral hypothalamus** | 75.418 | 190.910 | -23.934 | 70.903 | -5.708 | 46.769 | -27.426 | 64.131 | -47.020 | 30.210 | 1.763 | 194.897 | 8.365 | 0.2126 |
| **Ventral thalamus** | 297.537 | 704.855 | -21.213 | 48.690 | -0.235 | 76.579 | -42.796 | 57.681 | -30.002 | 89.395 | -36.387 | 78.300 | 10.63 | 0.1005 |
| **Mesencephalon** | **42.571** | **113.149** | **-21.686** | **71.105** | **-9.306** | **76.241** | **-23.943** | **80.299** | **-29.365** | **99.732** | **-78.252** | **23.179** | **13.63** | **0.0340** |
| **Tectum striatum periventiculare** | **57.510** | **114.624** | **-21.420** | **69.715** | **4.243** | **109.531** | **-15.521** | **83.506** | **-29.869** | **98.161** | **-79.141** | **22.289** | **15.32** | **0.0179** |

| | | | | | | | | | | | | | |
|---|---|---|---|---|---|---|---|---|---|---|---|---|---|---|
| **Tectum neuropil** | **37.331** | **109.012** | **-21.529** | **69.190** | **-26.762** | **67.149** | **-15.007** | **86.803** | **-39.500** | **63.029** | **-77.268** | **22.802** | **14.44** | **0.0251** |
| **Tegmentum** | 39.517 | 124.061 | 8.110 | 92.321 | 3.669 | 93.448 | -15.921 | 96.286 | 17.422 | 194.888 | -75.174 | 19.563 | 10.47 | 0.1064 |
| **Torus longitudinaris** | 35.835 | 149.637 | -12.067 | 86.544 | 14.942 | 55.505 | -10.175 | 72.455 | -37.238 | 50.462 | -67.748 | 34.232 | 12.38 | 0.0539 |
| **Torus semicircularis** | -7.377 | 62.780 | 11.122 | 97.720 | -31.135 | 46.241 | -38.575 | 61.244 | -33.451 | 60.988 | 135.008 | 535.466 | 4.074 | 0.6670 |
| **Rhombencephalon** | 23.515 | 105.898 | 6.676 | 93.455 | 15.239 | 145.962 | -15.378 | 92.925 | 1.507 | 148.202 | -70.124 | 26.584 | 7.217 | 0.3013 |
| **Area postrema** | 137.402 | 253.885 | -1.229 | 118.690 | 42.051 | 148.781 | 3.426 | 75.235 | 259.906 | 833.334 | -59.319 | 43.842 | 7.552 | 0.2728 |
| **Cerebellum** | 20.625 | 94.015 | -6.745 | 91.403 | -13.825 | 86.647 | -8.818 | 90.775 | 18.487 | 139.471 | -52.710 | 63.131 | 5.226 | 0.5152 |
| **Corpus cerebelli** | 25.989 | 99.967 | -9.859 | 86.326 | -19.215 | 80.820 | 1.436 | 104.083 | -1.128 | 130.913 | -52.387 | 64.036 | 6.138 | 0.4079 |
| **Eminentia granularis** | 14.053 | 65.931 | -6.307 | 78.247 | -24.197 | 91.420 | -35.405 | 58.903 | -6.527 | 123.434 | -36.247 | 45.002 | 3.948 | 0.6838 |
| **Inferior olive** | 109.116 | 279.744 | 3.548 | 94.794 | 61.682 | 298.007 | -28.080 | 59.369 | 56.595 | 264.600 | -51.787 | 33.621 | 5.348 | 0.5000 |
| **Interpeduncular nucleus** | 22.457 | 107.499 | 0.654 | 93.356 | 13.324 | 110.398 | 9.148 | 145.807 | -41.536 | 61.566 | 30.380 | 216.454 | 4.28 | 0.6388 |
| **Lateral reticular nucleus** | 140.962 | 294.438 | 22.907 | 151.843 | 9.078 | 157.205 | 46.128 | 95.565 | 44.557 | 162.882 | -42.173 | 57.649 | 5.841 | 0.4413 |
| **Lobus caudalis cerebeli** | 25.707 | 86.273 | 4.142 | 82.703 | 1.066 | 108.516 | -24.288 | 60.112 | 48.452 | 293.914 | -45.758 | 77.736 | 8.904 | 0.1791 |
| **Locus coeruleus** | 17.859 | 105.558 | 0.821 | 86.997 | -20.598 | 76.757 | -16.107 | 88.129 | -28.842 | 68.792 | -60.076 | 43.039 | 7.232 | 0.3000 |
| **Mauthner** | 7.266 | 70.902 | 3.258 | 87.619 | 17.344 | 76.833 | 41.112 | 176.068 | 12.558 | 190.172 | -37.113 | 74.044 | 5.149 | 0.5249 |
| **Medial vestibular nucleus** | 89.871 | 168.577 | 8.513 | 82.107 | 20.176 | 130.921 | 177.046 | 583.144 | 136.959 | 492.827 | -78.889 | 25.484 | 12.1 | 0.0597 |
| **Noradrenergic neurons of the interfascicular and vagal areas** | 98.684 | 215.212 | 31.522 | 179.366 | 4.531 | 87.899 | -9.475 | 96.441 | 208.736 | 717.745 | -57.682 | 55.332 | 6.792 | 0.3405 |
| **Raphe-inferior** | 30.991 | 119.854 | -23.144 | 63.171 | 14.393 | 96.993 | -29.271 | 60.029 | 50.419 | 213.944 | -59.368 | 38.006 | 4.394 | 0.6235 |
| **Raphe-superior** | 10.799 | 90.296 | 13.465 | 118.456 | 39.021 | 111.724 | 28.942 | 138.162 | -6.153 | 110.287 | 6.997 | 111.633 | 2.062 | 0.9139 |
| **Tangential vestibular nucleus** | 38.850 | 186.743 | -13.496 | 79.943 | -27.921 | 59.014 | -17.666 | 78.742 | -41.951 | 55.774 | -37.037 | 40.699 | 2.507 | 0.8677 |
| **Valvula cerebelli** | 29.665 | 88.541 | -4.485 | 87.170 | 19.801 | 117.958 | -11.360 | 93.183 | -40.850 | 91.134 | -59.498 | 31.783 | 9.846 | 0.1313 |

| | | | | | | | | | | | | | | |
|---|---|---|---|---|---|---|---|---|---|---|---|---|---|---|
| **Eyes** | 30.640 | 56.685 | 33.738 | 72.389 | -11.869 | 53.321 | 22.849 | 78.963 | -8.450 | 49.171 | -44.888 | 24.546 | 10.39 | 0.1091 |
| **Olfactory epithelium** | 22.804 | 56.378 | -5.917 | 59.491 | 102.171 | 129.448 | 1.097 | 60.244 | 86.422 | 144.723 | 5.351 | 77.333 | 8.654 | 0.1940 |
| **Vagal ganglia** | 219.738 | 563.361 | -9.230 | 50.341 | 5.882 | 34.577 | -8.453 | 44.259 | -10.865 | 69.670 | -19.934 | 56.387 | 4.673 | 0.5864 |
| **Spinal cord** | 72.492 | 156.693 | 53.983 | 243.863 | 45.668 | 156.561 | 86.021 | 294.595 | 214.364 | 756.521 | -58.564 | 48.806 | 7.512 | 0.2761 |
| **Neuropil region** | 95.421 | 163.172 | 26.543 | 167.520 | 101.894 | 257.875 | 53.450 | 229.212 | 334.086 | 1060.110 | -61.812 | 32.066 | 6.979 | 0.3228 |

#### Isoeugenol

Table S10. **Mean fluorescence intensity in brain regions of interest (ROIs) measured using light-sheet imaging during the first 8 seconds post application of a light stimulus, after a 3-minute exposure to isoeugenol.** Bold denotes ROIs where anaesthetic administration was found to have a statistically significant effect on activity.

| ROI | Control | | 0.2mM | | 0.4mM | | 1.6mM | | 2.4mM | | Kruskal-Wallis statistic | p-value |
|---|---|---|---|---|---|---|---|---|---|---|---|---|
| | Mean | SD | Mean | SD | Mean | SD | Mean | SD | Mean | SD | | |
| Telencephalon | 22.290 | 160.490 | 29.035 | 143.080 | 266.547 | 395.427 | 8.142 | 86.799 | -49.244 | 35.755 | 4.089 | 0.5366 |
| Olfactory Bulbs | 2.046 | 166.347 | -7.788 | 85.533 | 135.401 | 214.534 | 1.223 | 102.325 | -1.194 | 85.921 | 3.676 | 0.5969 |
| Anterior commissure | 405.516 | 1188.209 | 199.602 | 339.953 | 177.639 | 210.103 | 130.474 | 277.540 | 38.816 | 296.997 | 5.358 | 0.3737 |
| Pallium | 1.016 | 133.890 | 87.337 | 279.349 | 342.176 | 500.144 | 77.449 | 156.802 | -41.442 | 41.459 | 5.681 | 0.3385 |
| Subpallium | 19.845 | 102.026 | 291.646 | 649.468 | 8.577 | 86.911 | -23.006 | 75.241 | -36.581 | 54.740 | 2.504 | 0.7759 |
| Diencephalon | 23.092 | 206.174 | 76.429 | 186.866 | 140.794 | 241.978 | 145.556 | 277.436 | -22.106 | 73.384 | 3.352 | 0.6459 |
| Dorsal thalamus | -29.390 | 95.640 | 634.629 | 1654.149 | 21.057 | 150.226 | 173.471 | 317.765 | -50.648 | 36.786 | 4.328 | 0.5032 |
| Eminentia thalami | 293.245 | 898.681 | 53.765 | 183.424 | 323.437 | 350.699 | 177.721 | 245.288 | 305.247 | 866.968 | 10.66 | 0.0586 |
| Habenulae | 39.863 | 190.345 | 84.739 | 245.981 | 265.688 | 504.980 | 186.744 | 471.694 | -7.159 | 60.184 | 3.444 | 0.6319 |
| Intermediate hypothalamus | -35.520 | 83.949 | 87.221 | 254.232 | 78.664 | 244.695 | 90.210 | 207.794 | -18.012 | 114.188 | 2.588 | 0.7632 |
| Pineal | 64.609 | 273.227 | -16.306 | 74.476 | 204.803 | 337.245 | 76.366 | 139.679 | -39.697 | 40.452 | 6.782 | 0.2374 |
| Posterior tuberculum | -51.077 | 31.000 | 61.617 | 213.978 | 106.265 | 376.967 | 38.575 | 137.316 | -25.638 | 68.016 | 2.304 | 0.8057 |
| Preoptic area | 195.724 | 508.348 | 52.875 | 159.388 | 422.846 | 594.791 | 98.106 | 198.091 | 32.290 | 165.801 | 7.88 | 0.163 |
| Pretectum | -12.708 | 122.378 | 767.293 | 2008.793 | 45.234 | 162.533 | 68.698 | 153.904 | -47.017 | 44.979 | 2.702 | 0.7459 |
| Rostral hypothalamus | -37.304 | 61.635 | -21.933 | 78.416 | 10.197 | 128.236 | 49.812 | 192.163 | -45.037 | 37.259 | 2.558 | 0.7678 |
| Ventral thalamus | -17.689 | 151.133 | 321.262 | 850.780 | 152.565 | 315.795 | 198.003 | 393.056 | -56.309 | 36.897 | 5.902 | 0.3159 |
| Mesencephalon | 138.243 | 332.709 | 558.046 | 1236.321 | 70.786 | 198.465 | 45.444 | 95.387 | -32.945 | 66.219 | 3.241 | 0.6629 |
| Tectum striatum periventiculare | 97.427 | 268.763 | 327.613 | 640.088 | 68.504 | 193.063 | 40.742 | 112.706 | -35.161 | 58.633 | 4.108 | 0.5339 |

| | | | | | | | | | | | | |
|---|---|---|---|---|---|---|---|---|---|---|---|---|
| **Tectum neuropil** | 302.345 | 503.425 | 487.344 | 1059.932 | 12.696 | 120.466 | 17.365 | 85.824 | -44.969 | 48.310 | 5.478 | 0.3603 |
| **Tegmentum** | 165.096 | 379.667 | 401.684 | 784.779 | 67.815 | 206.733 | 25.179 | 95.265 | -44.104 | 45.801 | 1.511 | 0.9117 |
| **Torus longitudinaris** | -24.812 | 57.745 | 142.062 | 313.694 | 13.709 | 116.468 | 140.789 | 326.680 | -33.839 | 59.665 | 4.719 | 0.4511 |
| **Torus semicircularis** | -48.547 | 40.681 | 85.151 | 245.550 | 13.245 | 131.381 | 14.134 | 77.344 | -26.696 | 59.679 | 2.997 | 0.7004 |
| **Rhombencephalon** | -1.698 | 190.214 | 261.386 | 467.387 | 167.302 | 375.786 | 55.147 | 222.941 | -35.752 | 36.159 | 2.568 | 0.7663 |
| **Area postrema** | -13.640 | 60.450 | 235.995 | 279.616 | 73.189 | 133.352 | 14.876 | 106.156 | 16.623 | 121.613 | 5.865 | 0.3196 |
| **Cerebellum** | 67.730 | 231.640 | 335.664 | 608.038 | 18.890 | 172.308 | 37.958 | 144.493 | -53.527 | 38.751 | 2.76 | 0.7369 |
| **Corpus cerebelli** | 60.804 | 204.945 | 322.311 | 583.341 | 14.499 | 171.379 | 43.786 | 152.935 | -62.208 | 31.502 | 2.661 | 0.7521 |
| **Eminentia granularis** | 81.479 | 403.475 | 281.971 | 540.635 | 89.138 | 244.378 | 40.261 | 152.464 | -37.320 | 44.578 | 3.055 | 0.6915 |
| **Inferior olive** | -32.744 | 74.013 | 173.954 | 315.983 | 101.753 | 214.976 | 50.243 | 216.564 | -19.130 | 41.056 | 3.171 | 0.6737 |
| **Interpeduncular nucleus** | -34.407 | 86.905 | 49.981 | 178.399 | 96.745 | 440.009 | 31.487 | 82.493 | -10.310 | 82.130 | 4.298 | 0.5073 |
| **Lateral reticular nucleus** | -20.550 | 80.491 | 178.387 | 307.750 | 128.813 | 309.336 | 21.110 | 148.286 | 1.166 | 73.969 | 2.739 | 0.7402 |
| **Lobus caudalis cerebeli** | 73.731 | 247.595 | 406.150 | 808.864 | 75.510 | 207.478 | 66.645 | 247.559 | -48.703 | 55.265 | 2.651 | 0.7537 |
| **Locus coeruleus** | 270.876 | 806.587 | 247.631 | 415.344 | -24.413 | 82.993 | 40.496 | 86.288 | -52.793 | 41.953 | 5.556 | 0.3518 |
| **Mauthner** | 142.322 | 345.885 | 66.950 | 247.543 | 30.201 | 158.452 | 30.003 | 159.693 | -45.758 | 42.540 | 1.876 | 0.866 |
| **Medial vestibular nucleus** | 53.138 | 257.214 | 197.500 | 378.461 | 130.233 | 312.994 | 12.840 | 103.705 | 16.712 | 134.830 | 0.2653 | 0.9982 |
| **Noradrenergic neurons of the interfascicular and vagal areas** | -26.381 | 81.428 | 196.680 | 312.815 | 97.407 | 241.645 | 16.775 | 105.577 | 0.411 | 103.028 | 2.945 | 0.7084 |
| **Raphe-inferior** | -36.667 | 102.404 | 100.025 | 296.288 | 88.748 | 294.861 | 31.357 | 193.339 | -46.511 | 47.405 | 2.772 | 0.7451 |
| **Raphe-superior** | 264.390 | 615.312 | 180.294 | 340.529 | -9.948 | 138.987 | -4.733 | 78.481 | -60.532 | 26.800 | 4.313 | 0.5054 |
| **Tangential vestibular nucleus** | -43.087 | 62.297 | 177.633 | 545.937 | 126.255 | 397.339 | -20.116 | 46.788 | -33.974 | 38.225 | 1.606 | 0.9005 |
| **Valvula cerebelli** | 263.815 | 510.801 | 229.211 | 539.478 | 13.012 | 176.066 | 28.336 | 97.012 | -50.675 | 47.789 | 3.162 | 0.675 |

| | | | | | | | | | | | | |
|---|---|---|---|---|---|---|---|---|---|---|---|---|
| **Eyes** | -38.539 | 45.887 | 224.463 | 506.872 | 13.358 | 80.252 | -2.514 | 45.006 | -40.594 | 41.469 | 3.935 | 0.5588 |
| **Olfactory epithelium** | 12.073 | 121.565 | 105.245 | 336.778 | 104.422 | 120.447 | -7.940 | 69.657 | 6.613 | 94.127 | 6.504 | 0.2602 |
| **Vagal ganglia** | -38.735 | 22.920 | 31.200 | 110.883 | 65.947 | 180.846 | -0.989 | 70.915 | -14.536 | 72.360 | 7.131 | 0.2111 |
| **Spinal cord** | -38.192 | 62.378 | 142.598 | 368.244 | 72.060 | 182.275 | 8.006 | 86.271 | 32.774 | 126.488 | 3.788 | 0.5803 |
| **Neuropil region** | -41.520 | 58.446 | 163.807 | 384.562 | 69.375 | 184.363 | 29.253 | 88.965 | -6.023 | 97.914 | 4.742 | 0.5871 |

## MS222

Table S11. **Mean fluorescence intensity in brain regions of interest (ROIs) measured using light-sheet imaging during the first 8 seconds post application of a light stimulus, after a 3-minute exposure to MS222.** Bold denotes ROIs where anaesthetic administration was found to have a statistically significant effect on activity.

| ROI | Control | | 0.50mM | | 0.70mM | | 1.00mM | | Kruskal-Wallis statistic | p-value |
|---|---|---|---|---|---|---|---|---|---|---|
| | Mean | SD | Mean | SD | Mean | SD | Mean | SD | | |
| Telencephalon | 145.255 | 381.441 | -10.145 | 105.270 | 7.169 | 99.795 | -60.441 | 32.599 | 5.162 | 0.1603 |
| Olfactory Bulbs | 55.899 | 162.501 | -25.731 | 42.949 | 5.924 | 76.117 | 9.038 | 74.195 | 1.031 | 0.7936. |
| **Anterior commissure** | **283.872** | **522.424** | **-25.954** | **83.010** | **108.386** | **222.904** | **-72.965** | **17.387** | **9.057** | **0.0285** |
| Pallium | 67.121 | 202.609 | -50.693 | 39.031 | -8.703 | 96.146 | -60.826 | 37.751 | 6.582 | 0.0865 |
| Subpallium | 199.298 | 465.942 | 56.033 | 233.696 | 19.182 | 94.790 | -62.218 | 40.721 | 6.276 | 0.0989 |
| **Diencephalon** | **80.168** | **130.965** | **-19.684** | **146.143** | **10.096** | **163.609** | **-71.609** | **26.478** | **9.168** | **0.0271** |
| **Dorsal thalamus** | **67.059** | **136.533** | **3.561** | **202.209** | **-38.951** | **87.637** | **-61.776** | **43.396** | **8.315** | **0.0399** |
| **Eminentia thalami** | **8.154** | **72.218** | **-43.441** | **55.809** | **-53.716** | **43.222** | **-74.639** | **20.577** | **8.599** | **0.0351** |
| Habenulae | 61.636 | 104.146 | -26.747 | 79.829 | -10.797 | 126.190 | -46.740 | 40.299 | 7.321 | 0.0623 |
| **Intermediate hypothalamus** | **114.099** | **228.537** | **-2.166** | **206.426** | **-48.052** | **78.723** | **-71.079** | **28.815** | **8.861** | **0.0312** |
| Pineal | 253.127 | 546.690 | -37.268 | 42.412 | 0.710 | 79.593 | -39.907 | 28.199 | 6.946 | 0.0736 |
| **Posterior tuberculum** | **89.171** | **187.930** | **7.396** | **207.243** | **-16.628** | **144.866** | **-63.501** | **37.653** | **7.946** | **0.0471** |
| **Preoptic area** | **85.810** | **107.480** | **-57.767** | **37.240** | **24.006** | **242.058** | **-69.945** | **33.699** | **10.87** | **0.0125** |
| Pretectum | 88.912 | 162.286 | 19.409 | 241.727 | -45.000 | 63.326 | -51.034 | 64.735 | 6.997 | 0.072 |
| **Rostral hypothalamus** | **73.725** | **159.246** | **-27.772** | **85.719** | **-51.738** | **51.384** | **-65.282** | **33.178** | **9.976** | **0.0188** |
| Ventral thalamus | 86.198 | 207.877 | -14.082 | 168.340 | -26.561 | 107.673 | -56.185 | 60.615 | 6.344 | 0.096 |
| **Mesencephalon** | **102.913** | **181.764** | **-3.665** | **210.728** | **-35.377** | **96.271** | **-78.072** | **24.223** | **10.74** | **0.0132** |
| **Tectum striatum periventiculare** | **96.304** | **165.606** | **47.696** | **349.803** | **-45.416** | **85.582** | **-76.589** | **24.847** | **11.11** | **0.0112** |

| | | | | | | | | | | |
|---|---|---|---|---|---|---|---|---|---|---|
| **Tectum neuropil** | **89.985** | **175.181** | **-33.559** | **103.856** | **-23.215** | **94.870** | **-75.360** | **15.738** | **9.594** | **0.0224** |
| **Tegmentum** | **110.189** | **162.627** | **196.395** | **791.787** | **-46.010** | **87.663** | **-75.230** | **38.357** | **10.44** | **0.0152** |
| **Torus longitudinaris** | 40.626 | 107.127 | -19.933 | 48.223 | -31.611 | 51.384 | -44.445 | 47.652 | 4.116 | 0.2492 |
| **Torus semicircularis** | **88.242** | **204.190** | **-60.184** | **36.796** | **-37.872** | **78.781** | **-82.473** | **9.063** | **8.901** | **0.0306** |
| **Rhombencephalon** | **125.453** | **273.923** | **0.738** | **224.945** | **-70.340** | **52.714** | **-76.125** | **32.997** | **10.61** | **0.0141** |
| **Area postrema** | **679.554** | **1662.135** | **100.291** | **350.693** | **-67.487** | **60.287** | **-66.517** | **26.358** | **8.98** | **0.0296** |
| **Cerebellum** | 81.717 | 172.061 | 99.866 | 485.499 | -65.104 | 64.843 | -44.414 | 90.481 | 7.665 | 0.0535 |
| **Corpus cerebelli** | **81.022** | **179.540** | **157.167** | **648.909** | **-67.933** | **57.040** | **-30.784** | **124.252** | **8.81** | **0.0319** |
| **Eminentia granularis** | 35.587 | 124.907 | 69.097 | 221.239 | -17.552 | 139.273 | -59.250 | 38.129 | 3.116 | 0.374 |
| **Inferior olive** | **154.592** | **299.647** | **-29.011** | **93.495** | **-62.084** | **59.396** | **-57.629** | **59.922** | **8.361** | **0.0391** |
| **Interpeduncular nucleus** | **128.650** | **272.381** | **-18.644** | **182.338** | **-60.821** | **70.054** | **-51.721** | **76.964** | **9.764** | **0.0207** |
| **Lateral reticular nucleus** | 73.625 | 186.528 | -28.126 | 44.861 | -55.049 | 55.876 | -53.015 | 23.226 | 4.526 | 0.21 |
| **Lobus caudalis cerebelli** | **24.206** | **99.264** | **-15.016** | **144.659** | **-55.822** | **73.771** | **-57.762** | **61.597** | **8.193** | **0.0422** |
| **Locus coeruleus** | **23.765** | **104.256** | **-12.047** | **179.705** | **-71.052** | **41.306** | **-82.327** | **16.583** | **11.18** | **0.0108** |
| **Mauthner** | **29.251** | **157.526** | **41.070** | **346.363** | **-71.625** | **38.301** | **-85.589** | **11.195** | **9.506** | **0.0233** |
| **Medial vestibular nucleus** | **137.104** | **337.639** | **7.095** | **194.179** | **-71.544** | **44.003** | **-92.108** | **2.558** | **11.76** | **0.0082** |
| **Noradrenergic neurons of the interfascicular and vagal areas** | 316.109 | 526.837 | 94.766 | 445.769 | -49.594 | 111.207 | -72.498 | 23.675 | 7.17 | 0.0667 |
| **Raphe-inferior** | **126.546** | **279.200** | **8.811** | **201.382** | **-68.078** | **37.787** | **-79.506** | **17.676** | **9.142** | **0.0275** |
| **Raphe-superior** | **160.614** | **221.065** | **27.209** | **288.977** | **-76.582** | **26.928** | **-35.799** | **74.861** | **10.19** | **0.017** |
| **Tangential vestibular nucleus** | **76.597** | **146.204** | **-55.505** | **53.017** | **-59.630** | **59.574** | **-78.704** | **16.248** | **10.85** | **0.0126** |
| **Valvula cerebelli** | 139.396 | 203.513 | 107.223 | 455.972 | -60.625 | 54.412 | 0.816 | 161.546 | 7.693 | 0.0605 |

| | | | | | | | | | | |
|---|---|---|---|---|---|---|---|---|---|---|
| **Eyes** | 83.962 | 181.841 | 14.238 | 79.552 | -4.002 | 109.056 | -36.456 | 36.500 | 6.264 | 0.0994 |
| **Olfactory epithelium** | -10.371 | 81.234 | -40.159 | 36.065 | 92.112 | 224.373 | 7.747 | 88.717 | 1.437 | 0.6968 |
| **Vagal ganglia** | 49.305 | 139.882 | -1.787 | 85.039 | -23.341 | 62.410 | -24.119 | 86.485 | 1.764 | 0.6228 |
| **Spinal cord** | 133.936 | 275.388 | -44.009 | 107.652 | -35.945 | 140.126 | -55.474 | 55.420 | 7.389 | 0.0605 |
| **Neuropil region** | 317.587 | 737.553 | -18.117 | 132.625 | -47.058 | 101.701 | -72.210 | 27.763 | 6.401 | 0.0937 |

#### Quinaldine sulfate

Table S12. **Mean fluorescence intensity in brain regions of interest (ROIs) measured using light-sheet imaging during the first 8 seconds post application of a light stimulus, after a 3-minute exposure to quinaldine sulfate.** Bold denotes ROIs where anaesthetic administration was found to have a statistically significant effect on activity.

| ROI | Control | | 0.08mM | | 0.20mM | | 0.80mM | | Kruskal-Wallis statistic | p-value |
|---|---|---|---|---|---|---|---|---|---|---|
| | Mean | SD | Mean | SD | Mean | SD | Mean | SD | | |
| Telencephalon | 342.238 | 863.057 | 96.970 | 303.947 | -51.631 | 36.868 | -14.333 | 122.775 | 6.219 | 0.1014 |
| Olfactory Bulbs | 108.500 | 380.199 | 107.200 | 207.884 | 6.055 | 133.314 | 24.256 | 73.429 | 1.81 | 0.6128 |
| Anterior commissure | 552.623 | 1260.245 | 272.240 | 494.020 | -51.463 | 29.337 | 191.950 | 509.841 | 6.298 | 0.098 |
| Pallium | 467.653 | 1090.251 | 72.766 | 308.315 | -50.670 | 44.987 | -0.606 | 166.958 | 7.548 | 0.0563 |
| Subpallium | 82.587 | 202.805 | 32.433 | 115.919 | -55.140 | 28.118 | 2.315 | 64.897 | 5.389 | 0.1454 |
| **Diencephalon** | **74.709** | **187.017** | **22.861** | **99.712** | **-68.165** | **35.358** | **-20.340** | **188.047** | **10.97** | **0.0119** |
| **Dorsal thalamus** | **55.675** | **131.133** | **3.951** | **85.314** | **-63.567** | **39.934** | **-26.118** | **171.132** | **9.923** | **0.0192** |
| **Eminentia thalami** | **310.911** | **605.709** | **39.745** | **147.615** | **-62.411** | **52.940** | **-67.362** | **43.487** | **9.733** | **0.021** |
| Habenulae | 289.273 | 558.011 | 147.192 | 430.143 | -39.834 | 92.485 | -53.259 | 51.389 | 6.151 | 0.1045 |
| Intermediate hypothalamus | 52.586 | 125.901 | 61.917 | 168.860 | 41.098 | 265.169 | -37.329 | 123.214 | 7.528 | 0.0568 |
| Pineal | 64.840 | 164.271 | 7.570 | 203.059 | -45.851 | 37.913 | 18.787 | 78.568 | 6.216 | 0.1016 |
| **Posterior tuberculum** | **62.183** | **140.914** | **30.889** | **102.595** | **-48.590** | **78.810** | **-63.302** | **69.217** | **12.29** | **0.0065** |
| **Preoptic area** | **206.602** | **361.337** | **46.793** | **97.765** | **-65.320** | **32.726** | **-72.393** | **30.501** | **10.38** | **0.0156** |
| **Pretectum** | **19.120** | **94.590** | **0.406** | **108.893** | **-59.837** | **47.947** | **-50.973** | **96.738** | **9.276** | **0.0258** |
| **Rostral hypothalamus** | **113.884** | **234.238** | **91.563** | **288.660** | **-34.267** | **106.900** | **-75.188** | **20.805** | **9.838** | **0.02** |
| **Ventral thalamus** | **95.916** | **180.997** | **7.141** | **102.332** | **-59.303** | **52.728** | **-66.200** | **60.379** | **9.991** | **0.0186** |
| **Mesencephalon** | **36.539** | **124.350** | **136.300** | **341.069** | **-49.969** | **74.894** | **0.104** | **214.454** | **8.043** | **0.0451** |
| Tectum striatum periventiculare | 29.546 | 107.584 | 98.631 | 234.851 | -46.518 | 78.054 | -18.521 | 149.852 | 7.017 | 0.0714 |

| | | | | | | | | | | |
|---|---|---|---|---|---|---|---|---|---|---|
| **Tectum neuropil** | 37.365 | 120.535 | 66.111 | 174.370 | -54.264 | 44.511 | -5.938 | 155.039 | 6.026 | 0.1104 |
| **Tegmentum** | **14.307** | **98.943** | **70.409** | **172.905** | **-36.199** | **126.363** | **-35.983** | **137.271** | **68.241** | **0.0413** |
| **Torus longitudinaris** | **66.463** | **197.975** | **-24.050** | **112.153** | **-70.857** | **12.877** | **16.181** | **107.318** | **7.815** | **0.05** |
| **Torus semicircularis** | **99.378** | **242.960** | **114.194** | **246.002** | **-70.220** | **20.790** | **125.249** | **457.982** | **9.318** | **0.0253** |
| **Rhombencephalon** | 5.546 | 111.022 | 70.707 | 235.000 | -39.498 | 112.467 | -31.542 | 152.798 | 5.642 | 0.1304 |
| **Area postrema** | **29.109** | **157.891** | **103.509** | **130.615** | **0.982** | **230.943** | **-43.177** | **59.304** | **9.997** | **0.0186** |
| **Cerebellum** | **15.433** | **109.288** | **9.048** | **107.249** | **-29.956** | **146.634** | **-61.486** | **66.103** | **9.832** | **0.02** |
| **Corpus cerebelli** | **6.444** | **95.799** | **13.096** | **109.270** | **-21.605** | **172.773** | **-63.678** | **56.107** | **9.236** | **0.0263** |
| **Eminentia granularis** | **318.819** | **689.275** | **672.061** | **1846.144** | **-53.258** | **55.598** | **55.842** | **392.694** | **8.753** | **0.0328** |
| **Inferior olive** | 66.153 | 219.214 | 229.268 | 547.172 | -8.051 | 191.902 | -39.618 | 111.926 | 4.56 | 0.207 |
| **Interpeduncular nucleus** | -6.480 | 84.659 | 70.642 | 169.665 | -27.261 | 98.024 | -73.698 | 31.415 | 7.616 | 0.0546 |
| **Lateral reticular nucleus** | 52.151 | 203.461 | 83.133 | 170.808 | -27.751 | 92.759 | 98.510 | 360.417 | 2.608 | 0.4561 |
| **Lobus caudalis cerebelli** | 3.231 | 127.136 | 37.261 | 126.939 | -26.091 | 147.982 | -36.852 | 108.948 | 4.27 | 0.2338 |
| **Locus coeruleus** | **3.966** | **81.399** | **64.680** | **154.942** | **-9.888** | **184.112** | **-66.099** | **50.270** | **9.21** | **0.0266** |
| **Mauthner** | **38.586** | **151.940** | **55.042** | **214.476** | **33.528** | **317.994** | **-45.023** | **111.018** | **8.534** | **0.0362** |
| **Medial vestibular nucleus** | 24.221 | 181.448 | 86.423 | 300.183 | -7.475 | 215.987 | -64.030 | 62.570 | 7.145 | 0.0674 |
| **Noradrenergic neurons of the interfascicular and vagal areas** | 51.871 | 184.343 | 385.054 | 806.785 | -36.135 | 93.322 | 61.281 | 251.259 | 6.784 | 0.0791 |
| **Raphe-inferior** | 16.188 | 159.115 | 94.089 | 312.664 | -10.508 | 186.094 | 3.977 | 224.891 | 3.145 | 0.3698 |
| **Raphe-superior** | 1.029 | 96.597 | 66.103 | 172.568 | -58.399 | 43.478 | -57.060 | 36.657 | 4.884 | 0.1805 |
| **Tangential vestibular nucleus** | 56.154 | 153.044 | 16.480 | 106.699 | -53.452 | 68.113 | -30.981 | 112.135 | 7.366 | 0.0611 |
| **Valvula cerebelli** | **-1.616** | **102.413** | **35.836** | **96.202** | **-51.306** | **66.450** | **-70.396** | **36.604** | **9.111** | **0.0279** |

| | | | | | | | | | | |
|---|---|---|---|---|---|---|---|---|---|---|
| **Eyes** | 94.182 | 172.912 | -6.736 | 33.332 | -28.837 | 63.851 | 59.974 | 108.540 | 7.474 | 0.0582 |
| **Olfactory epithelium** | 70.458 | 224.920 | -17.351 | 48.569 | -51.888 | 33.949 | 8.254 | 66.689 | 3.963 | 0.2655 |
| **Vagal ganglia** | 71.924 | 191.415 | 104.633 | 132.076 | 17.314 | 184.286 | 123.381 | 259.434 | 4.23 | 0.2377 |
| **Spinal cord** | **46.679** | **128.172** | **425.739** | **801.345** | **-29.578** | **145.757** | **-41.491** | **60.788** | **8.94** | **0.0301** |
| **Neuropil region** | 56.727 | 231.159 | 361.007 | 610.467 | -29.868 | 137.604 | -24.820 | 107.061 | 7.517 | 0.0571 |

#### Behavioural measures of tolerability – locomotion

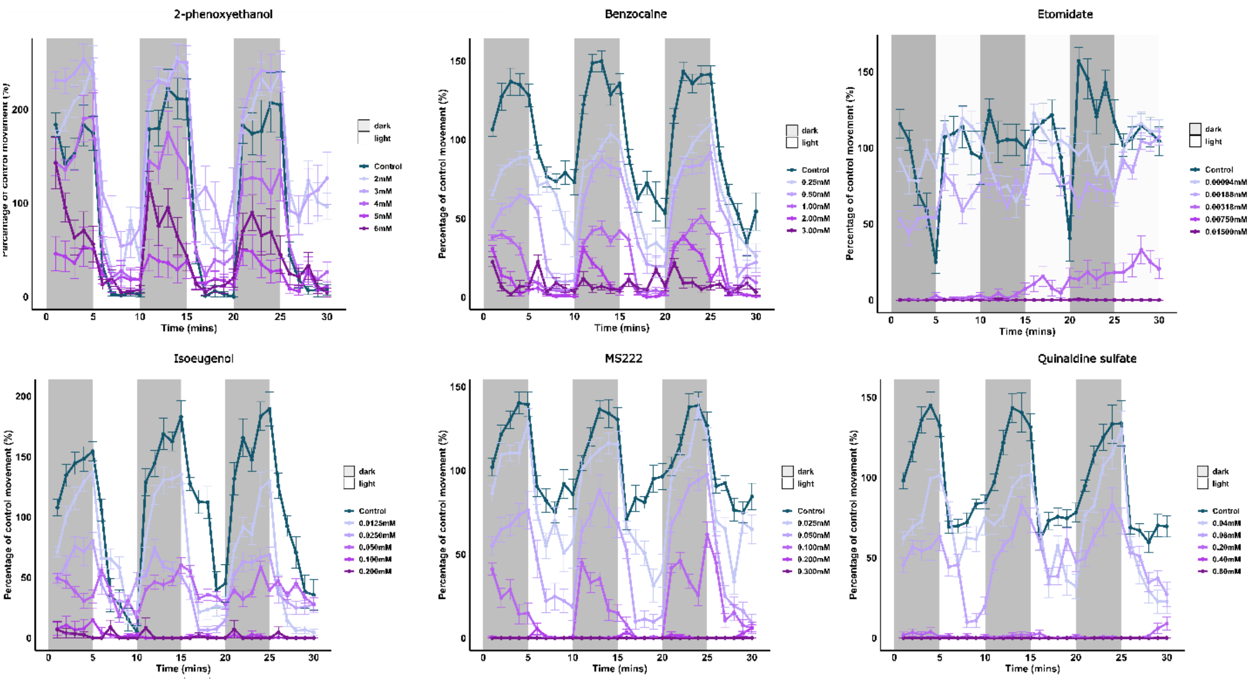


**Figure S2. Effects of different anaesthetic compounds on embryo-larval movement, normalised to a 10-minute acclimation period.** Grey shaded boxes denote dark phase periods and white areas denote light phase periods. Data are shown as means for the 30-minute experimental duration, +/- SEM (n=16). Data were normalised against the overall movement of the control group during the acclimation period (first 10 minutes of the assay), in order to reduce the variability between different experimental runs.

The data from the general locomotor and light dark assays are summarised in Figure S2. For all compounds analysed with the exception of isoeugenol (for dark only), there was a statistically significant interaction between time and treatment (2-phenoxyethanol, light: $\chi2$= 62.259, p< 0.001, dark: $\chi2$= 25.4709, p=0.005; benzocaine, light: $\chi2$=34.666, p< 0.001, dark: $\chi2$= 62.224, p<0.001; etomidate, light: $\chi2$= 131.783, p< 0.001, dark: $\chi2$= 371.070, p<0.001; isoeugenol, light: $\chi2$= 120.909, p< 0.001; MS222, light: $\chi2$= 34.025, p< 0.001, dark: $\chi2$= 122.665 , p<0.001; quinaldine sulfate, light: $\chi2$= 132.163, p< 0.001; quinaldine sulfate, dark: $\chi2$= 16.516, p=0.009)). As expected, the higher concentrations of anaesthetics tested across the selected range generally led to decreased locomotion, with the exception of 2-phenoxyethanol (2mM and 3mM, in the light phase), where, in some cases, elevated locomotion was exhibited compared with control groups. For isoeugenol, in the dark phase, we observed a significant effect of treatment ($\chi2$= 234.3253, p< 0.0001), but not time.

#### Behavioural measures of tolerability – olfactory-mediated avoidance behaviour

**Table S13. The effect of the anaesthetic compounds tested on total shift of each group from the central point of the arena along the x axis measured during an assay of olfactory-mediated avoidance behaviour**. Data analysed by a generalised least squares estimation model (GLS) to account for differences in heteroscedasticity between the different treatments. Bold denotes statistically significant variables. † denotes compounds dissolved in ethanol instead of water.

| Coefficients | | Value | Std. error | t-value | p-value |
|---|---|---|---|---|---|
| **Intercept** | | **6.424630** | **0.4548336** | **14.125232** | **<0.0001** |
| Average distance travelled | | -0.002794 | 0.0015085 | -1.852191 | 0.0684 |
| **Side of administration** | Left | | | | |
| | **Right** | **-0.590551** | **0.1743223** | **-3.387699** | **0.0012** |
| **Treatment** | **Water control** | | | | |
| | **Cadaverine (20mM)** | **1.066671** | **0.3622344** | **2.944700** | **0.0044** |
| | 2-phenoxyethanol (2mM) † | 0.776132 | 0.4718979 | 1.644704 | 0.1047 |
| | Benzocaine (0.2mM) † | -0.435796 | 0.5235524 | -0.832382 | 0.4082 |
| | Etomidate (0.016mM) † | 0.252526 | 0.6004404 | 0.420568 | 0.6754 |
| | Isoeugenol (0.1mM) † | 0.281224 | 0.3517207 | 0.799568 | 0.4268 |
| | MS222 (0.1mM) | -0.194175 | 0.5571213 | -0.348533 | 0.7285 |
| | **Quinaldine sulfate (0.1mM in Water)** | **1.587505** | **0.4088773** | **3.882595** | **0.0002** |

#### Cardiovascular physiology

##### Atrial and ventricular beat rates

Both atrial (ABR) and ventricular (VBR) beat rate decreased under treatment concentrations for 2-phenoxyethanol (ABR: $\chi^2_{(4)}$ = 24.31, p<0.0001 and VBR: $\chi^2_{(4)}$=23.06, p<0.0001), benzocaine (ABR: $\chi^2_{(4)}$=23.8, p<0.0001 and VBR: $\chi^2_{(4)}$=23.64, p<0.0001), etomidate (ABR: $\chi^2_{(4)}$=29.96, p<0.001 and VBR: $\chi^2_{(4)}$=32.76, p<0.001), isoeugenol (ABR: $\chi^2_{(4)}$=23.95, p<0.0001 and VBR: $\chi^2_{(4)}$=23.71, p<0.0001), MS222 (ABR: $F_{(2)}$=14.31, p=0.0001 and VBR: $F_{(2)}$=15.33, p<0.0001) and quinaldine sulfate (ABR: $F_{(3)}$=80.88, p<0.0001and VBR: $F_{(3)}$=21.69, p<0.0001). These differences were observed under the middle and highest test concentrations for benzocaine (ABR and VBR: 1mM: p=0.03, 1.6mM: p<0.0001), etomidate (ABR: 0.128 mM: p= 0.0045, 0.256 mM: p<0.001, and VBR: 0.128 mM: p= 0.0013, 0.256 mM: p<0.001), isoeugenol (ABR: 0.4mM p=0.005, 0.8mM <0.0001 and VBR: 0.4mM: p=0.004, 0.8mM: p<0.0001) and quinaldine sulfate (ABR 1.6mM: p<0.0001, 3.2mM: p<0.0001; VBR 1.6mM: p=0.0001, 3.2mM: p<0.0001). For 2-phenoxyethanol and MS222 these differences were only seen between the highest treatment and control (2-phenoxyethanol: ABR: p =0.001, VBR: p=0.0002 and MS222: ABR: p<0.0001, VBR: p<0.0001).

**Blood flow and blood linear velocity**

Blood flow was decreased for all compounds except for etomidate (2-phenoxyethanol: $\chi^2_{(4)}$=23.55, p<0.0001, benzocaine: $F_{(3)}$=12.97, p<0.0001, isoeugenol: $\chi^2_{(4)}$=19.71, p=0.0002, MS222: $F_{(2)}$=7.03, p=0.0046 and quinaldine sulfate: $F_{(3)}$=13.85, p<0.0001). Decreased blood flow as seen under all treatments in comparison to the control for quinaldine sulfate (3.2mM: p<0.0001, 1.6mM: p<0.0001 and 0.8mM p=0.0093). Blood flow was decreased at both the middle and highest concentrations tested for both 2-phenoxyethanol (4mM: p=0.03 and 7mM: p<0.0001) and benzocaine (1mM: p=0.008, 1.6mM: p<0.0001). For MS222 decreased blood flow was observed under the low (0.5mM: p=0.04) and middle (1mM: p=0.0026) concentrations.

There was an overall effect of treatment on the linear velocity of blood flow for all compounds (2-phenoxyethanol: $F_{(3)}$=29.9, p<0.0001, benzocaine: $F_{(3)}$=32.35, p<0.0001, etomidate: $\chi^2_{(4)}$=11.35, p=0.01, isoeugenol: $\chi^2_{(4)}$=21.63, p<0.0001, MS222: $F_{(2)}$=11.73, p=0.0004 and quinaldine sulphate: $\chi^2_{(2)}$=12.5, p=0.002 $F_{(3)}$=20.24, p<0.0001). Blood flow linear velocity was reduced under all treatments in comparison to the control for both benzocaine (0.6mM: p<0.038, 1mM: p=0.0006 and 1.6mM p<0.0001) and quinaldine sulfate (0.8mM: p=0.0004, 1.6mM: p<0.0001 and 3.2mM: p<0.0001). A reduction in the linear velocity of blood flow was observed under the middle and high concentration for 2-phenoxyethanol (4mM: p=0.0015, 7mM: p<0.0001) and isoeugenol (0.4mM: p=0.025, 0.8mM: p<0.0001). For MS222, the low and middle concentrations both decreased blood linear velocity (0.5mM p=0.0463, 1mM: p=0.0002). We were not able to detect any statistically significant differences between the different etomidate concentrations: while this compound has an overall depressive effect on linear blood flow, there were no significant differences between the concentrations tested.

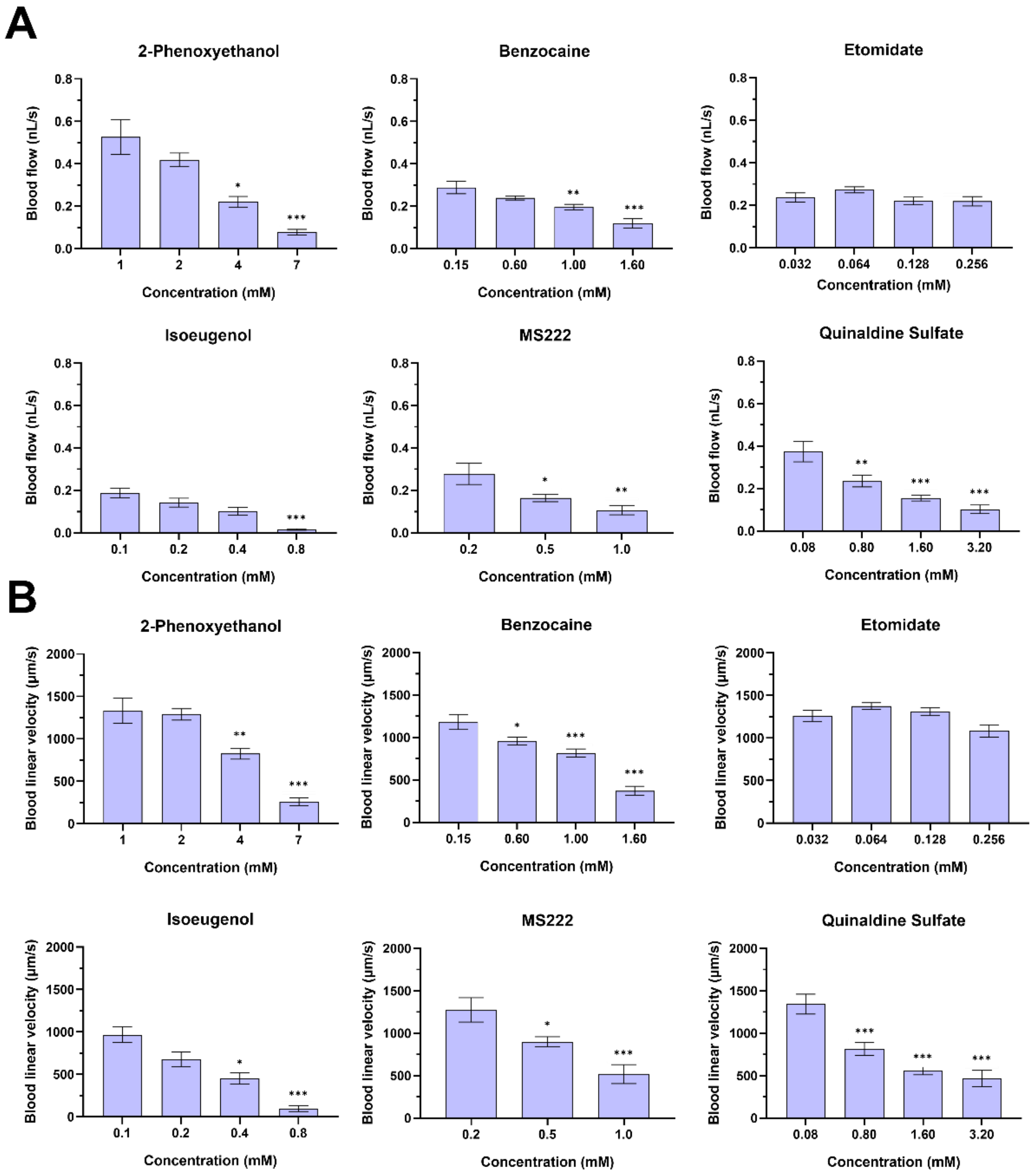


**Figure S3. Effects of the six test anaesthetics on blood flow (A) and blood linear velocity (B) in 4dpf zebrafish.** Data are presented as mean ± SEM across all fish tested (n=8). Statistically significant differences compared to control (lowest concentration for each compound) are presented as *p< 0.05, **p<0.01 and ***p<0.001.

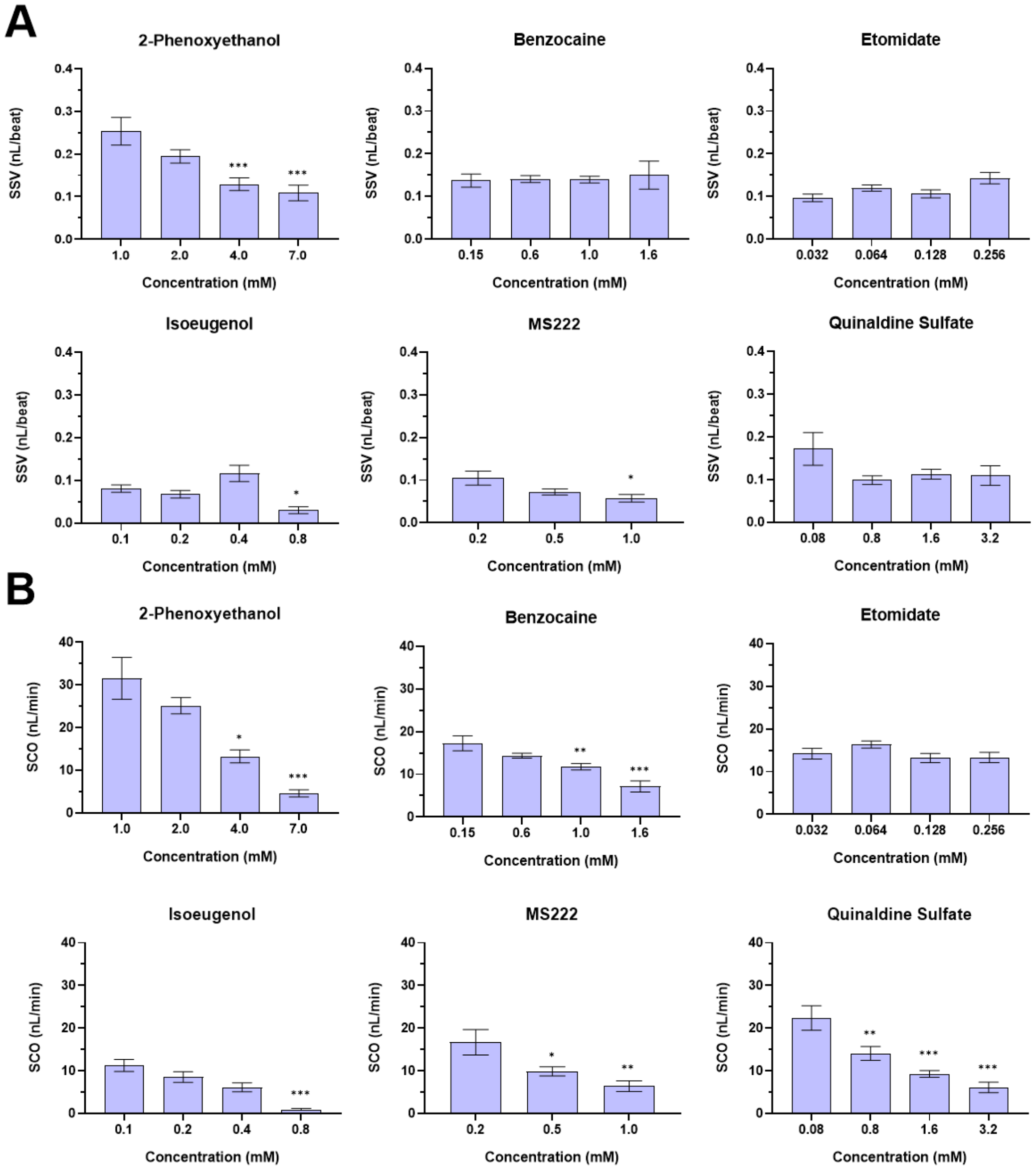


**Figure S4. Effects of the six test anaesthetics on surrogate stroke volume (SSV) (A) and surrogate cardiac output (SCO) (B) in 4dpf zebrafish.** Data are presented as mean ± SEM across all fish tested (n=8). Statistically significant differences compared to control (lowest concentration for each compound) are presented as *p< 0.05, **p<0.01 and ***p<0.001.

#### Surrogate stroke volume (SSV)

A decrease in SSV was observed for 2-phenoxyethanol ($F_{(3)}$=9.26, p=0.0002), isoeugenol ($\chi^2_{(4)}$=16.02, p=0.0011), and MS222 ($F_{(2)}$=4.26, p=0.028). For 2-phenoxyethanol this was observed under the middle (4mM: p=0.001) and highest treatment (7mM: p=0.0002) concentrations. For isoeugenol and MS222 a decrease was only observed under the highest concentration tested (isoeugenol 0.8mM: p=0.021 and MS222 1mM: p=0.018). In contrast, no effect of treatment concentration on SSV was observed for benzocaine, etomidate, or quinaldine sulfate.

**Surrogate cardiac output (SCO)**

There was evidence of an effect of concentration on SCO for all compounds except for etomidate (2-phenoxyethanol: $F_{(3)}=23.55$, $p<0.0001$, benzocaine: $F_{(3)}=12.97$, $p<0.0001$, isoeugenol: $\chi^2_{(4)}=19.71$, $p=0.0002$, MS222: $F_{(2)}=7.03$, $p=0.0046$ and quinaldine sulfate: $F_{(3)}=13.85$, $p<0.0001$). A decrease in SCO was observed for treatment concentrations in comparison to the control for quinaldine sulfate (0.8mM: $p=0.0093$ 1.6mM: $p<0.0001$ and 3.2mM: $p<0.0001$). SCO decreased under the mid and high concentrations for both 2-phenoxyethanol (4mM: $p=0.0032$, 7mM: $p<0.0001$) and benzocaine (1mM: $p=0.008$ 1.6mM: $p<0.0001$). For MS222, SCO was also decreased under the 0.5mM ($p=0.04$) and 1mM ($p=0.003$) treatments.

**Table S14. The effect of the anaesthetic compounds tested on cardiac function (values presented as means ± standard error.** Bold denotes statistical significance (p<0.05).

| Compound | Concentration (mM) | Atrial Beat Rate (ABR) | Ventricular Beat Rate (VBR) | Blood Flow (nL/s) | Linear Velocity (µm/s) | Surrogate Stroke Volume (SSV) (nL/beat) | Surrogate Cardiac Output (SCO) (nL/min) |
|---|---|---|---|---|---|---|---|
| 2-phenoxyethanol | 1 | 122.605±5.528 | 123.064±8.378 | 0.526±0.082 | 1331.474±150.430 | 0.253±0.032 | 31.565±4.929 |
| | 2 | 134.392±2.893 | 130.172±3.391 | 0.419±0.032 | 1286.790±66.558 | 0.194±0.016 | 25.146±1.902 |
| | 4 | 105.033±3.421 | 102.961±3.540 | **0.221±0.025** | **825.250±63.902** | **0.129±0.015** | **13.255±1.520** |
| | 7 | **44.036±3.542** | **44.073±3.328** | **0.077±0.014** | **260.266±47.458** | **0.109±0.019** | **4.620±0.827** |
| Benzocaine | 0.15 | 130.362±8.876 | 130.376±8.867 | 0.288±0.029 | 1181.608±87.385 | 0.137±0.015 | 17.296±1.749 |
| | 0.6 | 103.592±3.215 | 102.966±3.690 | 0.239±0.010 | **959.588±44.131** | 0.141±0.008 | 14.364±0.592 |
| | 1 | **84.925±3.885** | **84.820±3.846** | **0.197±0.012** | **819.028±46.066** | 0.140±0.008 | **11.806±0.747** |
| | 1.6 | **52.486±4.983** | **52.499±4.987** | **0.119±0.022** | **372.877±52.367** | 0.1499±0.033 | **7.169±1.309** |
| Etomidate | 0.032 | 147.160±6.399 | 147.178±6.374 | 0.237±0.070 | 1257.752±218.751 | 0.097±0.030 | 14.2231±4.218 |
| | 0.064 | 137.415±7.145 | 137.455±7.145 | 0.273±0.045 | 1375.393±122.535 | 0.120±0.023 | 16.372±2.708 |
| | 0.128 | **125.075±5.070** | **125.078±5.068** | 0.220±0.006 | 1309.357±139.059 | 0.106±0.029 | 13.201±3.429 |
| | 0.256 | **93.551±7.217** | **93.880±7.031** | 0.219±0.069 | 1082.771±227.455 | 0.143±0.040 | 13.340±3.610 |
| Isoeugenol | 0.1 | 137.551±5.060 | 137.600±5.051 | 0.188±0.024 | 967.554±91.234 | 0.081±0.009 | 11.261±1.411 |
| | 0.2 | 124.616±4.172 | 124.753±4.198 | 0.143±0.021 | 676.724±83.751 | 0.068±0.009 | 8.581±1.251 |
| | 0.4 | **58.706±11.897** | **61.084±11.529** | 0.102±0.017 | **450.797±66.421** | 0.117±0.019 | 6.119±1.048 |
| | 0.8 | **34.797±4.287** | **37.353±3.972** | **0.016±0.004** | **94.847±32.046** | **0.030±0.008** | **0.965±0.211** |
| MS222 | 0.2 | 155.077±6.102 | 155.128±6.115 | 0.278±0.050 | 1275.864±143.320 | 0.105±0.017 | 16.690±2.980 |
| | 0.5 | 136.626±4.726 | **136.610±4.714** | **0.165±0.018** | **899.663±59.922** | 0.072±0.007 | **9.885±1.069** |
| | 1 | **105.581±8.464** | **106.319±7.696** | **0.107±0.021** | **518.901±111.797** | **0.057±0.009** | **6.424±1.274** |
| | 5 | **5.375±2.783** | **5.375±2.783** | **0** | **0** | **0** | **0** |
| Quinaldine sulfate | 0.08 | 164.880±5.416 | 147.399±12.729 | 0.373±0.048 | 1343.397±117.250 | 0.172±0.038 | 22.403±2.877 |
| | 0.8 | 143.267±4.547 | 141.357±5.263 | **0.235±0.027** | **814.007±76.092** | 0.099±0.010 | **14.090±1.619** |
| | 1.6 | **83.281±5.943** | **87.3545±9.251** | **0.155±0.013** | **558.324±47.797** | 0.113±0.011 | **9.295±0.798** |
| | 3.2 | **56.392±6.079** | **58.306±2.544** | **0.102±0.020** | **466.891±95.063** | 0.109±0.023 | **6.1363±1.222** |

### Supplementary methods

##### Experimental animals

Adult zebrafish were held in aquaria at the University of Exeter, at 28°C ± 1°C under optimal conditions for spawning (14h:10h light:dark cycle, with 20min dusk-dawn transition periods). Culture water consisted of mains tap water filtered by reverse osmosis (Environmental Water Systems UK Ltd.) and then reconstituted with Analar-grade mineral salts to a standard synthetic freshwater composition (final ion concentrations: 117 mg·L−1 CaCl2.2H20, 25.0 mg·L−1 NaHCO3, 50 mg·L−1 MgSO4.7H20, 2.3 mg·L−1 KCl, 1.25 mg·L−1 Tropic Marine Sea Salt, giving a conductivity of 300 mS). The culture water was aerated and heated to 28°± 1°C before being supplied to individual zebrafish tanks on a recirculatory system. Water was routinely monitored for temperature, pH, conductivity, ammonia, nitrite and nitrate, all of which were maintained within appropriate limits for zebrafish. Embryos were collected shortly after spawning (at lights on) and cultured in Petri Dishes under the same conditions as the adults, and in density not exceeding 1 embryo/ml, until use. Embryo water was changed (50:50 volume) with fresh culture water once per day.

##### Determination of maximum tolerated and effective anaesthetic concentrations

Initial anaesthetic test concentrations (Table 1) were established from pilot testing. The stock test compound was freshly prepared in culture water (or appropriate solvent as required), and the pH checked and adjusted as necessary (to between 6.8-7.5) using 1 M NaOH/ 1 M HCl.

Four and a half dpf zebrafish larvae were placed in a 24-well seizure plate with one larva per well in 600 µl of culture water. For each anaesthetic, four seizure plates were prepared providing 16 animals per test concentration. Each plate consisted of five treatment concentrations and a control (water or solvent where appropriate).

Each trial consisted of an induction and subsequent recovery period. Firstly, 500µl of culture water was removed from each well and a 15-minute timer was set to denote the start of the induction period where 500µl of the assigned anaesthetic concentration (Table x) was added (from lowest to highest concentration). After the induction period an assessment of touch, heartrate and balance was performed for 30 minutes. Touch response was assessed by carefully approaching larvae with a pipette tip and using the following scoring system: *(3= no response, 2 = moves when touched and 1=moves when approached i.e. before touching).* Heartrate was assessed under a widefield microscope by counting for at least 10 seconds which was then used as an estimate of beats per minute (bpm).

After the 30-minute assessment period, 500µl of solution was removed from each well and replaced with the same volume of fresh culture water. This step was repeated, and before the addition of the second round of water a 15-minute timer was started to indicate the beginning of the recovery period. Once the timer had finished, the assessment was repeated. Larvae were disposed of in benzocaine (0.6M in 1% ethanol).

**Table S15. Test anaesthetic compounds and range of their concentrations used for assessing the maximum tolerated and effective anaesthetic concentrations in 4.5 dpf zebrafish larvae.**

| Compound | CAS No. | Exposure Concentrations (mM) | Exposure Concentrations (mg/L) |
|---|---|---|---|
| Benzocaine | 94-09-07 | 0.2, | 33.038, |
| | | 0.3, | 49.557, |
| | | 0.6, | 99.114, |
| | | 0.8, | 132.152, |
| | | 1.6, | 264.304, |
| | | 3.2, | 528.608, |
| | | 4.8, | 792.912, |
| | | 6.4, | 1057.216, |
| | | 12 | 2114.432 |
| Etomidate | 33125-97-2 | 0.004, | 0.977156, |
| | | 0.008, | 1.954312, |
| | | 0.016, | 3.908624, |
| | | 0.032, | 7.817248, |
| | | 0.063, | 15.39021, |
| | | 0.12, | 29.31468, |
| | | 0.2, | 48.8578, |
| | | 0.4, | 97.7156, |
| | | 0.6, | 146.5734, |
| | | 0.8, | 195.4312, |
| | | 0.9, | 219.8601 |
| | | 1.0, | 244.289, |
| | | 1.2, | 293.1468, |
| | | 1.4, | 342.0046, |
| | | 1.8, | 439.7202, |
| | | 2.2, | 537.4358, |
| | | 2.4 | 586.2936 |
| Isoeugenol | 97-54-1 | 0.1, | 16.4201, |
| | | 0.2, | 32.8402, |
| | | 0.4, | 65.6804, |
| | | 0.8, | 131.3608, |
| | | 1.6, | 262.7216, |
| | | 3.2 | 525.4432 |
| MS-222 | 886-86-2 | 0.1, | 26.129, |
| | | 0.5, | 130.645, |
| | | 1.0, | 261.29, |
| | | 5.0, | 1306.45, |
| | | 10, | 2612.9, |
| | | 20 | 5225.8 |
| 2-PE | 122-99-6 | 0.2, | 27.632, |
| | | 1.75, | 241.78, |
| | | 2.0, | 276.32, |
| | | 3.5, | 483.56, |

| | | | |
|---|---|---|---|
| | | 7.0, 14, 28, 31.5, 42, 56 | 967.12, 1934.24, 3868.48, 4352.04, 5802.72, 7736.96 |
| Quinaldine sulfate | 655-76-5 | 0.05, 0.1, 0.2, 0.4, 0.8, 1.6, 3.2, 4.8, 5.6, 6.4 | 12.0635, 24.127, 48.254, 96.508, 193.016, 386.032, 772.064, 1158.096, 1351.112, 1544.128 |
| Ethanol | 64-17-5 | 1, 5, 10, 15, 20% | |

**Table S16. Test anaesthtic compounds and their concentrations (in mM) used across all the assays presented in this study.**

| | 2-PE | Benzocaine | Etomidate | Isoeugenol | MS222 | Quinaldine sulfate |
|---|---|---|---|---|---|---|
| *In vivo* $Ca^{2+}$ functional brain imaging without stimulation | 2.000, 3.000, 5.000, 7.000 | 0.150, 0.300, 0.450, 0.600, 0.800 | 0.002, 0.004, 0.008 | 0.100, 0.200, 0.400 | 0.200, 0.300, 0.350, 0.400, 0.500, 0.700, 1.000 | 0.040, 0.050, 0.100, 0.200, 0.800 |
| *In vivo* $Ca^{2+}$ functional brain imaging with stimulation | 3.000, 5.000, 7.000, 28.000 | 0.450, 0.600, 0.800 | 0.004, 0.008, 0.016, 0.200, 0.600 | 0.200, 0.400, 1.600, 2.400 | 0.500, 0.700, 1.000 | 0.080, 0.200, 0.800 |
| Locomotion and light-dark phase activity | 2.000, 3.000, 4.000, 5.000, 6.000 | 0.025, 0.050, 0.100, 0.200, 0.300 | 0.00094, 0.00188, 0.00318, 0.00750, 0.01500 | 0.0125, 0.025, 0.050, 0.100, 0.200 | 0.025, 0.050, 0.100, 0.200, 0.300 | 0.040 0.080 0.200 0.400 0.800 |
| Olfaction-mediated avoidance behaviour | 2.000 | 0.200 | 0.016 | 0.100 | 0.100 | 0.100 |
| Cardiovascular assessment | 1.000, 2.000, 4.000, 7.000 | 0.300, 0.600, 1.000, 1.600 | 0.016, 0.032, 0.064, 0.128, 0.256 | 0.100, 0.200, 0.400, 0.800 | 0.200, 0.500, 1.000, 5.000 | 0.080, 0.800, 1.600, 3.200 |
| Determination of anaesthetic bioavailability | N/A | 1.600, 3.200, 6.400 | 0.350, 0.700, 1.400 | 0.200, 0.400, 0.800 | 0.500, 1.000, 2.000 | 0.200, 0.400, 0.800 |

**Determination of anaesthetic bioavailability to zebrafish**

Bioanalysis of all compounds except for 2-phenoxyethanol was performed on 4.5 dpf embryos in triplicates at three relevant concentrations. Four zebrafish embryos, in 300 µL of test solution were transferred to a 96 well MultiScreenHTS BV Filter Plate (Merck Millipore, Ireland). The test solution was removed under the vacuum and larvae were washed with culture water to eliminate residual test solution and subsequently transferred in 300 µL of pure water to a 96-well plate (Porvair Sciences, UK). 300µL HPLC- grade acetonitrile with internal standard was added and samples were homogenised for 3 min. to achieve extraction of analytes. LCMS grade water (900µL) was added to each well and after mixing the plate was centrifuged at 4000 rpm for 30 min. Supernatants from each well were transferred to a 96-well plate and removed for analysis. At the same time 300 µL of external solution was sampled and processed in the identical way to provide information of method’s analytical background.

Whenever the background was measurable the concentrations were subtracted from concertation of embryo samples.

It should be noted that we were able to determine the internal anaesthetic compound concentration and update for all agents tested except for 2-phenoxyethanol, which was found unsuitable for LC-MS analysis due to lack of formation of stable parent and fragment ions.

**LC-MS Analysis**

The analysis was on TSQ Vantage triple quadrupole mass spectrometer equipped with heated electrospray (HESI II) source coupled to Surveyor MS Pump Plus HPLC pump with HTC PAL autosampler (all Thermo Fisher Scientific, San Jose, CA).

Chromatographic separation was achieved using reversed-phase, 3 µm particle size, C18 Hypersil GOLD column 50 mm × 2.1 mm i.d. (Thermo Scientific, San Jose CA, USA). All the analytes were separated using a linear gradient of water and methanol, both containing 0.1% of ammonium hydroxide The initial conditions for the gradient consisted of 20% of methanol with ammonium hydroxide which was increased to 100% in 1.5 min and maintained for another 1.5 min before returning to the initial 20%. The flow rate was 500 µL/min. Retention times of analytes are presented in table Table S19.Temperature of autosampler was set at 6˚C, while the column was kept at a room temperature.

HESI probe was operating in both positive mode with an ion-spray voltage of 3.75 kV and negative with 2.7 kV. The heated capillary temperature was set at 275 °C and the vaporizer temperature was 350 °C. Nitrogen was employed as sheath and auxiliary gas at a pressure of 60 and 2 arbitrary units, respectively. The argon CID gas was used at a pressure of 1.5 mTorr and the optimum collision energy (CE) for each transition was automatically optimised by the software. Quantification of the analytes was performed using characteristic multiple reaction monitoring (MRM) transitions of precursor ion listed in Table S19.

Quantitative analysis was performed using an Internal Standard method.

**Table S17. Mass spectrometric and chromatographic parameters used in the analysis.**

| Compound | Ionisation Mode | Parent ion [m/z] | Product ion [m/z] | Collision energy [eV] | Retention time [min] |
|---|---|---|---|---|---|
| Benzocaine | + | 166.1 | 138.2 | 16 | 1.58 |
| Etomidate | + | 245.1 | 141.1 | 12 | 1.77 |
| Isoeugenol | - | 163.1 | 148.1 | 14 | 1.74 |
| MS222 | + | 166.1 | 138.2 | 14 | 1.58 |
| Quinaldine sulphate | + | 144.1 | 77.1 | 34 | 1.77 |

**Behavioural assessment of aversion**

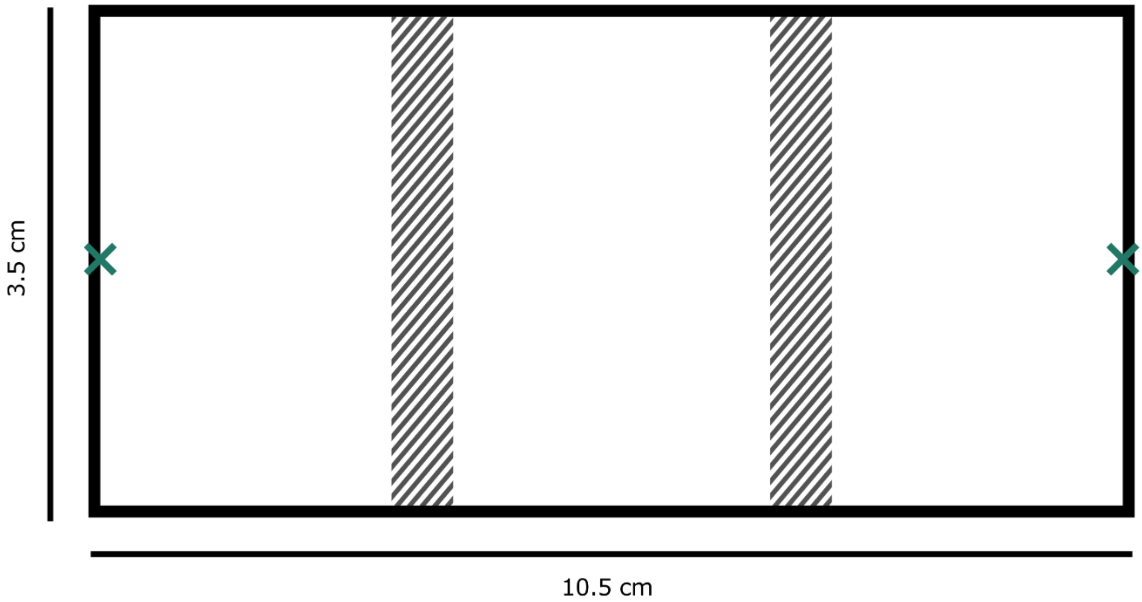


**Figure S4. Experimental setup for assessing olfaction-mediated aversion (top view).** The hatched rectangles denote the position of the removable barriers, and the X marks depict the sites of administration of the anaesthetic compounds and the water/solvent control.
